# *Toxoplasma* infection alters dopamine-sensitive behaviors and host gene expression patterns associated with neuropsychiatric disease

**DOI:** 10.1101/2021.08.16.456298

**Authors:** Graham L. Cromar, Jonathan Epp, Ana Popovic, Yusing Gu, Violet Ha, Brandon Walters, James St. Pierre, Xuejian Xiong, John Howland, Sheena Josselyn, John Parkinson, Paul W. Frankland

## Abstract

*Toxoplasma gondii* is a single celled parasite thought to infect 1 in 3 worldwide. During chronic infection, *T. gondii* can migrate to the brain where it promotes low-grade neuroinflammation with the capacity to induce changes in brain morphology and behavior. Consequently, infection with *T. gondii* has been linked with a number of neurocognitive disorders including schizophrenia (SZ), dementia, and Parkinson’s disease. Beyond neuroinflammation, infection with *T. gondii* can modulate the production of neurotransmitters, such as dopamine. To further dissect these pathways and examine the impact of altered dopaminergic sensitivity in *T. gondii*-infected mice on both behavior and gene expression, we developed a novel mouse model, based on stimulant-induced (cocaine) hyperactivity. Employing this model, we found that infection with *T. gondii* did not alter fear behavior but did impact motor activity and neuropsychiatric-related behaviurs. While both behaviors may help reduce predator avoidance, consistent with previous studies, the latter finding is reminiscent of neurocognitive disorders. Applying RNASeq to two relevant brain regions, striatum and hippocampus, we identified a broad upregulation of immune responses. However, we also noted significant associations with more meaningful neurologically relevant terms were masked due to the sheer number of terms incorporated in multiple testing correction. We therefore performed a more focused analysis using a curated set of neurologically relevant terms revealing significant associations across multiple pathways. We also found that *T. gondii* and cocaine treatments impacted the expression of similar functional pathways in the hippocampus and striatum although, as indicated by the low overlap among differentially expressed genes, largely via different proteins. Furthermore, while most differentially expressed genes reacted to a single condition and were mostly upregulated, we identified gene expression patterns indicating unexpected interactions between *T. gondii* infection and cocaine exposure. These include sets of genes which responded to cocaine exposure but not upon cocaine exposure in the context of *T. gondii* infection, suggestive of a neuroprotective effect advantageous to parasite persistence. Given its ability to uncover such complex relationships, we propose this novel model offers a new perspective to dissect the molecular pathways by which *T. gondii* infection contributes to neuropsychiatric disorders such as schizophrenia.

## INTRODUCTION

*Toxoplasma gondii* is an obligate intracellular protozoan of the phylum Apicomplexa and is thought to infect approximately one third of the global population^1^. While the definitive host of *T. gondii* are felines, *T. gondii* is thought to be capable of infecting any nucleated cell of any warm blooded animal (“intermediate hosts”). During acute infection, triggered by the hosts immune response, the fast growing multiplicative, tachyzoite, form of the parasite transforms into a slower growing, bradyzoite, form that forms cysts within tissues such as muscle and the brain, persisting throughout the life of the host^2^. While this chronic phase of infection is typically considered asymptomatic, its intimate association with the brain is thought to result in low-grade neuroinflammation with the capacity to induce changes in brain morphology and behavior^3^. For example, animal studies have shown that chronic *T. gondii* infection is associated with hyperactivity as well as reduced fear or increased risky behavior ^4–6^. While initial studies focused on mouse hosts overcoming their aversion to cat urine^5, 7^, a more recent study suggests that *T. gondii* infection is associated with a more general loss of predator avoidance behavior, an important distinction that may help facilitate non-sexual transmission between intermediate hosts^8^. Similar altered patterns of behavior have been observed in chimpanzees which are prey for leopards^9^ suggesting that altered behavior patterns in *T. gondii* infected humans (which includes among others, altered response to cat urine odor) have a common evolutionary basis^10–12^.

*T. gondii* has been linked with a number of neurocognitive disorders including dementia, schizophrenia (SZ), and Parkinson’s disease^13, 14^. For example, for SZ, which is thought to affect ∼1% of the population^15^, *T. gondii* is among the most common non-genetic risk factors, with meta-analyses of several large scale population studies reporting an adjusted odds ratio of 1.43^16, 17^. Interestingly, in AIDS patients, reactivation of chronic *T. gondii* infection can be accompanied by psychiatric disturbances such as delusions, auditory hallucinations and thought disorders in as many as 60% of patients although it is not clear these effects can be attributed to *T. gondii* alone^18^. Conversely, treating SZ individuals that are seropositive for *T. gondii* infection, with anti-parasitic drugs improves therapeutic response^19^. Nevertheless, little is known concerning the pathways by which *T. gondii* may elicit or worsen the symptoms of neurodegenerative diseases. Potential mechanisms include the modulation of neurotransmitters, such as dopamine^20, 21^, changes in hormone levels ^7, 22^, as well as neurophysiological changes induced by neuroinflammation^23^. Focusing on the former, chronic infection by *T. gondii* in a mouse model of Huntington’s has been shown to activate indoleamine-2,3-dioxygenase (IDO) which catalyzes the oxidation of tryptophan, a precursor of serotonin, to produce kynurenic acid (KYNA)^21^. In addition to increasing oxidative stress and apoptosis, KYNA can act as an antagonist of the α7 nicotinic acetylcholine receptor and may serve to modulate levels of GABA and glutamate with downstream consequences for glutamatergic, GABAergic and dopaminergic dysfunction^24^. Notably, KYNA levels are elevated in patients with SZ^25^. From a neurophysiological perspective, chronic infection with *T. gondii* results in sustained inflammation which through changes in cytokines, is likely to result in recruitment of immune cells and neurophysiological changes similar to those observed in schizophrenia and other disorders such as autism, and bipolar disorder^3, 8, 26^. For example, interferon-gamma (IFN-γ), interleukin-12b (IL-12b), and tumor necrosis factor (TNF) have all been positively correlated with cyst load and behavior^8^.

To better understand how *T. gondii* infection contributes to neuropsychiatric disorders, we propose a novel mouse model capable of providing more mechanistic insights into disease pathogenesis. Key to this model is the ability of cocaine to act via the dopamine pathway to induce SZ-like symptoms in mice^27^. Furthermore, it has been shown that cocaine activates striatal microglia, increasing production of TNF-α and thereby suppressing cocaine-induced sensitization^28^. Using this model allows us to compare the effects of *T. gondii* infected mice with cocaine-treated mice to investigate whether infected mice exhibit altered sensitivity to manipulations of dopamine using established behavioral paradigms (e.g., stimulant-induced hyperactivity). Using cocaine-treated mice, we examine both behavioral (stimulant sensitization, sensory motor gating and motor function) and functional consequences (via RNASeq) of *T. gondii* infection in mice, to identify pathways that may be disrupted in neurological disorders.

## RESULTS

### Chronic infection by T. gondii in mice does not alter fear behavior but does alter behaviors associated with neurological disorders

To examine the effects of chronic *T. gondii* infection on behaviors that are commonly impaired in neuropsychiatric disorders, we infected male mice with *T. gondii* and subsequently subjected them to a series of behavioral tests (**Figure 1A**; see Methods). As expected, histological examination revealed that cysts were present in the brains of mice infected with tachyzoites (**Figure 1B**). To further examine if such effects were strain specific, mice were infected with one of two different type III strains of *T. gondii:* CEP and VEG. Both strains were chosen due to their lower virulence in mice and ability to form brain cysts. In brief, adult mice (∼P60) were injected i.p. with ∼500,000 parasites. Eight weeks post-infection, mice were subjected to a series of tests (see Methods) investigating: 1) fear and anxiety behaviors (contextual fear conditioning, elevated plus maze (EPM) and open field); 2) motor ability (rotor rod); and 3) neuropsychiatric-related behaviors (cocaine sensitization and prepulse inhibition). For cocaine sensitization experiments, mice were injected with 20 mg/kg of cocaine daily for five days five minutes before testing. One week after the end of the sensitization the mice were tested drug free followed by a 20 mg/kg cocaine challenge 24 hours later. Prepulse inhibition (PPI) is a measure of sensory motor gating that evaluates ability to filter out irrelevant stimuli. In this case, this was assessed using the acoustic startle response. We used this task because it is impaired in (i) schizophrenia patients and (ii) rodent models of schizophrenia, and (iii) impairments in rodent models are reversed with antipsychotic drugs. In an additional cohort of mice, cocaine or saline was administered for 5 days prior to sacrifice, and striatum and hippocampus samples processed for histology and RNA sequencing. Since both strains of *T. gondii* (CEP and VEG) produced similar results (**Supplementary Figure 1**), here we report the results only for strain CEP.

**Figure 1:**
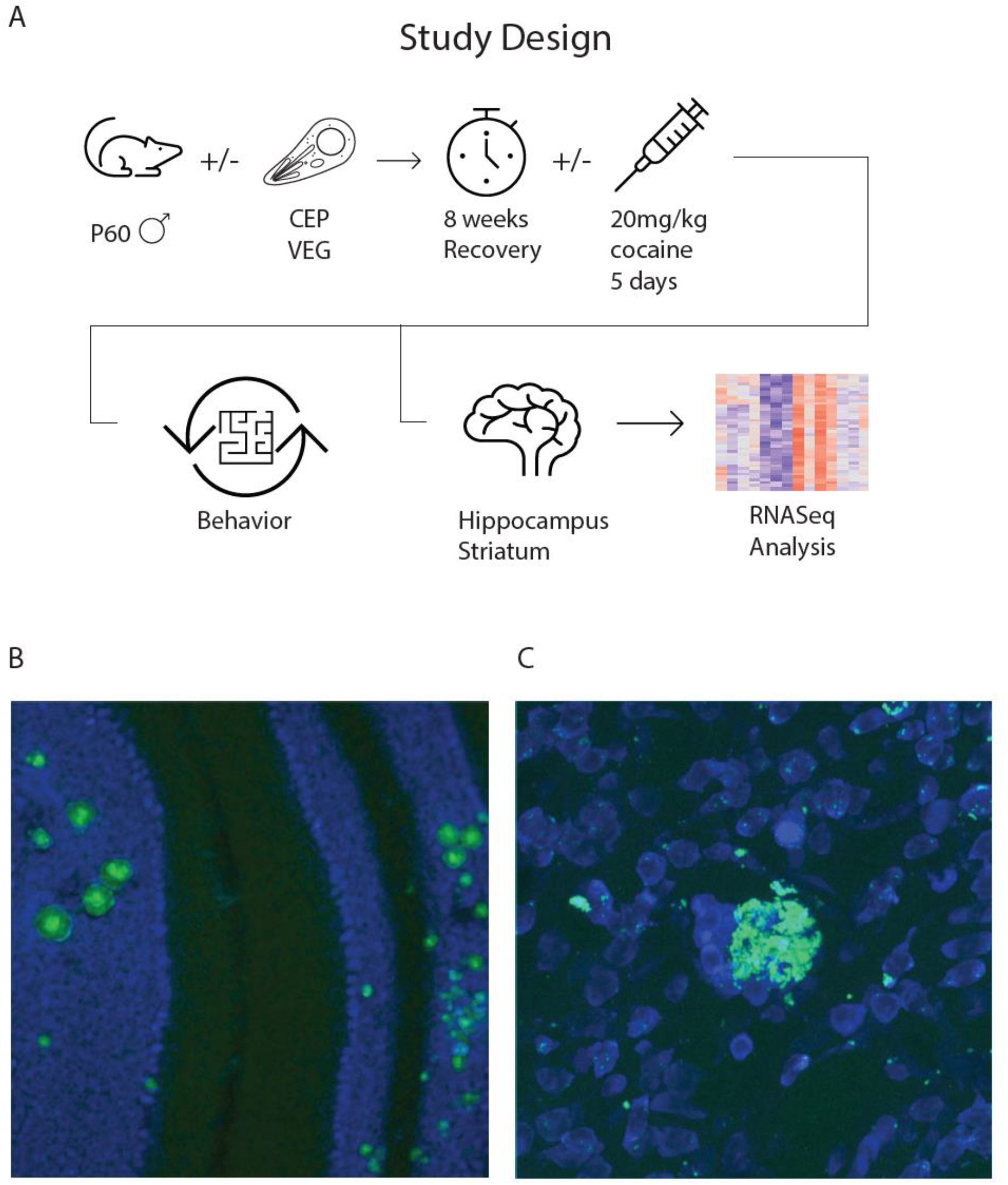
Study design and histological images of *T. gondii* infection in the mouse brain. A. Schematic overview of study design. B-C. Images showing sections of mouse brain exhibiting wide-spread infection of GFP-expressing *T. gondii* (strain CEP; green Propidium iodide counterstain; blue; larger cysts and smaller inclusions can be seen throughout the brain including in the cerebellum (B) and striatium (C).

Focusing on behaviors, we found that mice infected with *T. gondii* acquired contextual fear conditioning normally. There were no differences in overall freezing levels (Repeated measures ANOVA effect of group *F*(1,37) = 0.048, *p = 0.83*; **Figure 2A**) or freezing levels over the course of training (Minute by group: F(4,148) = 0.076, *p = 0.99;* **Figure 2B**), indicating that shock reactivity is unaltered in infected mice. *T. gondii* infection also did not alter the subsequent retention of the contextual fear memory (independent samples t-test, t(39) = 0.91, *p =* 0.37) or anxiety-associated behaviors. Infected mice did not show altered anxiety-related behavior in the elevated plus maze (**Figure 2C**). They did not spend significantly more time than control mice in the open arms (Two-way ANOVA, Zone x Group: F(1,37) = 0.067, *p =* 0.80). Further, we observed no significant group differences in the time spent in each zone of the open field test (Two-way ANOVA, Group × Zone: F(2,74) = 1.67, *p = 0.20;* **Figure 2D**). These results are in contrast with a recent study of B6CBAF1/J mice infected with the type II *T. gondii* strain, ME49, which reported that infected mice spent significantly more time in the open arms of the EPM and less time in the open field, the latter apparently conflicting result explained by an increase in exploratory behavior^8^.

**Figure 2:**
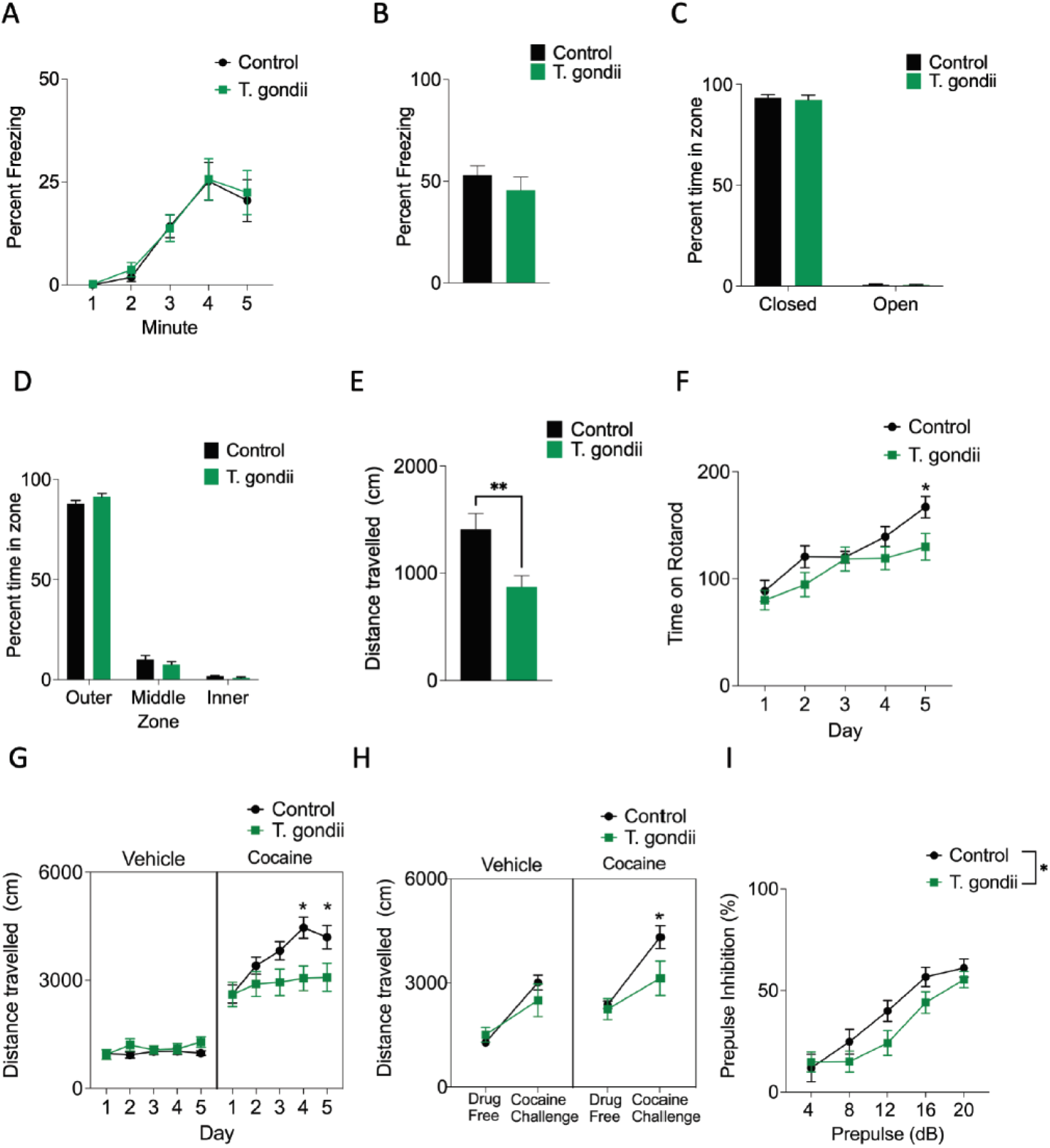
Behavioral characterization of *T. gondii* (CEP) injected mice. A-B. Infected mice exhibited normal acquisition (A) and expression (B) of contextual fear memory. C-D. Anxiety related behaviors including time spent in the open arm of the elevated plus maze (C) or inner zone of the open field (D) did not differ between infected and control mice. E. Infected mice travelled significantly less distance in the open field compared to controls. F. On the rotarod, *T. gondii* infected mice exhibited impaired acquisition relative to control mice indicative of potential motor impairments. G-H. In the behavioral sensitization paradigm, *T. gondii* infected mice showed a blunted sensitization to the effects of cocaine, as exhibited by a significantly decrease in distance travelled following successive administrations of cocaine (G). One week following sensitization *T. gondii* infected mice still exhibited a blunted response to a challenge dose of cocaine compared to control mice. I. *T. gondii* infected mice exhibited blunted prepulse inhibition compared to control mice.

In terms of motor activity, we found that infection with *T. gondii* significantly reduced overall locomotor activity in the open field test (t(37) – 2.99, *p =* 0.0049; **Figure 2E**). Additionally, *T. gondii* infection slowed rotarod acquisition (Repeated measures ANOVA, significant session x group interaction, F(4,37) = 4.00, *p =* 0.0041). Specifically, control mice outperformed *T. gondii* infected mice on the final day of training (*p = 0.033)* (**Figure 2F**). These results are consistent with previous studies that have shown *T. gondii* infection results in sensorimotor deficits in mice without major brain damage or cognitive dysfunction^29^.

To assess whether dopaminergic sensitivity is altered in *T. gondii* infected mice, we examined behavioral sensitization to the hyper-locomotion inducing effects of cocaine (**Figure 2 G-H**). There was an effect of *T. gondii* infection (Three-way ANOVA, significant group by drug by session effect: F(4,120) = 4.13, *p =* 0.0036) We found that *T. gondii* infected mice show an equivalent initial locomotor response to cocaine (day 1, *p = 0.97*) but show a significantly blunted sensitization upon repeated cocaine administration (Post-hoc Newman-Keuls test; Days 4 and 5, *p ≤ 0.0011)*. We also observed a lasting effect of *T. gondii* infection on locomotor activity when mice were tested one week following sensitization (Two-way ANOVA main effect of group F3,60) = 9.39, *p = 0.000035).* Control and infected mice that had previously been administered saline show comparable locomotor activity (Post-hoc Newman-Keuls test *p= 0.63)* when tested one week later. However, *T. gondii* infected mice that had previously been exposed to repeated cocaine administration again showed a blunted response compared to control mice challenged with cocaine (Post-hoc Newman-Keuls test, *p = 0.030)*. However, their locomotor response did not differ significantly from mice that had not previously received cocaine (Post-hoc Newman-Keuls test, *p = 0.061)*.

We also examined paired-pulse inhibition as it is a behavior that is commonly disrupted in neuropsychiatric conditions, as well as in their animal models. Here we observed that *T. gondii* infected mice showed an impairment in their prepulse inhibition response compared to control mice (**Figure 2I**). Levels of PPI increased with increasing prepulse intensity (Two-way ANOVA, main effect of prepulse intensity F(4,110) = 6.69, *p = 0.000074*). However, this increase was blunted in infected mice (Two-way ANOVA, main effect of group F(1,110) = 4.79, *p = 0.031*

While we did not observe altered fear or anxiety related behaviors which, have been described previously as a result of *T. gondii* infection we did observe alterations in motor performance and interestingly in behaviors that are commonly associated with neuropsychiatric-conditions. These observations are significant given the previously described link between *T. gondii* infection and risk of neurocognitive disorders such as schizophrenia. Notably, these effects were replicated for two different strains of the parasite (**Supplementary Figure 1**). In the following sections we used RNASeq to investigate how changes in gene expression, associated with chronic infection by *T. gondii*, may contribute to these changes in behavior. Since we observed no significant behavioral differences between strains VEG and CEP, only a single strain (CEP) was used for expression profiling.

### Exposure to cocaine and chronic infection by T. gondii result in distinct, tissue-specific, and treatment-specific gene expression profiles

To examine the impact of cocaine and *T. gondii* infection on gene expression in the mouse brain we performed RNASeq on the striatum and hippocampus from samples collected from all four treatment groups (+/- cocaine and +/- *T. gondii*). Both tissues are established sites of *T. gondii* infection; striatum was selected for its role in motor functions and response to rewarding and aversive stimuli, while the hippocampus is involved in memory and learning. Four biological replicates were performed. The resulting ∼50M reads per sample were filtered for quality and mapped to the mouse genome using STAR^30^, resulting in a matrix of read counts for each gene/sample. Mouse gene counts were normalized and analyzed using DESeq2 (see Methods – Statistical Analysis). Out of a possible 8,563 *T. gondii* genes only 351 were detected and only 3 with greater than 5 reads, reflecting the relative abundance of mouse tissue and lower activity associated with the bradzyzoite stage of the parasite^31^. In our analysis of host gene expression, we applied an ANOVA-like model to identify genes that exhibit differential expression upon each treatment individually or in combination, including any interactions between treatments (see Methods). Overall, we identified 4439 genes that were differentially expressed (DE) relative to controls (**Supplementary Files 1-3**). These include 2912 genes associated with the hippocampus samples and 3077 genes associated with the striatum samples. 1550 of these genes were common to both tissues.

The majority of DE genes (2344 and 2739 for hippocampus and striatum respectively) were associated with a single treatment (either *T. gondii* infection or cocaine), and these treatments resulted in more genes being upregulated than downregulated (**Figure 3**). Furthermore, across most of the single-treatment outcomes (i.e., up or downregulated upon treatment with cocaine or *T. gondii*), the number of DE genes common to both tissues (3.0-15.3%) was significantly smaller than expected (25.9%; Fisher’s Exact Test p = 3.73e-12 to p = 2.20e-16; **Supplementary File 4**). DE genes exhibiting upregulation in response to *T. gondii* were an exception, showing greater overlap than expected (36.2% overlap versus an expected 25.9%, Fisher’s Exact Test p = 2.20e-16) and comprising the largest gene set for both tissues (hippocampus=1150; striatum=1555). Notably, a larger number of genes responded to the cocaine treatment in the hippocampus (Fisher’s Exact Test p = 7.35e-15), while more genes responded to *T. gondii* in the striatum (Fisher’s Exact Test p = 2.05e-14), likely reflecting the different targets of the two treatments. Among the genes responding to both treatments, the largest overlap between the two tissues were 37 genes that were significantly upregulated in response to *T. gondii* infection, but downregulated in response to cocaine, suggesting differing effects on the same targets. In the next section we attempt to identify the functional consequences associated with these changes in gene expression.

**Figure 3:**
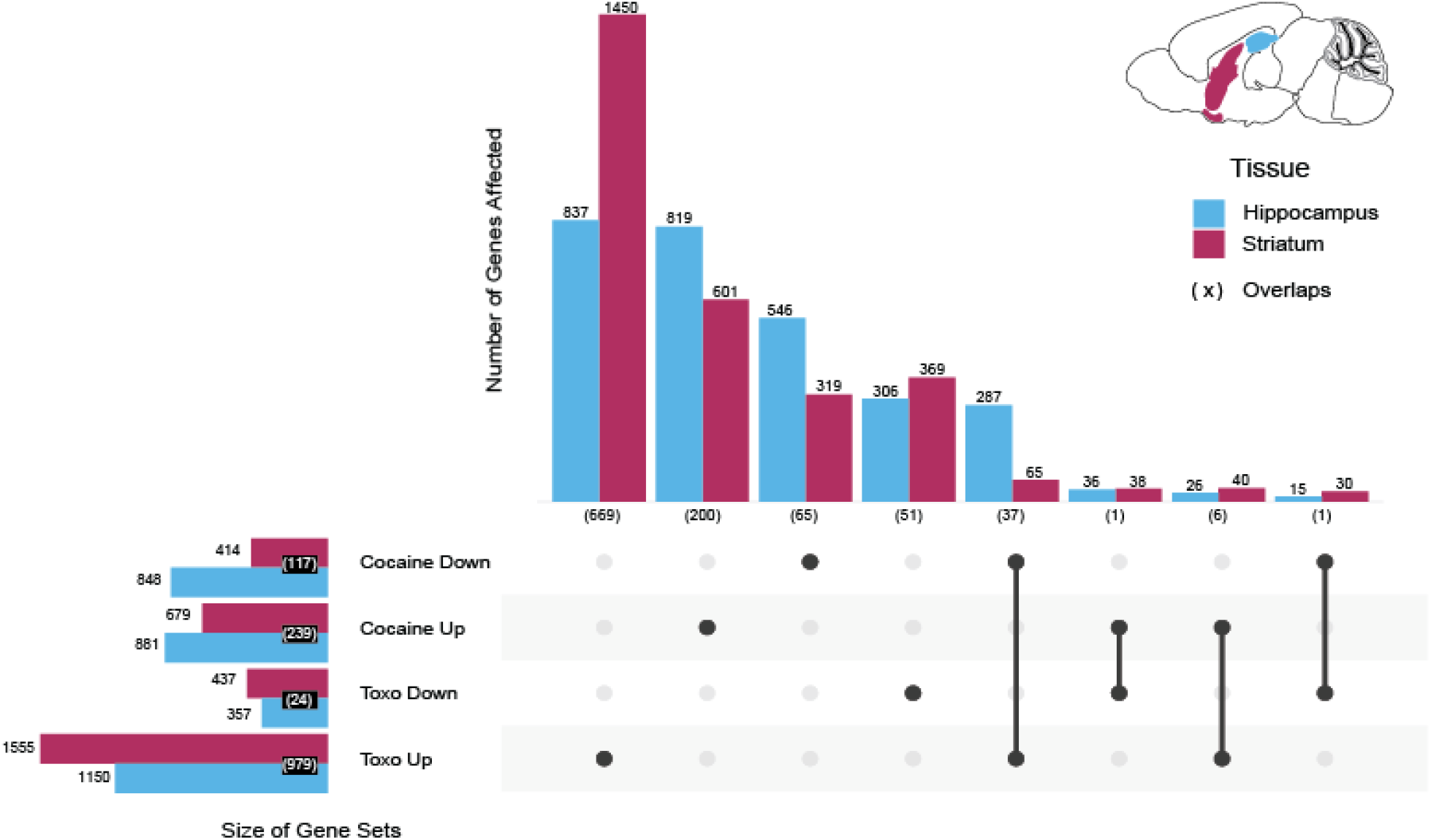
**Expression of differentially expressed genes responding to cocaine and *T. gondii* infection**. Shown are numbers of genes by response (size of Gene Sets) with overlaps by treatment category and tissue for both single and combination responses (vertical bars). Within categories, tissue overlaps are shown (bracketed). Genes represented are mouse genes identified as being significantly differentially expressed (likelihood ratio test, FDR<0.1) across pairs of conditions within a tissue.

### DE genes uncover similar high-level functional pathways in striatum and hippocampus

To provide an overview of functions associated with DE genes, we first investigated pathway representation as defined by the Reactome Pathway Database^32^ (**Supplementary Figure 2A**). Of the 2912 and 3077 genes differentially expressed in the hippocampus and striatum respectively, 1469 and 1439 were annotated with one of the 26 Reactome pathway categories. Of these, there was a significant enrichment of genes annotated with “Immune System” (453 - 30.8% and 513 - 35.6% in the hippocampus and striatum respectively; Fisher’s exact test p = 2.2e-16,). While this enrichment is consistent with the previous study that investigated the type II strain, ME49^8^, we found comparatively few genes annotated with ”Neuronal System” (61 hippocampus, 22 striatum) although we note that the previous study also reported lower p-values for neuronal-related categories relative to immune-related categories. This may reflect limitations in annotation coverage with the Reactome database; we found for example many DE genes expressing immunoglobulin chains which were unannotated by Reactome (83/107 hippocampus, 103/140 striatum). Nevertheless, both tissues were associated with similar numbers of DE genes for 22 of the 26 top level pathways (z-score of mean difference < 1.0; **Supplementary Figure 2B**). The greatest disparity was observed with genes annotated to “Gene Expression” (z-score = 3.427) perhaps reflecting a greater recruitment of genes responding to *T. gondii* infection in the striatum. Genes annotated to more than one Reactome pathway reflect related functions, with the largest overlaps involving “Immune System” (**Supplementary Figure 3**). To further dissect functional enrichments, given the limitations in annotation coverage we next focused on the use of Gene Ontology (GO) terms^33, 34^.

### Gene set enrichment identifies a wealth of neurologically relevant GO biological processes statistically enriched in DE genes

To identify more specific pathways and processes impacted by exposure of the mouse brain to *T. gondii* and/or cocaine we performed gene set enrichment analysis (GSEA) using the Cytoscape plug-in, BiNGO^35^. This initial analysis confirmed our high-level observation of a robust, general immune response which is a natural consequence of *T. gondii* infection (**Supplementary File 5**). However, the large number of enriched terms (2755 and 2014 for hippocampus and striatum respectively) render it challenging to identify consistent functional themes. Additional analyses focusing on DE subsets, representing each of the eight categories of response (i.e., up or downregulated upon exposure to cocaine and/or infection with *T. gondii*) were further unable to identify functional trends among enriched terms (data not shown). Similar observations were obtained using the largely independent Reactome and KEGG pathway data sets (see Methods and **Supplementary File 6**).

Given our interest in the neurological significance of DE genes and how these might vary across the treatments and tissues, we therefore developed a literature-guided, systematic approach to filter neurological-related terms from the larger sets of 2000+ terms identified as significantly enriched (see Methods). This process allowed us to collapse 734 GO terms into 34 distinct higher-level ‘term groups’ (**Supplementary File 7**). Each of these 34 term groups were then assigned to a treatment/tissue if at least one GO term associated with that group was enriched in that treatment/tissue (**Figures 4A** and **B**). Interestingly, genes down regulated in response to *T. gondii* infection were less likely to be associated with neurologically relevant annotations, the exception being those genes downregulated in the hippocampus due to infection by *T. gondii* but unaffected by cocaine exposure. In contrast, cocaine exposure led to many genes involved in multiple neurological term groups to be either up or down regulated.

**Figure 4:**
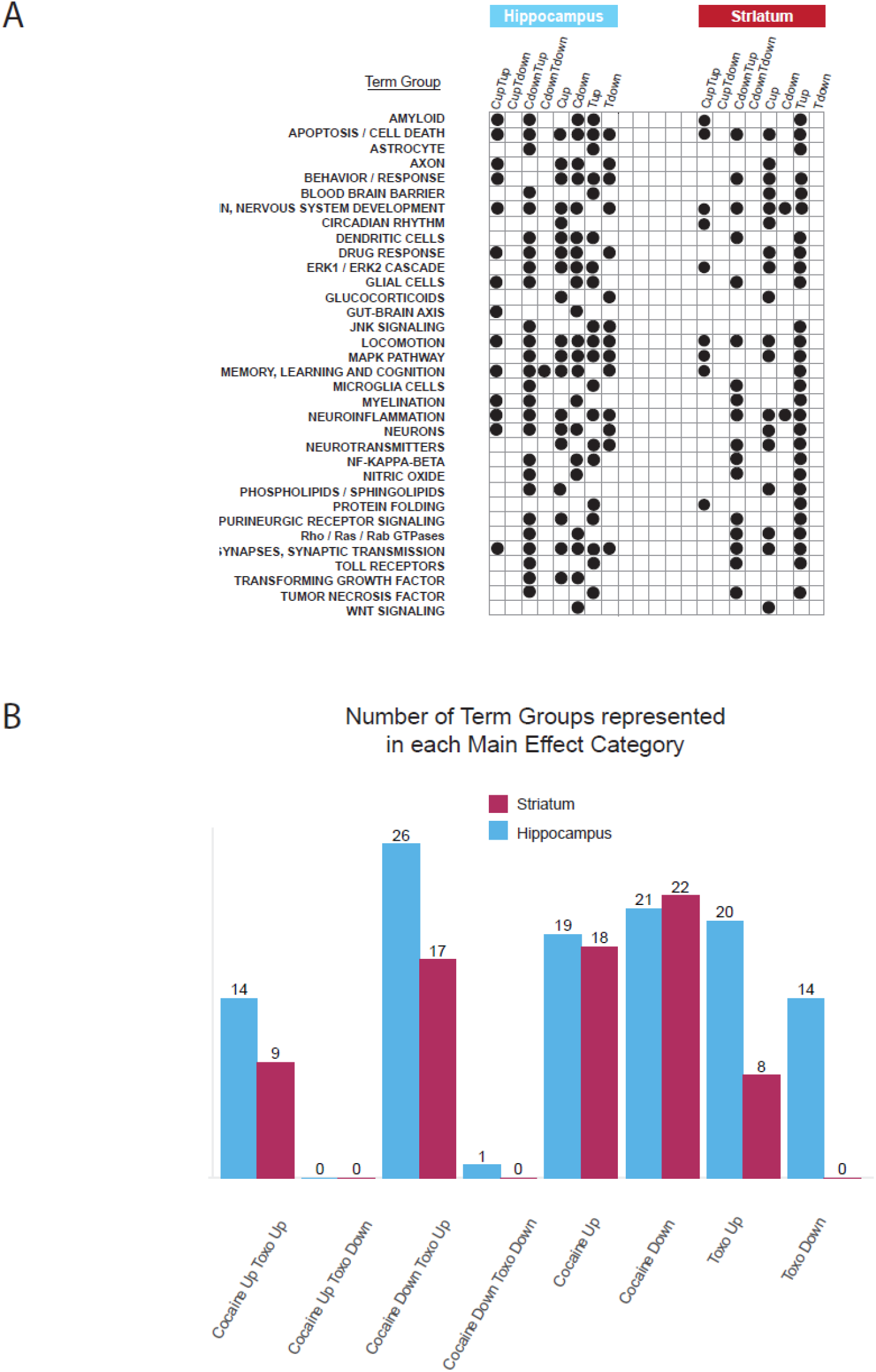
Summary of neurologically relevant, enriched annotations. A - Occurrence of neurologically relevant term groups across main effect and interacting effect categories in hippocampus and striatum. B - Number of term groups represented in each main effect category in each tissue.

Among the genes upregulated by *T. gondii* were many involved in neuroinflammation processes that are typically only expressed during infection. Neuroinflammation has been linked to several neuropsychiatric disorders (e.g., Parkinson’s disease) which also correlate with *T. gondii* infection^13^ and it has been suggested that this may be driven through the activation of microglia and astrocytes in response to latent toxoplasmosis^36, 37^. Among the DE genes we identified, 71 were associated with the genesis, growth, activation, or regulation of glial cells, including 19 associated with microglia and 15 associated with astrocytes (**Supplementary File 7**). TNF-alpha, an inflammatory cytokine released by activated microglia, was found to be upregulated in response to *T. gondii* infection. This is particularly interesting as TNF-alpha can result in the internalization of synaptic AMPA receptors on striatal neurons, depressing cocaine-induced behavioral sensitization^28^. Normally, repeated doses of cocaine are needed to reduce sensitization. However, the persistent activation of microglial by *T. gondii*, would explain the observed lack of sensitization in these mice upon cocaine challenge. A further 64 DE genes upregulated by *T. gondii* were associated with the growth and proliferation of dendritic cells which contribute to various forms of neuroinflammation associated with neurodegenerative autoimmune disease, injury and CNS infections^38^. Among genes that are upregulated in response to *T. gondii* and cocaine we identified 247 genes associated with mitogen-activated protein kinases (MAPKs) as well as 105 DE genes annotated to NFκB signaling, regulation and activation. MAPKs are crucial regulators in the production of inflammatory mediators^39^ whereas NFκB is one of the most important transcription factors for many proinflammatory genes.

Potentially driving these responses, the observed gene expression patterns show evident signs of cellular stress. 587 DE genes were associated with apoptosis/cell death of neurologically relevant targets including 32 associated with positive regulation of neuron death which are over-expressed in response to *T. gondii* and/or cocaine. Additionally, 52 DE genes were associated with protein folding, 23 of which relate to the GO term ‘response to topologically incorrect protein’ which are overexpressed in striatum in response to *T. gondii* alone or in combination with cocaine. Finally, 42 DE genes were found to be associated with Toll-like receptors (TLRs) and their regulation. TLRs recognize endogenous molecules liberated from damaged tissues, thereby modulating host inflammatory responses to CNS parasitic infections including toxoplasmosis. TLRs have been implicated in neurogenesis and learning and memory even in the absence of any underlying infectious etiology^40^.

In addition to eliciting responses related to neuroinflammation, we also identified 73 DE genes responding to cocaine and/or *T. gondii* associated with terms relating to the metabolism, transport, secretion and regulation of various neurotransmitters and their receptors. Further, 185 DE genes relate to synapses and synaptic transmission. DE genes associated with signal transduction mediated by small GTPases (120 DE genes) likely play roles in both neuroinflammation and neurotransmission. RAP1, for example, is activated in response to dopamine via activation of PKA which itself activates over 100 additional substrates regulating neuronal excitability and behavior^41^. Here we find that *RAP1A* is significantly under-expressed in response to cocaine in striatum perhaps indicating there are compensatory mechanisms for increased activity. On the other hand, *RAP1B* is over-expressed in response to *T. gondii* in both tissues suggesting *T. gondii*, independent of cocaine, has effects on this important behavioral axis potentially resulting in long term changes in gene expression downstream of RAP1B. Calcineurin, which in addition activates T cells, has been linked to receptors for several brain chemicals including dopamine, glutamate and GABA^42^. Calcineurin deficient mice have been reported to exhibit symptoms similar to humans with schizophrenia including working memory impairment, attention deficits and, impaired social behavior^43^ and, a reduction in calcineurin expression in the hippocampus of (human) SZ patients has been reported^44, 45^. Interestingly, we observed significant down-regulation of two positive regulators of calcineurin signaling (LMCD1 and SLC9A1) in response to cocaine and *T. gondii* in hippocampus. Given previous associations between *T. gondii* infection and KYNA production^21^, and its link to SZ^37^, we were intrigued to find that in response to *T. gondii* infection kynureninase (KYNU) [EC:3.7.1.3] was significantly upregulated in both tissues, while the adjacent enzyme in the pathway, 3-hydroxyanthranilate 3,4-dioxygenase (HAAO) [EC:1.13.11.6] was also upregulated in the striatum; cocaine exposure had no impact.

Despite the large number of GO-terms identified as enriched in DE genes, by collapsing multiple neurological-related terms into 34 term groups, we were able to identify a wealth of genes implicated in modulating the host brains response to infection by *T. gondii.* In the next section we further show how these analyses can be further enriched through developing a customized ontology.

### Recovery of neurologically relevant annotations is aided by a novel, context enrichment approach

Given the previously mentioned associations between *T. gondii* infection and neurological risk, albeit consistent with previous work^8^, it was surprising to find neurologically relevant terms ranked among the least significantly enriched GO annotations. As previously noted, due to the need to perform multiple testing correction, the breadth of GO annotations can reduce the power to detect significant association and hence mask the identification of biologically meaningful enrichments^35, 46, 47^. Here, we propose that DE genes identified in this study, represent two distinct subsets, one associated with a robust, highly enriched, general immune response and the other representing a smaller subset, associated with neurological effects. A successful separation of these would reduce the breadth of annotations, increasing the statistical power to detect significant associations with neurologically relevant annotations.

To test this, we first defined a set of root keywords (“neuro”, “brain”, “behavior”, “locomot”, “memory”, and “synap”) which we considered, based on our curation of the literature, to be both relevant to the context of our study and non-immune system specific. An AmiGO^48^ search for GO biological process terms associated with these keywords was then performed and resulted in 1632 GO terms which we deemed collectively to define the functional “context”. The terms retrieved by each keyword (**Supplementary File 8**) were largely complimentary as shown by the relatively small number of overlapping terms (**Supplementary Figure 5**). Using the collated full-text descriptions of these GO terms, an analysis of word frequencies confirms that they indeed represent a full range of neurologically relevant concepts (**Figure 5A**). These terms were next used to filter the DE gene set, removing genes that were not associated with at least one annotation matching the defined context, a process we refer to as “context enrichment”. The intent was to distill, from among 4439 DE genes identified across both tissues, a neurologically related DE subset distinct from those genes relating to the expected, general immune response.

**Figure 5:**
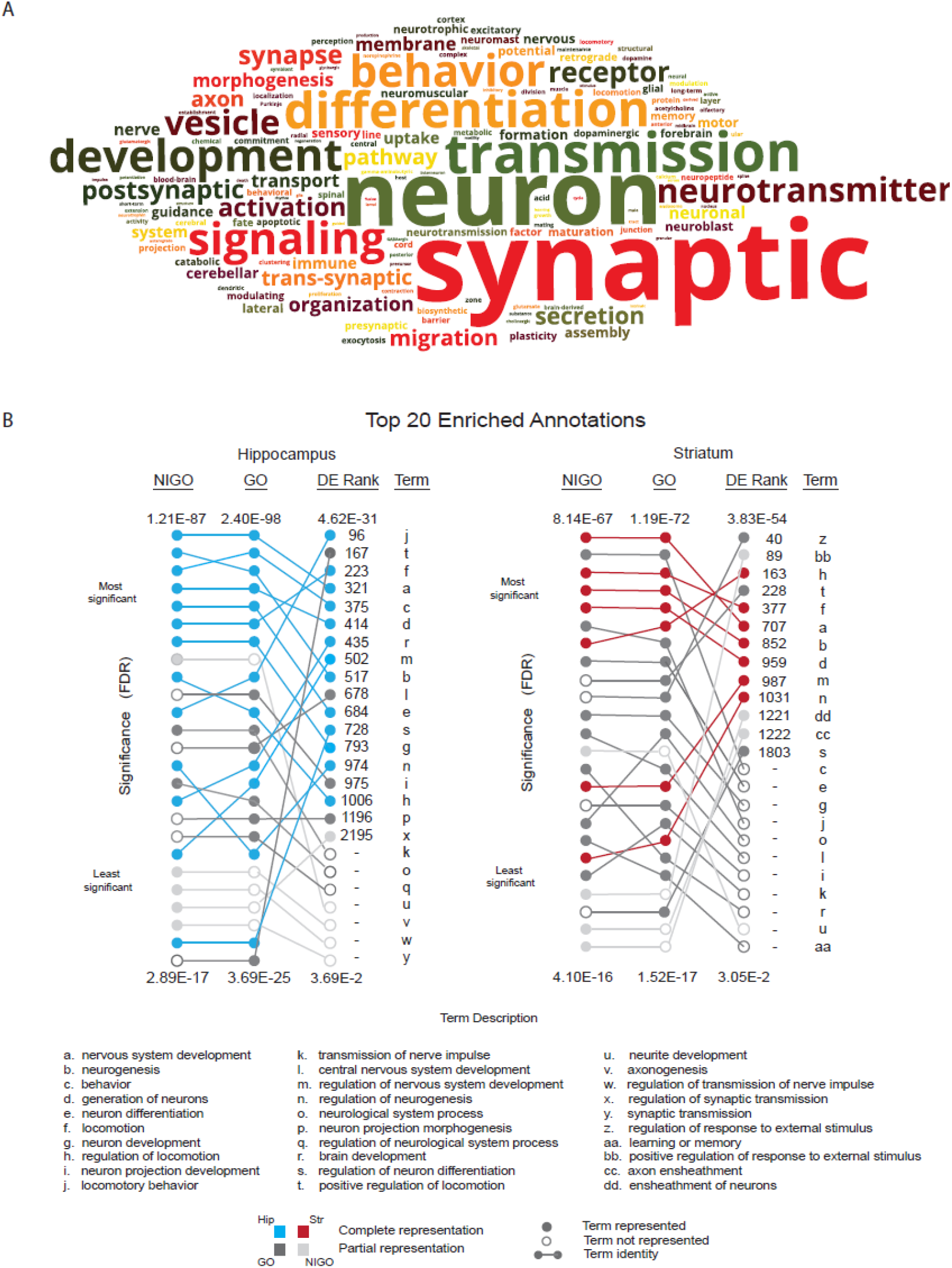
Context enrichment of DE genes. A - Frequency of words in full-text descriptions of GO terms defining the context filter. These GO terms result from an AmiGO search based on six key words defining a neurological ‘context’. B - For the context filtered subset of DE genes, a comparison of top 20 enriched annotations (GO vs. NIGO) in hippocampus and striatum and their relative rankings in a GO enrichment analysis of the full DE gene set.

GSEA was performed on the resulting 641 context enriched genes (466/2912 hippocampus and 450/3077 striatum genes, with an overlap of 275 genes) and results were visualized for each tissue as a Cytoscape network (**Supplementary File 10**). These graphs appear to capture many of the processes identified in our earlier curation approach and again highlight the diverse range of biological processes potentially contributing to neurological effects. Comparisons of the top 20 enriched GO annotations in the context enriched set reveal considerably lower rankings in the full GO set of terms, accompanied with lower corrected p-values (**Figure 5B**). For example, the top ranked term in the GSEA results for hippocampus context enriched genes, “Locomotory Behavior” was the 321^st^ term result for DE genes. Further, the relative order of these top twenty terms was also scrambled between gene sets while among the top 20 terms, 7/20 in hippocampus and 11/20 in striatum were not detected as significant in the larger GO gene set. Indeed, of the 641 context enriched genes, 114 were missed in the larger analysis illustrating how genes and terms associated with immune responses masked our ability to detect a greater depth of neurologically relevant signals. By way of example, we discovered a modest but significant overlap (Fisher’s Exact Test p = 0.0492) between our context enriched gene set and genes highlighted in a recent meta-analysis of differentially methylated positions associated with psychosis, schizophrenia and treatment-resistant schizophrenia^49^ which was not detectable using the full DE gene set.

To further validate our context enrichment approach, we performed GSEA using NIGO, a GO-derived subset focusing on neuro-immune function terms^47^. Comparing the top 20 term results obtained using the NIGO dataset with our enriched dataset revealed a high degree of correspondence in ranking of terms, confirming our ability to capture relevant terms. While NIGO might appear to represent a suitable solution to the enrichment analysis we note that over the past decade since NIGO was published, considerable efforts have been made to address deficiencies in GO with respect to both neurologically relevant gene annotations and the ontology itself^50–52^. Thus, we expect many terms to be present among current DE gene annotations that are not reflected in NIGO and therefore excluded from the analysis. This likely accounts for observed discrepancies in term detection and ranks between NIGO and GO for our AmiGO-based set. Updating NIGO would require a systematic clipping of the Gene Ontology and was therefore outside the scope of our study. Nevertheless, the simpler approach developed here proved capable of overcoming current limitations of GO annotations by revealing additional neurologically relevant terms and could be readily applied more widely to focus on other specialized gene sets.

### Gene subsets present in both striatum and hippocampus exhibit complex responses to the two treatments implying they operate in parallel or intersecting pathways

Given the ANOVA-like model used to identify DE genes, we were also able to define subtypes of genes exhibiting statistically significant “interactions”, in which the combined effect of cocaine exposure and *T. gondii* infection elicited a change in expression that was not simply the additive effect of each treatment in isolation. In total we identified 40 hippocampus and 165 striatum DE genes, that could be organized into 8 ‘functional effect’ groups based on their gene expression patterns (**Supplementary File 9**). While we note that the hippocampus had fewer genes with interactions than striatum, direct comparisons between the two tissues should be cautioned as these tissues were processed and normalized independently. To set the stage for future follow up investigations we highlight several examples of antagonistic or synergistic interactions suggestive of competitive or complementary pathways.

For both tissues, the most common type of interaction involved genes that exhibited downregulation in response to both cocaine and *T. gondii* by themselves, but had no effect in combination (i.e., rescue of the *T. gondii* and cocaine effects; subgroup ‘b’, **Figure 6A**). This pattern was enriched relative to other patterns (p < 0.01 for both tissues; Chi-square Goodness of Fit Test under the null hypothesis that all categories are equally represented). GSEA of KEGG pathways for this subgroup suggested that cocaine may compete with or otherwise impede *T. gondii* in its attempt to down regulate expression of host proteins critical to the fidelity of cellular protein production at a post-transcriptional and/or post-translational level. We found that 27/31 annotated striatum proteins in this set were associated to the KEGG pathway “Protein families: genetic information processing” and, significant KEGG Pathway enrichments included “spliceosome” (Hypergeometric test p = 4.6596E-6) and “chaperones and folding catalysts” (Hypergeometric test p = 1.5494E-4; **Figure 6B, Supplementary File 5**). Also included in this subgroup are two genes (*ATF2* and *ARNTL*) annotated to “Dopaminergic synapse”. Altered *ATF2* expression has been associated with progressive reduction in postsynaptic dopamine response^53^ whereas ATF2 with ARNTL (and CLOCK) comprise a heterodimeric enhancer of circadian clock genes (*PER1*, *PER2*, *PER3*). The latter are indeed differentially expressed, although they are not within this subgroup. Notably, the dysregulation of circadian rhythms has been associated with neurological disorders involving dopamine including Parkinson’s Disease^54^.

**Figure 6:**
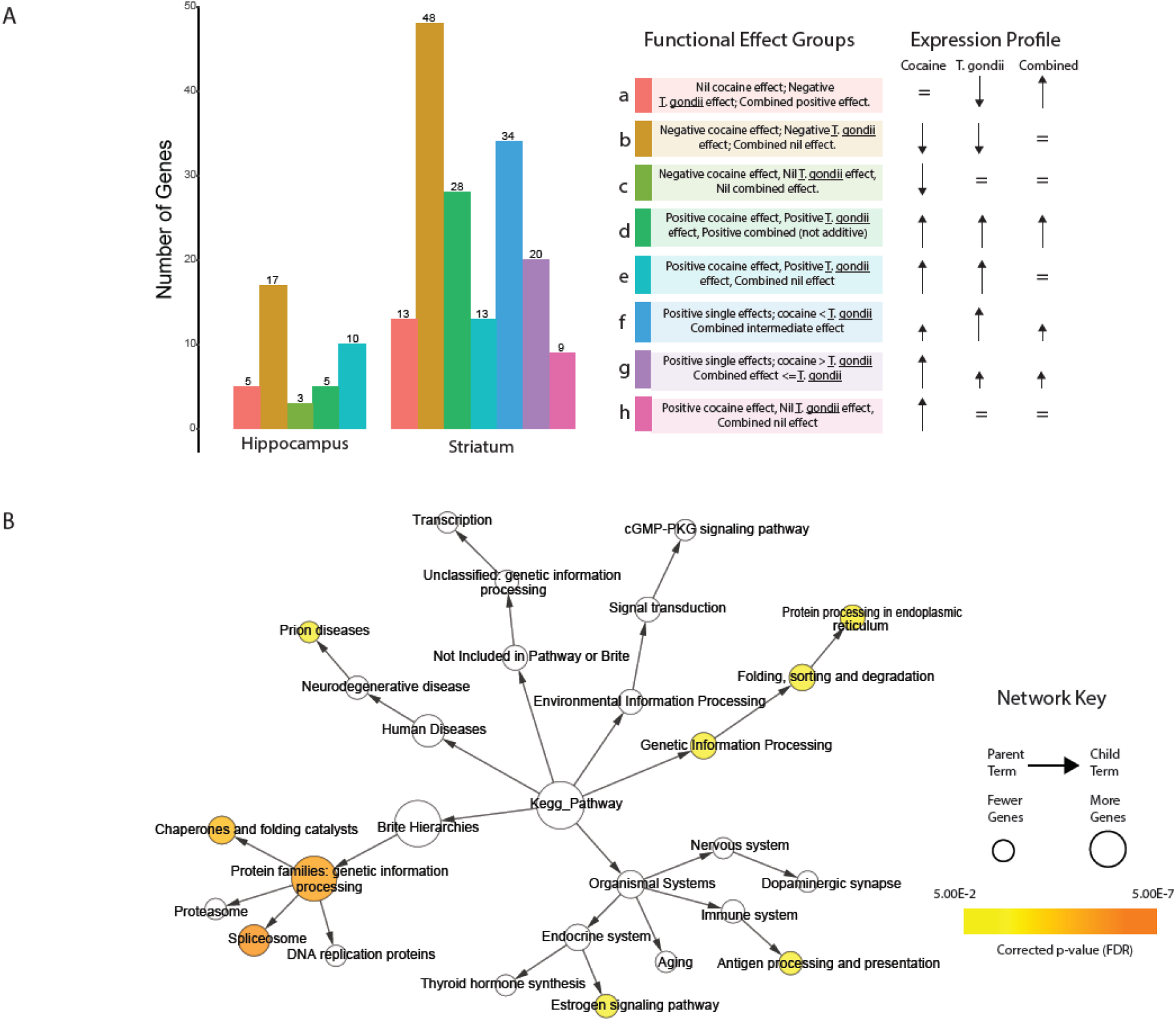
Functional characterization of mouse genes with interacting effects. A - Frequency of mouse genes with statistically significant interactions by functional effect category (hippocampus and striatum). B – Network visualization of KEGG pathway enrichments for functional effect group b in striatum (largest category). For context, all significantly enriched terms before correction for multiple testing are shown. Colors represent significant terms after multiple testing correction (hypergeometric test, FDR < 0.05) as implemented in the Cytoscape plugin BiNGO^35^.

Focusing on functional effects in the hippocampus, we identified a unique subgroup (subgroup ‘c’, **Figure 6A**) comprising 3 DE genes whose expression was downregulated by cocaine, with no response to *T. gondii* or the combined condition (i.e., rescue of the cocaine effect). These were *VGLL4*, encoding a transcriptional co-factor^55^; *SOX21*, encoding a transcriptional activator^56^; and *PSTPIP2*, whose product regulates F-Actin bundling and enhances motility in macrophages^57^. *SOX21* is associated with the GO terms “positive regulation of myelination”, “axon ensheathment in central nervous system”, “central nervous system myelination”, and “negative regulation of Schwann cell proliferation”. From a neurological perspective, it is interesting to note that a gene or genes that may cause dis-regulation of myelination in response to cocaine sensitization are mitigated or masked in the presence of *T. gondii*. In addition, *SOX21* appears to have effects on several neurologically relevant developmental pathways via “regulation of Wnt signaling pathway”. Beyond the trivial explanation of *T. gondii* infection rescuing the cocaine effect, the alternative explanation is that these may represent functions that the parasite needs to ensure are not activated through other stimuli.

Turning to the striatum, we identified three unique interacting subgroups (subgroups ‘f’, ‘g’ and ‘h’; **Figure 6A**). Subgroup f was functionally diverse, comprising 34 DE genes that responded more to *T. gondii* alone than to either cocaine or the combination of *T. gondii* and cocaine. Subgroup ‘g’ comprising 20 DE genes, responded more to cocaine alone than either *T. gondii* or the combination of *T. gondii* and cocaine. Significantly enriched annotations associated with this subgroup included GO terms relating to apoptosis and cell death as well as “modulation by symbiont of host apoptotic process” leading us to speculate that while cocaine exposure promotes cell death, *T. gondii* may act to suppress the hosts apoptotic mechanisms, mitigating cocaine toxicity while protecting *T. gondii* from the host’s attempts to clear the parasite. Finally, the 9 DE genes comprising subgroup ‘h’ were upregulated in response to cocaine but did not respond to *T. gondii* either in the presence or absence of cocaine. Significant enrichments in this subgroup suggest an involvement in TGFbeta signaling. TGFbeta has been associated with neuroinflammatory pathways and identified as neuroprotective following brain ischemia^58^. Here genes such as *NNROS* and *FBN1* associated with “negative regulation of transforming growth factor beta receptor signaling pathway” and “sequestering of TGFbeta in extracellular matrix” are upregulated by cocaine but this effect is supressed in the presence of *T. gondii* perhaps implying that the neuroprotective aspects of TGFbeta signaling are advantageous to the parasite’s persistence.

## DISCUSSION

Given that hyperactive dopaminergic function is a hallmark of schizophrenia, we developed a mouse model with associated behavioral tests to reveal altered dopaminergic sensitivity in *T. gondii* infected mice and focused on identifying the molecular pathways by which *T. gondii* infection contribute to disease pathogenesis. Using this model, we found that infected mice exhibited no change in memory or anxiety-associated behaviors. This contrasts with a previous study based on infection with strain ME49, in which the authors reported that infected mice spent significantly more time in the open arms of the EPM and an increase in exploratory behaviors in the open field (although time spent in the central zone of the field was reduced)^8^. Such differences may arise as a consequence of the breed of mouse or strain of parasite used. Here we used offspring from a cross between C57BL/6NTacfBr [C57B6] and 129Svev [129] mice, together with *T. gondii* strains VEG and CEP. The earlier study used B6CBAF1/J mice infected with the type II *T. gondii* strain, ME49^8^. The type III strains, VEG and CEP exhibit lower virulence in mice relative to ME49, and we used an initial inoculum of ∼500,000 parasites compared to 50-1000 parasites used in the other study. The decrease in anxiety and fear-related behaviors that have been observed in some previous studies following *T. gondii* infection are typically described as behavioral manipulations that facilitate predation mediated transmission of the parasite from intermediate to definitive hosts. However, there are other behavioral changes that may serve the same purpose such as decreased motor activity or coordination as we observed here.

Beyond memory and anxiety-associated diseases, we further found that infection with *T. gondii* significantly reduced locomotor activity levels in both the open field test and the elevated plus maze and impaired rotarod acquisition consistent with previous studies^29^. Finally, cocaine sensitization followed by prepulse inhibition resulted in *T. gondii* infected mice showing an equivalent initial locomotor response to cocaine (day 1) but exhibited blunted sensitization upon repeated cocaine administration. Furthermore, while drug free control and infected mice show normal activity level, infected mice exhibited reduced sensitization to cocaine after one week of abstinence, as well as displaying exaggerated prepulse inhibition compared to control mice.

In terms of behavior, our primary goal in the current study was to determine the impact of *T. gondii* infection on behaviors known to be impacted in neuropsychiatric conditions. Our findings in the cocaine sensitization and PPI experiments clearly indicate that *T. gondii* alters the normal pattern of behaviors in these tasks although these effects were unexpectedly in the opposite direction of what is typically observed in individuals with schizophrenia. Interestingly, we noted that *T. gondii* infection resulted in a lack of sensitization in mice upon cocaine challenge. Such depressed sensitization may be driven through the persistent activation of microglia, resulting in the elevated expression of TNF-alpha and subsequent internalization of synaptic AMPA receptors ^28^. In any case, given the magnitude of *T. gondii* infection as a risk factor for schizophrenia, the infection related behavioral sensitization to cocaine and the corresponding changes in gene expression may provide critical insight into disease etiology.

Following behavioral tests, gene expression profiling on two regions of the mouse brain, hippocampus and striatum, identified 4,439 unique genes that are significantly differentially expressed in mice in response to chronic *T. gondii* infection and/or pre-sensitization to cocaine. At least 2024 of these genes are associated with neurologically relevant functional annotations implicating a broad range of potential contributing pathways linked to neuroinflammation, neurotransmission, memory and cognitive function, locomotion, neurodevelopmental pathways, synaptic function, myelination, oxidative stress, and others. Strikingly, while some differences were observed in the distribution of detail level annotations associated with hippocampus and striatum, at a high level the affected pathways were broadly similar across tissues suggesting a general response to both treatments, although technologies such as single cell RNASeq would help resolve cell-specific responses.

Most expression changes we observed consisted of genes responding positively to either *T. gondii* or cocaine, suggesting that while they may have complimentary effects on similar functional pathways, the specific genes/pathways on which they act are different. This is advantageous from a model perspective since it is widely believed that *T. gondii* represents a risk factor rather than a causal factor in the onset of SZ. Only a small number of statistically significant interactions, with the ability to complicate interpretation, were identified. In these cases, we speculate that *T. gondii* and cocaine affect gene regulation in pathways that intersect in complex ways. For example, we identified genes where cocaine treatment or *T. gondii* infection increased expression, while their combination had no impact on expression (Functional effect group e), suggesting opposing effects on parallel pathways, while for another set of genes the influence of *T. gondii* on expression dominates over the expression difference associated with cocaine treatment (e.g., Functional effect groups c, g and h), perhaps indicative of different targets within a single pathway.

It has been suggested that *T. gondii* impact neurodegenerative diseases through modulating the activity of neurotransmitters, particularly dopamine. For example, chronic (but not acute) infection with *T. gondii* increases local concentrations of dopamine in the brains of mice similar to patients suffering from schizophrenia^59^. Such increases may be driven through the expression of aromatic acid hydroxylase genes by *T. gondii*, which appear similar to mammalian tyrosine hydroxylase, a rate-limiting enzyme in dopamine synthesis^60, 61^. Given we were unable to detect significant expression of *T. gondii* genes in our samples, we are unable to comment on the likelihood that exogenous manipulation of dopamine by *T. gondii* may contribute directly to the behavioral effects observed. However, we have demonstrated that chronic infection can be established in mice following a high-dose injection, leading to observable tissue cysts that were visualized in this study by imaging the accumulation of the GFP-tagged *T. gondii.* In addition, cocaine is said to elicit SZ-like symptoms in mice and acts through the dysregulation of dopamine^27^; a paradigm we leveraged for our analysis. Notwithstanding the importance of dopamine, there is reason to suspect that *T. gondii* may leave a larger neurological footprint. Various neurotransmitters including serotonin (5-HT), GABA, glutamate, as well as other signals have been proposed to play contributing roles^18, 37, 62^. Testosterone, for example, has been shown to be elevated in *T. gondii* infected mice and humans and is presumed to be associated with aggressiveness and more dominant behaviors^11^.

Another contributing mechanism that has been proposed is the disruption of tryptophan metabolism. *T. gondii* is a tryptophan auxotroph and like many micro-organisms must scavenge this amino acid from their host. The immune system has evolved a method to degrade tryptophan which is effective in restricting the growth of many such pathogens. This is achieved through the production of indoleamine 2,3-dioxygenase (IDO) in immune cells, which degrades L -tryptophan to N-formylkynurenine^18, 37,62^. However, this has the effect of increasing the production of other catabolites such as 3-hydroxykynurenine and, anthranilic, xanthurenic, quinolinic and kynurenic acids. Some of these affect neurons or their functions. For example, quinolinic acid is an agonist of the N-methyl-D-aspartate (NMDA) receptor whereas both 3-OH-kynurenine and quinolinic acid can cause neuronal death by a mechanism mediated by reactive oxygen species^63^. In addition, high concentrations of kyneurenic acid (KYNA) have been detected in the cerebrospinal fluid of schizophrenic patients and may contribute to SZ associated cognitive deficiencies^37^. Here we find that two adjacent enzymes in this pathway, kynureninase (KYNU) [EC:3.7.1.3] and 3-hydroxyanthranilate 3,4-dioxygenase (HAAO) [EC:1.13.11.6] are indeed over-expressed in response to *T. gondii*, suggesting that enhanced Tryptophan catabolism is at play in our model. Finally, astrocytes which are preferred by *T. gondii* for replication, are the principal source of kynurenic acid (KYNA) and are known to be activated in response to latent toxoplasmosis.

In analyzing functional annotation enrichments, we did not exclude any evidence categories from the GO analysis in our assessments. Given the robust (and expected) immune response we felt it necessary to extend our search for potential neurological associations among the DE genes detected. In general, while we found that GO was the most detailed resource for functional annotations, it was also the least readily interpretable due to the difficulty of visualizing the vast number of related terms in the ontology especially given the broad immune response that due to multiple correction testing, masked many relevant terms. The body of knowledge underlying the GO is itself very heterogeneous. One problem is that many gene products are involved in multiple cellular processes, leading to a lack of comprehensive annotation of all gene products and GO enrichment analyses require accurate specification of the background distribution to yield meaningful results. With respect to neurological annotations, several recent and ongoing efforts have attempted to correct the problem of sparse neurological annotations^50–52^. Thus, difficulty in extracting neurologically relevant terms at statistically significant levels was unlikely due to lack of attention to neurological annotations. Rather, we found that focusing our analysis on the relevant functional context increased the statistical power of term enrichment and identified a more focused DE gene subset associated with a broad set of neurologically relevant associations.

It would be interesting to test *T. gondii* in other mouse genetic backgrounds e.g., COMP deficient mice which have increased dopamine in the prefrontal cortex but not the striatum due to a deficient catechol-o-methyl transferase. COMP inhibitors improve working memory in rodents and humans. A COMP polymorphism resulting in 4x activity occurs in high rates in schizophrenics and their unaffected siblings. High activity impairs prefrontal cognition and may increase risk of schizophrenia. Similarly, there is some evidence that cannabis use is associated with increased risk for psychotic disorder and symptoms. Cannabis-induced acute psychotic states treated in psychiatric services are considered early signs of schizophrenia and related disorders.

While this study highlights some exciting associations between *T. gondii* infection and pathways involved in neurocognitive disorders, we nevertheless note several limitations in our study. First, we were unable to monitor the expression of genes by the parasite. Despite being relatively slow growing, the bradyzoite form of the parasite expresses and secretes several proteins that can impact host tissue function. Second, many genes lack functional annotations which limits our ability to resolve functions at the level of individual conditions, as well presenting challenges in the recovery of significant terms related to neurophysiology. Third, we did not monitor levels of neurotransmitters which may otherwise have allowed us to correlate their expression with changes in gene expression. Fourth, the presence of complex *T. gondii* by cocaine interactions though limited, while interesting, are challenging to interpret. Finally, we applied a liberal cutoff (FDR < 0.1) to define DE genes to ensure that our gene set capture a broad view of gene expression patterns and avoid being overwhelmed by immune response pathways. While we acknowledge that this approach increases the number of false positives, we note this approach allowed the recovery of neurophysiologically relevant terms that would otherwise be masked by immune-related terms.

## CONCLUSIONS

We present a novel mouse model to explore molecular mechanisms underlying neurological risk factors associated with chronic *Toxoplasma gondii* infection. We have shown that chronic infection by the parasite, in combination with cocaine, produces clear alterations in behavioral paradigms that are commonly impaired in neurological disorders. Numerous behavioral changes have previously been observed following *Toxoplasma gondii* infection. Here we show a novel behavioral phenotype, that is blunted behavioral sensitization to the stimulant effects of cocaine that strengthens the link between infection and neuropsychiatric related behaviors. In addition, we identified 4439 genes that were significantly differentially expressed under these conditions. Through a systematic, curation approach we found nearly half of these genes were enriched for potentially neurologically relevant GO biological processes, highlighting potentially relevant pathways. We further demonstrated, using a more focused, novel context enrichment approach, a rapid screening technique that identified 641 neurologically relevant genes, including 114 that were not picked in the initial GSEA due to its low statistical power. Through demonstrating the capacity of this model to yield a wealth of new data on genes and pathways implicated to respond to infection by *T. gondii*, we expect further use of this model will allow the dissection of the molecular pathways by which infection with *T. gondii* contributes to disease pathogenesis enabling more efficient therapy and prevention of schizophrenia.

## METHODS

### Ethics Statement

All procedures were conducted in accordance with policies of the Hospital for Sick Children Animal Care Committee and conformed to both Canadian Council on Animal Care (CCAC) and National Institutes of Health (NIH) Guidelines on Care and Use of Laboratory Animals.

### Experimental Animals

Mice used in this study were the F1 progeny of a cross between C57BL/6NTacfBr [C57B6] and 129Svev [129] mice (Taconic Farms, Germantown, NY). Mice were bred in our colony at The Hospital for Sick Children and maintained on a 12 h light/dark cycle with access to food and water *ad libitum*. A total of 39 mice were used for behavioral experiments with CEP, 40 mice were used for behavioral experiments with VEG and an additional cohort of mice were used for the RNA-seq experiments; one mouse treated with *T. gondii* and cocaine died prior to sampling.

#### T. gondii culture

Two type III strains were used in this study: 1) strain VEG; and 2) a CEP strain which had been engineered to contain a stably integrated GFP cassette ^64^. Strains were propagated by passage in monolayers of primary human foreskin fibroblast cells (ATCC SCRC-1041), in Dulbecco’s modified Eagle’s medium (Wisent 319-015-CL) supplemented with 10% cosmic calf serum (HyClone SH30087.04), 20% M199 (Wisent 316-010-CL) and either penicillin/streptomycin (Wisent 450-201-EL) or, 10 U/mL penicillin and 10 μg/ml streptomycin (Gibco, Thermo Fisher Scientific Inc., Grand Island, NY), as previously described^65^. Genomic material was extracted, and PCR tested to ensure the lines were free from Mycoplasma infection. Tachyzoite-stage parasites were collected in 14 ml phosphate buffered saline (PBS) for injection into mice.

### Infection

Adult mice, 60 days of age were injected intraperitoneally (IP) with 150 μl of PBS containing approximately 500,000 tachyzoites from either strain of *Toxoplasma gondii* (GFP tagged CEP or VEG). Control mice were injected with PBS only. Mice were given an 8-week recovery period prior to behavioral testing and genomic analysis, during which the acute phase of infection was expected to subside, leading to chronic infection and associated tissue cyst formation.

### Behavioral Experiments

Following the 8-week recovery period after injection, mice were assessed for fear and anxiety related behaviors (contextual fear conditioning, elevated plus maze, open field), motor ability (Rotarod) and neuropsychiatric related behaviors (including cocaine sensitization and prepulse inhibition). Behavioral procedures were conducted during the light phase of the cycle. Experiments were conducted blind to the treatment condition of the mouse, and according to local animal care protocols.

#### Cocaine Sensitization

For cocaine sensitization experiments, mice were habituated to an open arena for three days (5 minutes per day). Then, mice were injected with 20mg/kg of cocaine daily for 5 days. Cocaine or saline was injected 5 minutes before being placed in an open field (4o cm x 40 cm). Locomotor activity was recorded over a period of 5 minutes using an overhead camera connected to an automated tracking system (Limelight software, Actimetrics). One week after the end of the sensitization the mice were tested in the same open field with locomotor activity recorded. All mice were first tested in a drug free state followed by a 20mg/kg cocaine challenge 24 hours later.

#### Prepulse Inhibition (PPI)

SR-Lab startle chambers (San Diego Instruments) were used to test PPI. Mice were first placed in individual plexiglass cylinders (internal diameter 3.2 cm) which were then placed on a sensor platform inside a sound-attenuating chamber. A piezoelectric accelerometer on the base of the platform was used to detect all movements of the mouse inside the cylinder. Background white noise (65dB) was presented throughout the testing session. After 2 minutes of acclimation to the chamber the mice were presented with a series of trials of either startle stimulus alone (40 ms duration, 120dB tone), prepulse stimulus alone (20 ms tone) or prepulse and startle stimulus pairings. The mice were presented with 5 different prepulse intensities (69 dB, 73 dB, 77 dB, 81 dB or 85 dB). For each prepulse intensity there were 12 prepulse only trials and 12 prepulse/startle trials. There were also 24 startle-only trials. Trial types were intermixed and an intertrial times of 15 s was used.

#### Elevated Plus Maze

The elevated plus maze consists of a plus-shaped apparatus with 2 open and 2 enclosed arms (10 cm wide, 50 cm long), each with an open roof, elevated 1 m from the floor. Mice were placed in the centre of the maze and allowed to explore for 5 mins. Their movements were tracked by an overhead camera and analyzed using Any-maze software. The proportion of time spent in the closed arms was calculated (time in closed/total time) and used as a measure of anxiety. In EPM, this translates into a restriction of movement to the enclosed arms.

#### Contextual fear conditioning

Contextual fear conditioning was performed in Med-associates conditioning chambers (31 cm x 24 cm x 21 cm) containing shock-grid floors. During training mice were placed in the chamber and were allowed to aclimitize for 2 minutes. Then, they received 3 foot-shocks (0.5 mA, 2 s duration spaced 1 minute apart) and were removed from the chamber 1 minute after the final shock. The following day, mice were placed back in the same conditioning chamber for 5 minutes. No shocks were delivered, and the time spent freezing (defined as the absence of all movement other than respiration) was recorded using Freeze Frame analysis software (Actimetrics).

#### Rotarod

Motor learning was assessed using an accelerating protocol on an automated Rota-rod system (Med-Associates). Mice were trained for 5 days with 4 trials per day. The maximum trial length was 5 minutes during which time the speed of the rotation accelerated linearly from 4 to 40 revolutions per minute. The length of time the mice stayed on the rod before falling was recorded for each trial. Mice were given an approximate intertrial interval of 10 minutes.

### Tissue Collection

The hippocampus and striatum were dissected from mouse brains, frozen in dry ice cooled isopentane and stored at -80C. Frozen tissues were homogenized first through bead-beating in RLT lysis buffer (Qiagen RNeasy kit, cat. 74104) using a single pre-chilled 5mm steel bead (Qiagen cat. 69989) and the Qiagen TissueLyser for 2x 2 min at 30 Hz, then using QIAshredder spin columns (Qiagen cat. 79654).

### RNA-Seq

Total RNA extraction was carried out using the Qiagen RNeasy kit (cat. 74104) with on-column *DNase*I treatment, as per the manufacturer’s instructions. Quality of RNA samples were checked on an Agilent Bioanalyzer 2100 RNA Nano chip following Agilent Technologies’ recommendation. Concentration was measured by Qubit RNA HS Assay on a Qubit fluorometer (ThermoFisher). Yields ranged from 214 ng/ul x 40 ul to 548 ng/ul x 40 ul. Only samples with RNA integrity number > 6.9 and little to no degradation apparent on electrophoretograms were accepted. Library preparation and sequencing were performed at The Centre for Applied Genomics (TCAG; Toronto, ON). Library preparation was performed following the NEBNext Ultra II Directional RNA Library Preparation protocol (New England Biolabs, Ipswich, MA). Briefly, 800 ng of total RNA was enriched for poly-A mRNA, fragmented into the 200-300-bases range for 4 minutes at 94°C and converted to double stranded cDNA, end-repaired and adenylated to create a 3’ overhang for ligation of Illumina adapters. Following PCR amplification and extension, each sample contained a different barcoded adapter to allow for multiplex sequencing. One ul of each of the RNA-seq libraries was loaded on a Bioanalyzer 2100 DNA High Sensitivity chip (Agilent Technologies) to check for size and absence of primer dimers. RNA libraries were quantified by qPCR using the Kapa Library Quantification Illumina/ABI Prism Kit protocol (KAPA Biosystems), pooled in equimolar quantities and single-end sequenced on a HiSeq 2500 platform, using 5 lanes of a high output flow cell with the v4 chemistry following Illumina’s recommended protocol to generate single-end reads of 100-bases in length (40 million reads per sample; 250-270 million reads per lane).

### Data Processing

#### Pre-processing and Annotation

We first processed sequence reads by removing adaptor and vector contaminated reads and trimmed low quality reads using Trimmomatic v.0.36^66^. Next, quality reads were mapped to the reference genomes *M. musculus* (Source: Ensembl; Taxonomic ID: 10090, Release 87) and *T. gondii* (Source: Ensembl; Taxonomic ID: 432359, Release 34) using STAR^30^. The expression level of annotated genes was assumed to be proportional to the number of mapped sequence reads. Expression levels of mouse genes were normalized across samples with reads per kilobase per million reads (RPKM).

#### Statistical Analysis

All gene differential expression analyses were performed using the DESeq2 R Package version 1.22.1 and R version 3.5.3^67^. Count normalization size factors were calculated across all samples using the median geometric mean-scaled count method (default method). These ‘global’ size factors were then used in all analyses. The following analysis was performed separately on the striatal and hippocampal samples. DE testing was performed using likelihood ratio tests on different negative binomial general linear models of gene expression. Briefly, likelihood ratio tests assess the significance of model terms by comparing the full model to a reduced model where those terms are missing. In order to assess the significance of *T. gondii* main effects, a likelihood ratio test was done for each gene comparing the performance of the model “gene expression ∼ *T. gondii* + cocaine” versus the reduced model “gene expression ∼ cocaine”. To assess cocaine main affects, a full model of “gene expression ∼ *T. gondii* + cocaine” was compared to the reduced model “gene expression ∼ *T. gondii*”. Finally, to assess the significance of *T. gondii* by cocaine interactions, a full model of “gene expression ∼ *T. gondii* + cocaine + *T. gondii* ’ cocaine” was compared to the reduced model “gene expression ∼ *T. gondii* + cocaine”. False discovery rate (FDR) correction was performed on p-values across all genes separately within each different likelihood ratio test (i.e., *T. gondii* main effect, cocaine main effect, and *T. gondii* by cocaine interaction), with an FDR cutoff of 10% taken for statistical significance. Significant results were grouped into 4 main categories: 1) significant *T. gondii* by cocaine interactions, 2) significant main effect of *T. gondii*, significant main effect of cocaine, and no significant *T. gondii* by cocaine interaction, 3) significant main effect of *T. gondii* only (no significant main effect of cocaine and no significant *T. gondii* by cocaine interaction) and 4) significant main effect of cocaine only (no significant main effect of *T. gondii* and no significant *T. gondii* by cocaine interaction). Groups 2-4 were further broken down based on whether *T. gondii* and cocaine main effects consisted of upregulation or downregulation in response to *T. gondii* or cocaine. Genes with significant interactions were clustered into unique interaction types using log2 fold-change signed p-values from pairwise Wald differential abundance tests between groups. Heatmaps of significant genes were generated using mean-centered and standard deviation-scaled log2 values of the DESeq2 size factor normalized counts (See **Supplementary Figures 4-11** and **Supplementary Files 3, 4**). An overview of the expression values and patterns may be found in **Supplementary Spreadsheet S1** in which categories are grouped and labeled alongside a heatmap showing their expression patterns.

To determine whether there were significant differences in the number of differentially expressed genes within gene subsets or conditions for each tissue, Fisher’s Exact Tests were performed in R (version 3.5.3). The expected values were calculated based on the null hypothesis that condition had no effect on the totals. Rather, the expected values in each category reflect the relative frequency of hippocampus-specific and striatum-specific genes for all differentially expressed genes. Fisher’s Exact Tests were likewise also performed to determine whether significant differences existed between the expected and observed number of *T. gondii* and cocaine responsive genes in hippocampus and striatum (**Supplementary File 4**). For the high-level functional analysis, z-scores were calculated in R (version 3.5.3) based on the mean differences between the number of hippocampus and striatum genes for each top-level Reactome pathway (**Supplementary Figure 2B**).

#### Functional analysis

To test if GO terms were enriched within DE genes and their subsets, gene set enrichment analysis was performed with the Cytoscape plugin, BiNGO^35, 68^ using the Benjamini-Hochberg false discovery rate procedure (FDR<0.05) for multiple testing correction. **Supplementary File 5** contains raw enrichment results for various DE gene subsets. To aid interpretation additional pathway annotations and ontologies were obtained from Reactome and KEGG^69^ and used to construct custom ontologies for input to BiNGO and gene set enrichments were performed as above. All ontology files and annotations used for this analysis are provided as Supplementary File 6. Note that we excluded KEGG Drug Development pathways as irrelevant to our analysis of gene biological function. Due to the large occurrence of generic, high-level immune system processes, significant GO terms were then curated to assess their potential neurological relevance according to the method below (see Curation). Raw data for bar charts and intersects were compiled in MS Excel and arranged for visualization using Python 3 scripts written in house. Prepared files were imported for visualization in RStudio and plotted using the R libraries: UpSetR, rJava, grid, tidyverse and venneuler. Intersects for genes in term groups were plotted in R using UpSetR with nintersects=45.

Context enrichment was performed as follows. We first defined a set of root keywords (“neuro”, “brain”, “behavior”, “locomot”, “memory”, and “synap”) which we considered, based on our curation of the literature, and in the context of our experiments to be both relevant and non-immune system specific. We then performed an AmiGO^48^ search for GO biological process terms containing these keywords among their full-text descriptions (see **Supplementary Figure 5** and **Supplementary File 8**). The resulting 1632 GO terms defined the functional “context” and were used to filter the DE gene set by removing from consideration those that did not contain at least one annotation matching the defined context. GO enrichment analysis was then performed on this subset per above using both the full GO ontology and NIGO, a neuro-immune focused GO subset developed using a clipping approach based on domain-specific relevance^47^ obtained from BioPortal^70^. We obtained the list of genes with differentially methylated positions associated with psychosis, schizophrenia and treatment-resistant schizophrenia from supplementary material in Hannon et al.^49^ The expected overlap between this set and our context enriched set was assessed using Fisher’s Exact Test assuming an estimated background frequency of 658/20,648 for SZ-associated methylated mouse genes.

#### Curation

Neurological relevance of statistically enriched GO terms occurring in DE gene annotations was confirmed by curation as follows. First, terms containing words indicative of neuronal function e.g., brain development, behavior, neuroinflammation or brain-specific cells or processes, were included. Second, using the form *pathway* + “in brain” e.g., “NF-kappaB in brain”, we conducted Google searches (i.e., page rank) and manually reviewed scholarly sources included in first page results for supporting evidence of neurologically associated function. Scholarly sources were defined as primary scientific literature accessible on PubMed or, reputable aggregate sources (e.g., GeneCards^71^, RefSeq^72^ or, Uniprot^73^) citing primary sources. Related terms were grouped based on GO hierarchy with each term being represented only once. For each experimental condition, a ‘1’ was scored where at least one neurologically relevant term in a given group was enriched (see **Supplementary File 7**).

## ACKNOWLEDGEMENTS

We would like to thank Dr. Steven Maere, Ghent University for providing helpful advice on the creation of custom annotation files for use in BiNGO.

## SUPPLEMENTARY FIGURES

**Suppl. Figure 1:**
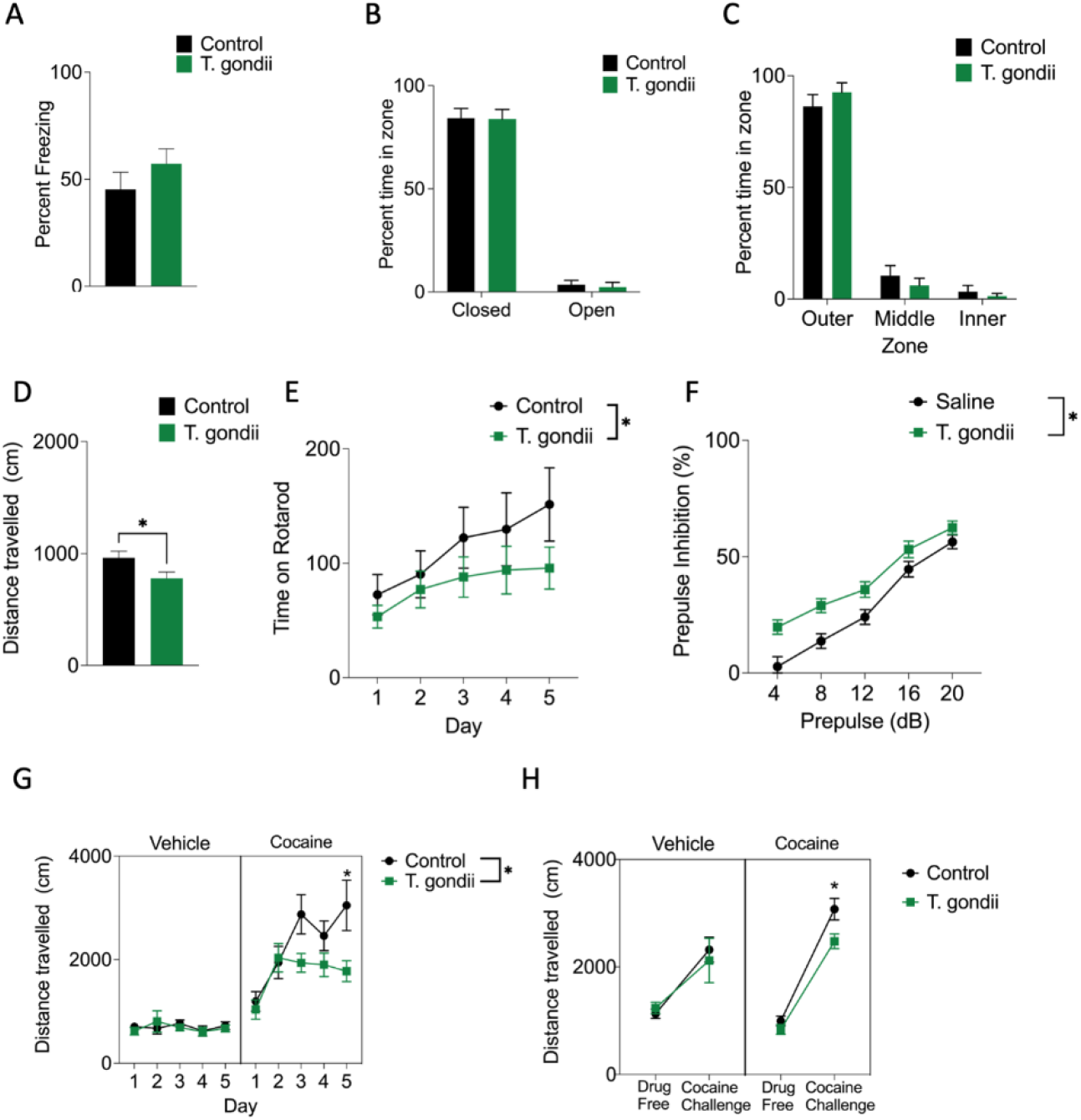
Behavioral testing of T. gondii (VEG) infected mice. A - Infected mice are not impaired in a contextual fear conditioning (t-test, *p =* 0.28) B-C-Infected mice did not show altered anxiety behaviors in the elevated plus maze (Two-way ANOVA No significant main effect of group (*p = 0.83)*or region by group interaction *p. =0.92* ) (B) or open field test (Two way ANOVA No significant main effect of group (*p =* 0.99) or group by zone interaction (*p = 0.34))* (C). D-infected mice exhibited a significant reduction in distance travelled in the open field (t-test t(18) = 2.24, *p = 0.038*). E – There was also a significant decrease in the time spent on the rotarod compared to control mice (Repeated measures ANOVA, main effect of group F(4,90) = 5.1, *p= 0.026.* group by session interaction *p* = 0.89. F – A reduction in prepulse inhibition was observed in infected mice compared to controls (Repeated measures ANOVA main effect of group F(4,90) = 31.69, *p ≤ 0.0001* group by intensity interaction *p =* 0.45). G-H – As was observed with the CEP strain, infected mice had a blunted locomotor response to repeated cocaine administration significant group by drug treatment effect F(1,80) = 8.09, *p =* 0.0057)(G) however, there was no significant group or group type interactions (*p’s ≥ 0.14)* during a subsequent cocaine challenge one week following sensitization (H).

**Suppl. Figure 2:**
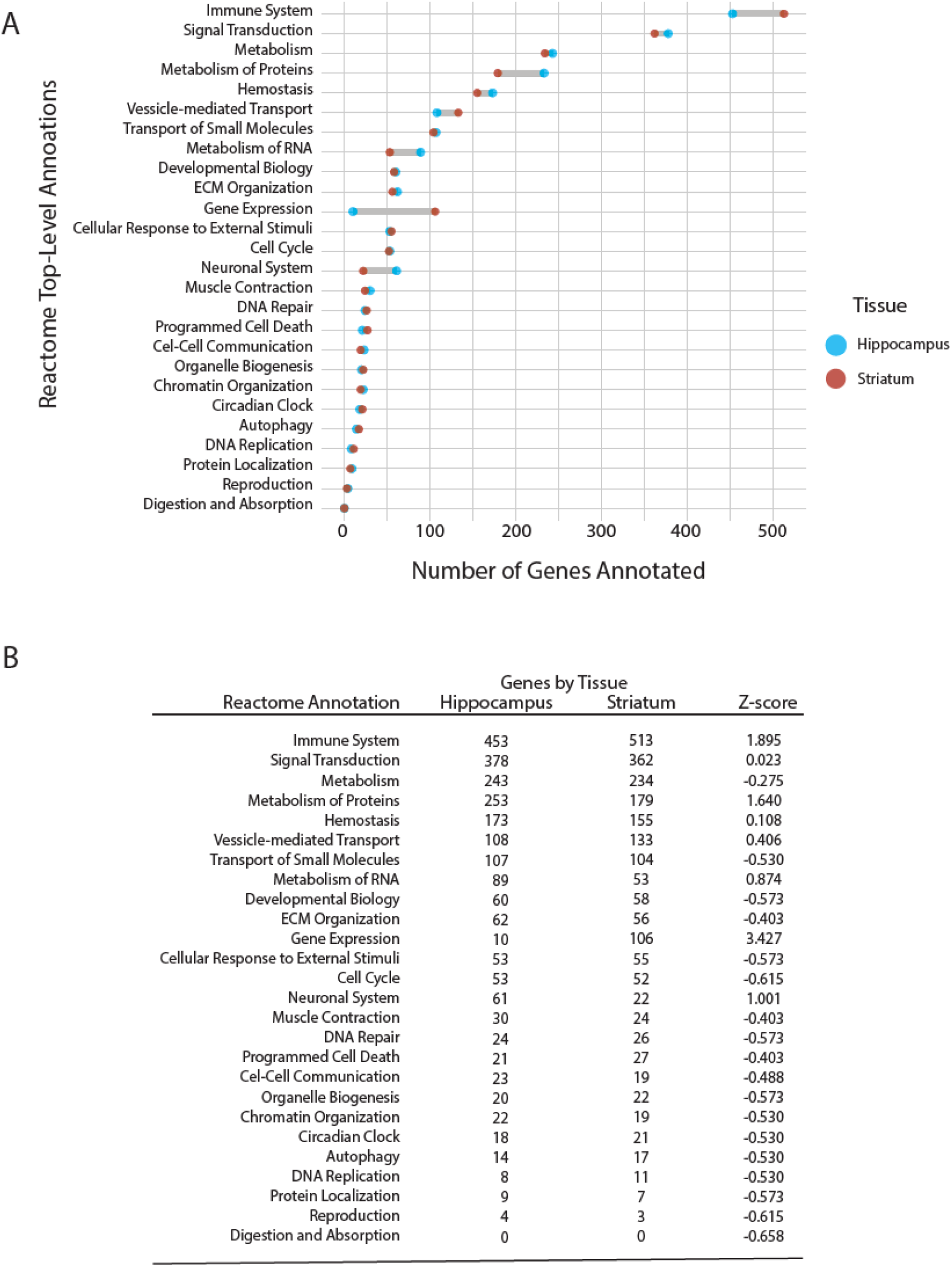
Enumeration of DE genes annotated to top-level Reactome Pathways. A - Cleveland Dot Plot comparing the number of DE genes annotated to top-level Reactome Pathways in hippocampus (blue) and striatum (red), ordered by decreasing average number of genes. B - Numbers of genes by category by tissue plotted in A.

**Suppl. Figure 3:**
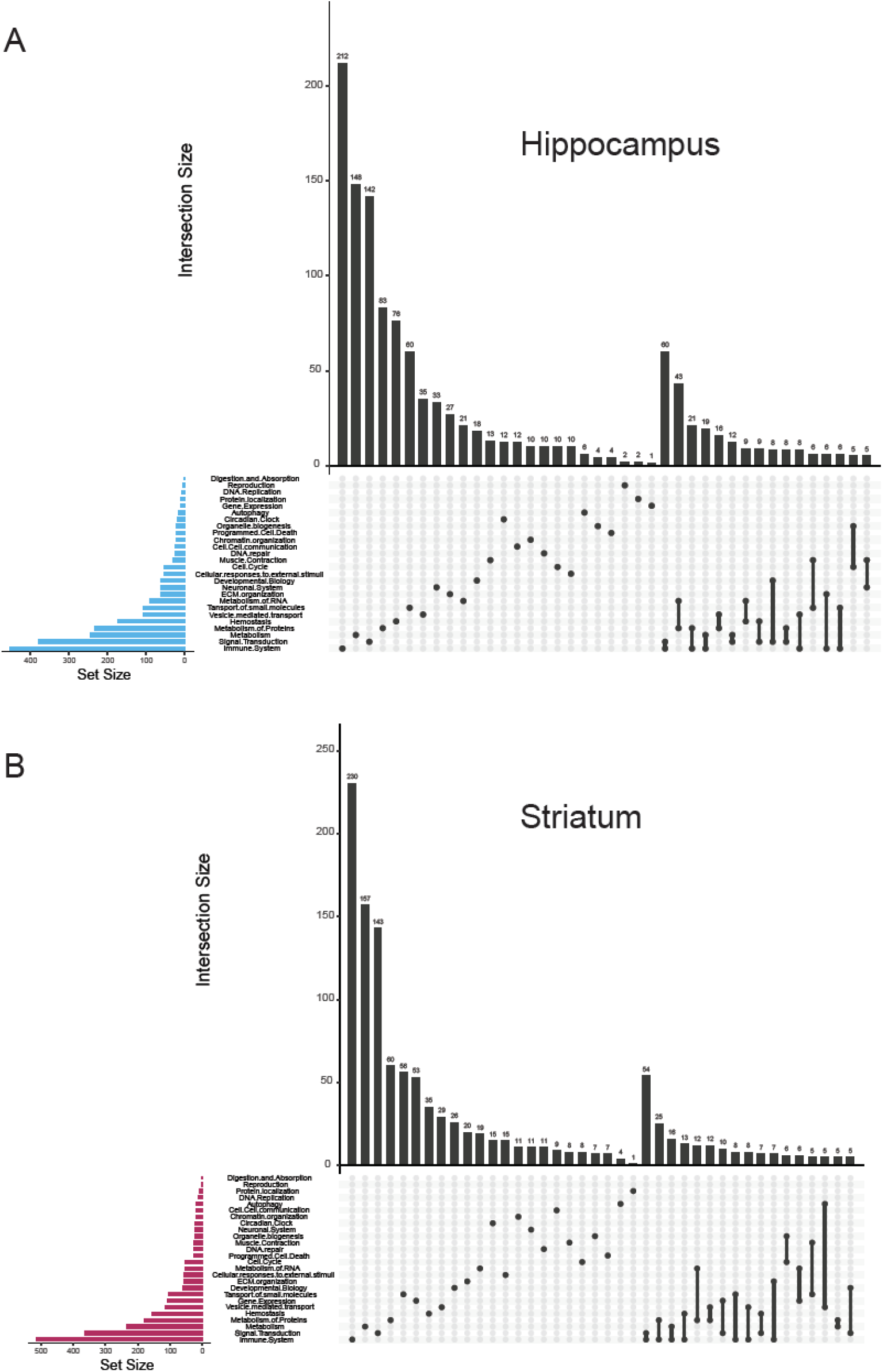
Overlap among Reactome Pathway annotations for DE genes. Upset Plots show overlaps (n>4) in genes annotated to multiple top-level reactome categories for hippocampus (A) and striatum (B).

**Suppl. Figure 4:**
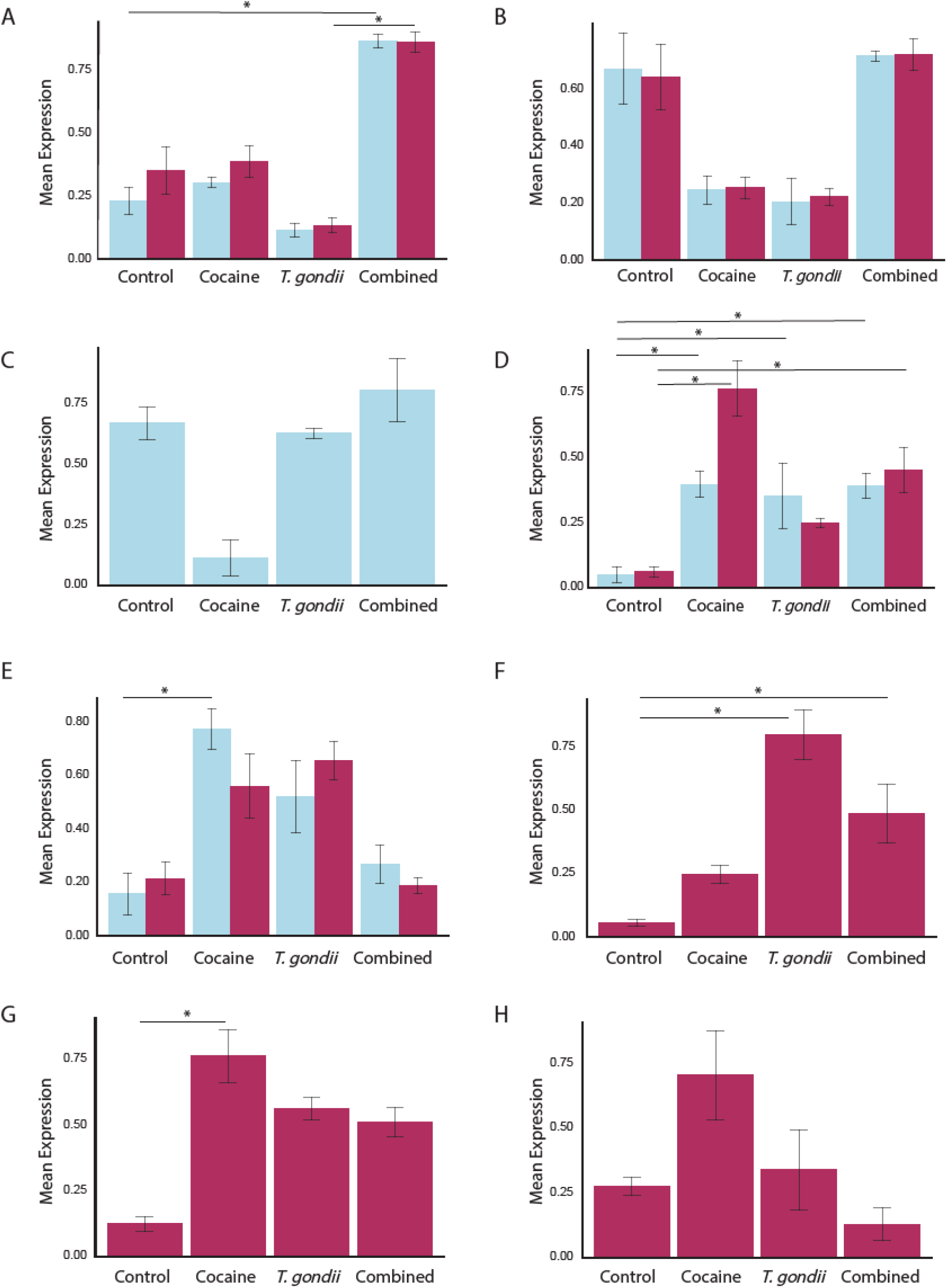
Expression patterns of interacting DE genes by functional effect group (A to H). Significant differences are indicated by an asterisk (Chi square; p < 0.05).

**Suppl. Figure 5:**
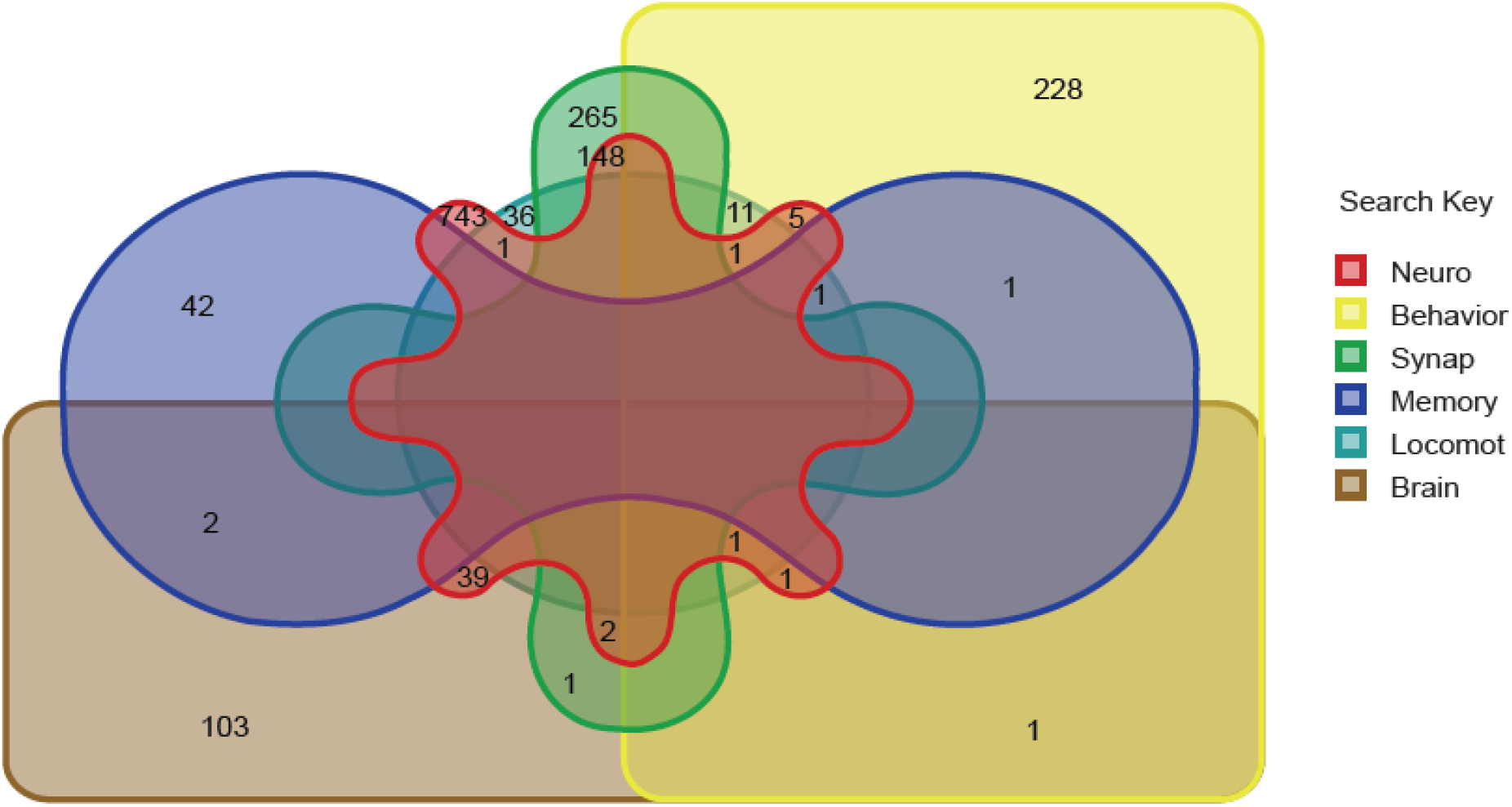
Numbers of GO terms returned in neurological keyword searches with overlaps.

**Suppl. Figure 6:**
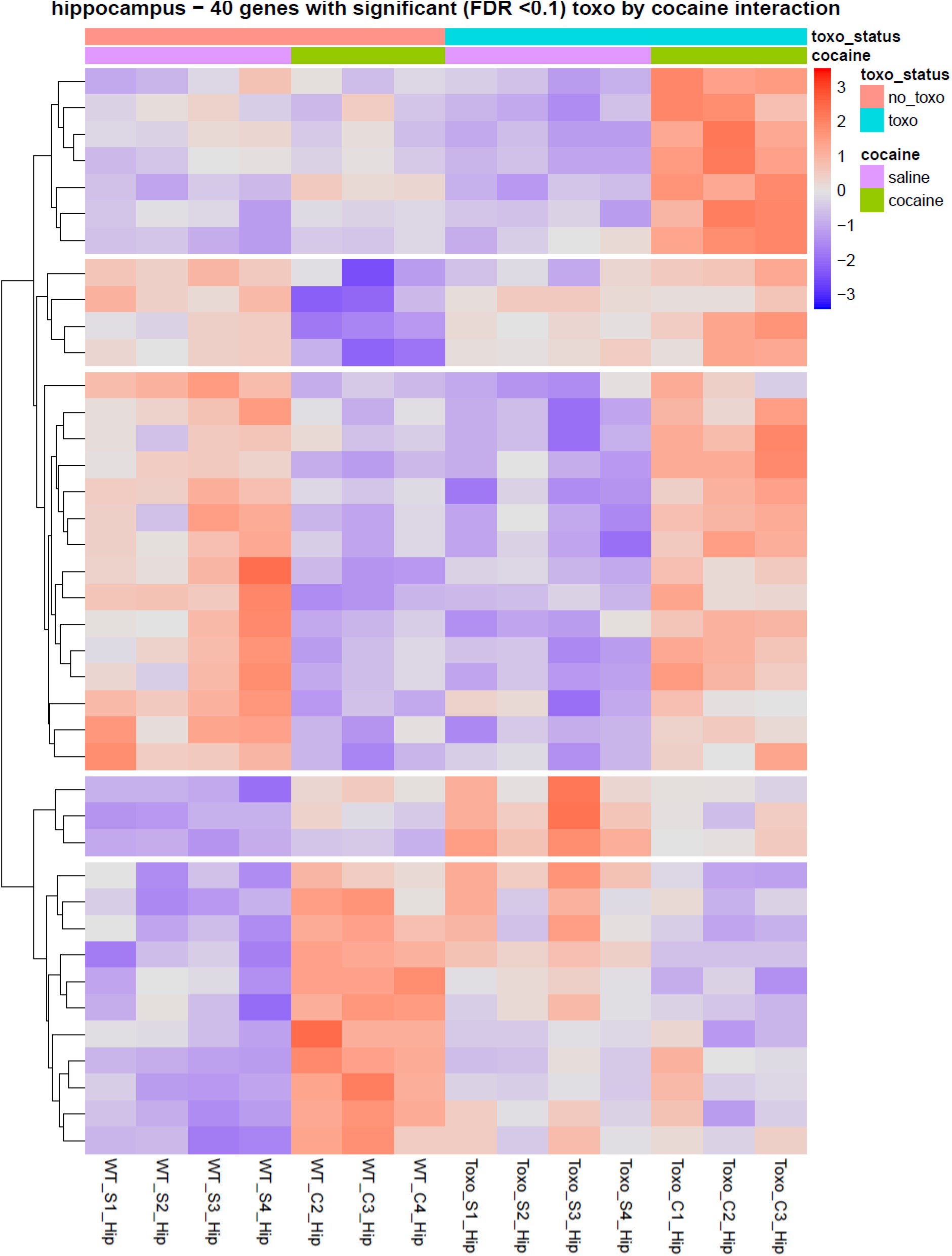
Heatmaps of gene expression defining DE gene responses by type and subtype.

**Suppl. Figure 7:**
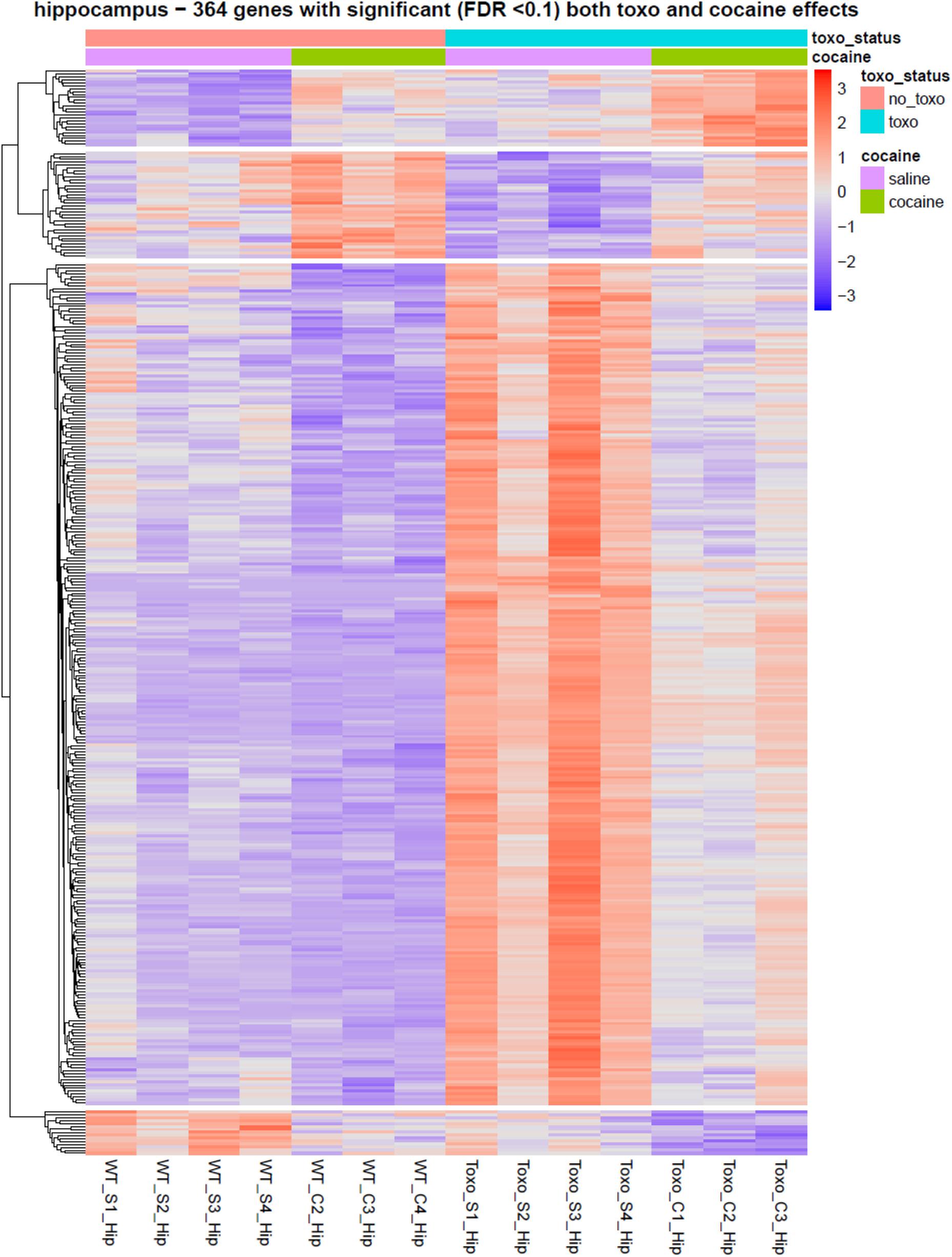
Heatmaps of gene expression defining DE gene responses by type and subtype.

**Suppl. Figure 8:**
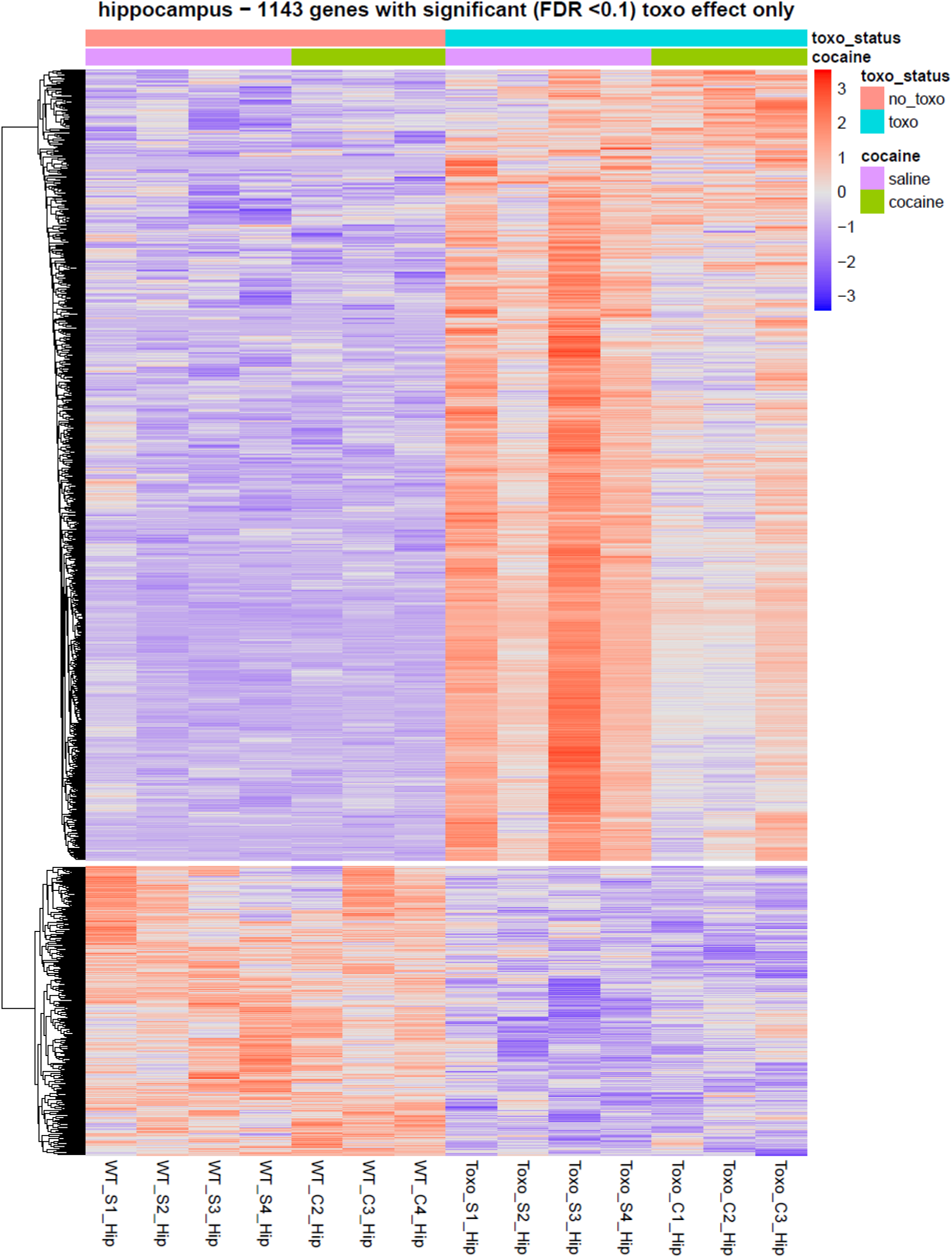
Heatmaps of gene expression defining DE gene responses by type and subtype.

**Suppl. Figure 9:**
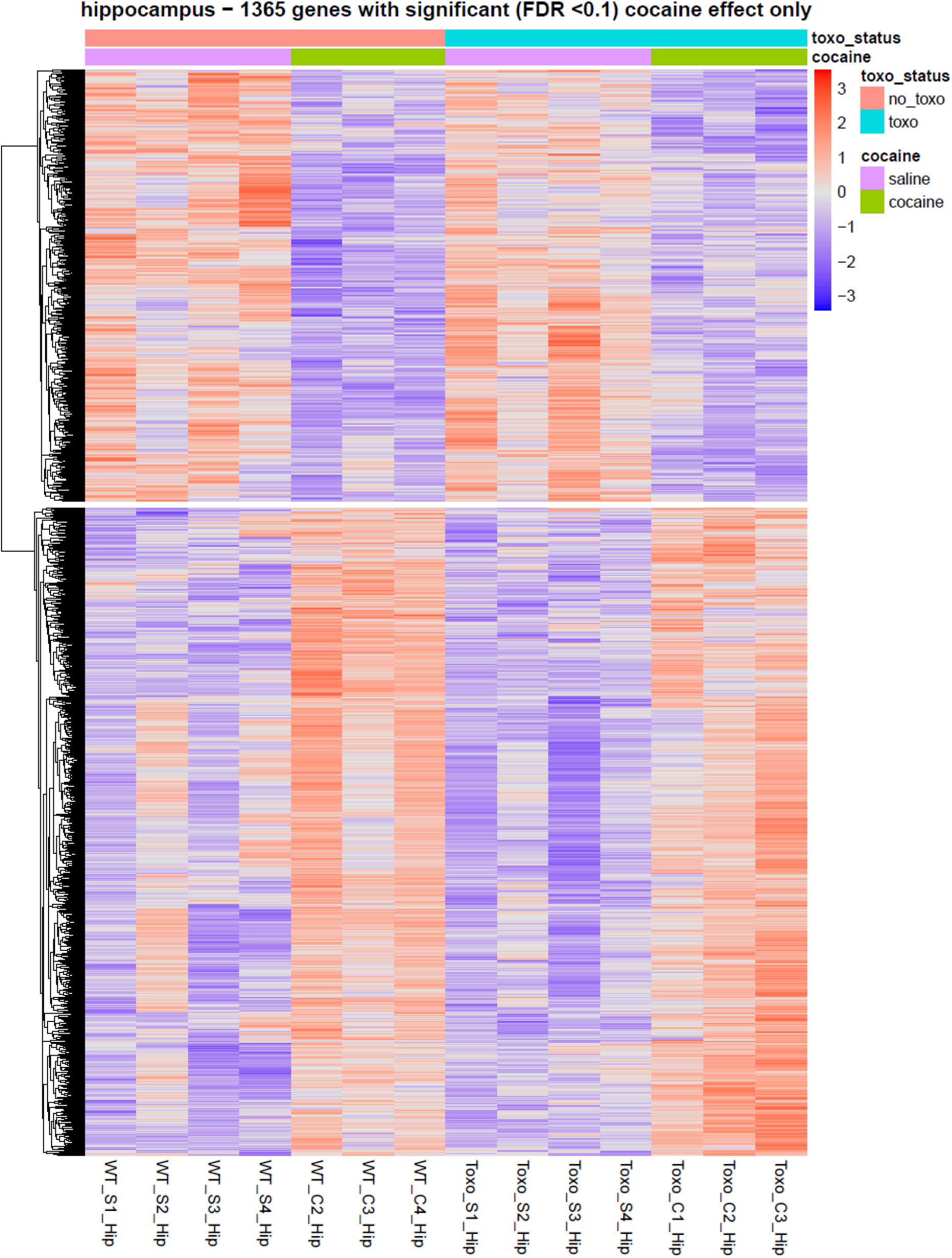
Heatmaps of gene expression defining DE gene responses by type and subtype.

**Suppl. Figure 10:**
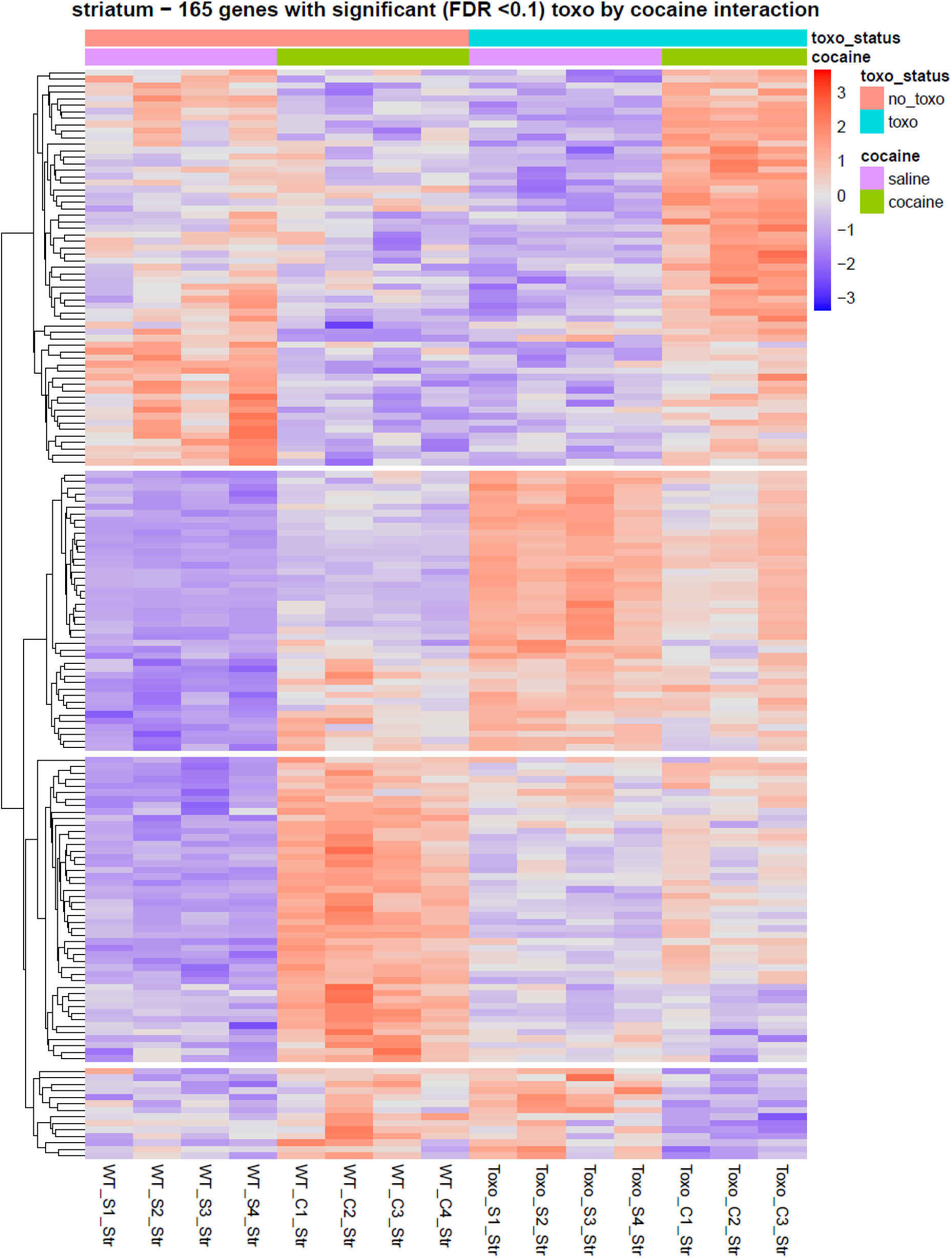
Heatmaps of gene expression defining DE gene responses by type and subtype.

**Suppl. Figure 11:**
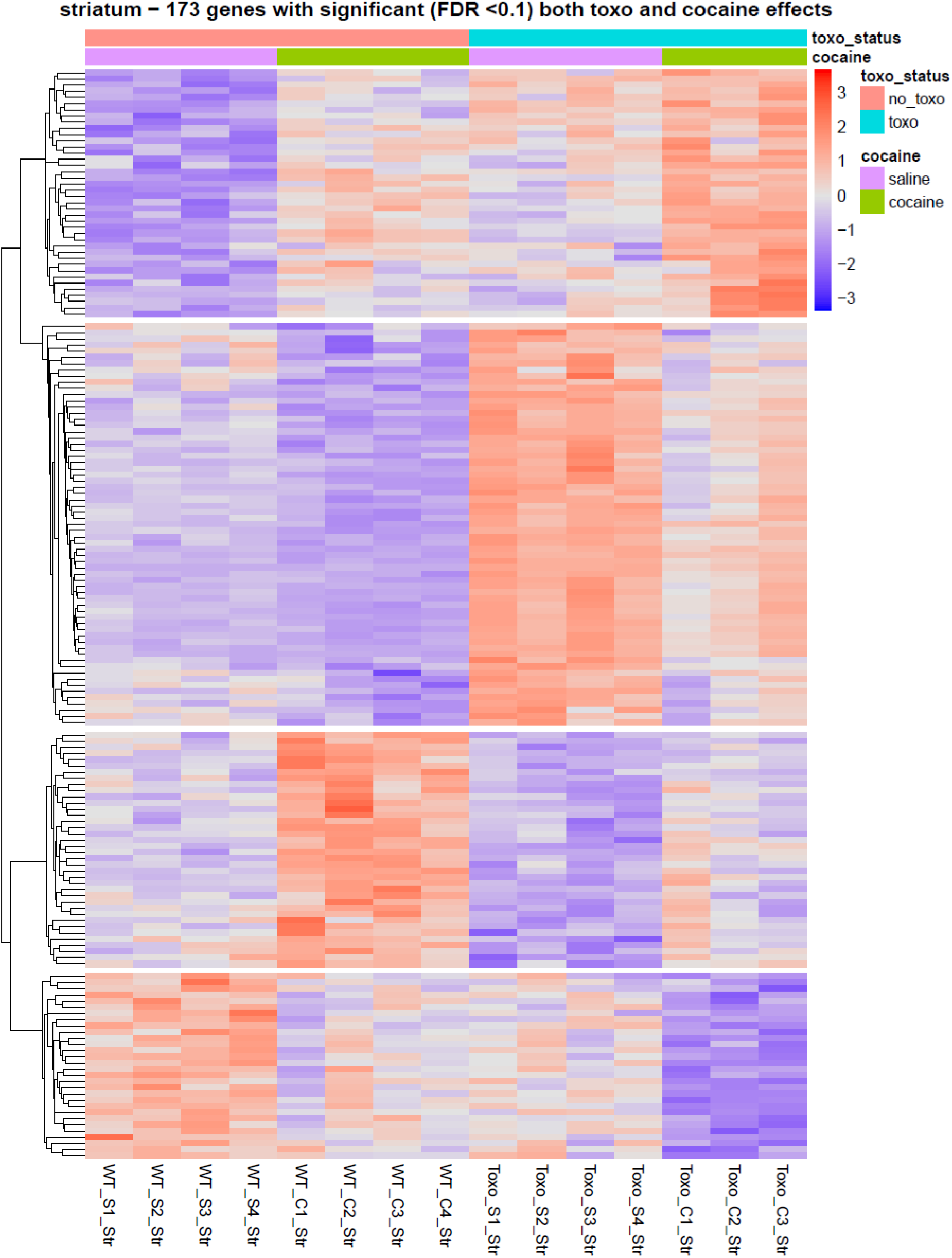
Heatmaps of gene expression defining DE gene responses by type and subtype.

**Suppl. Figure 12:**
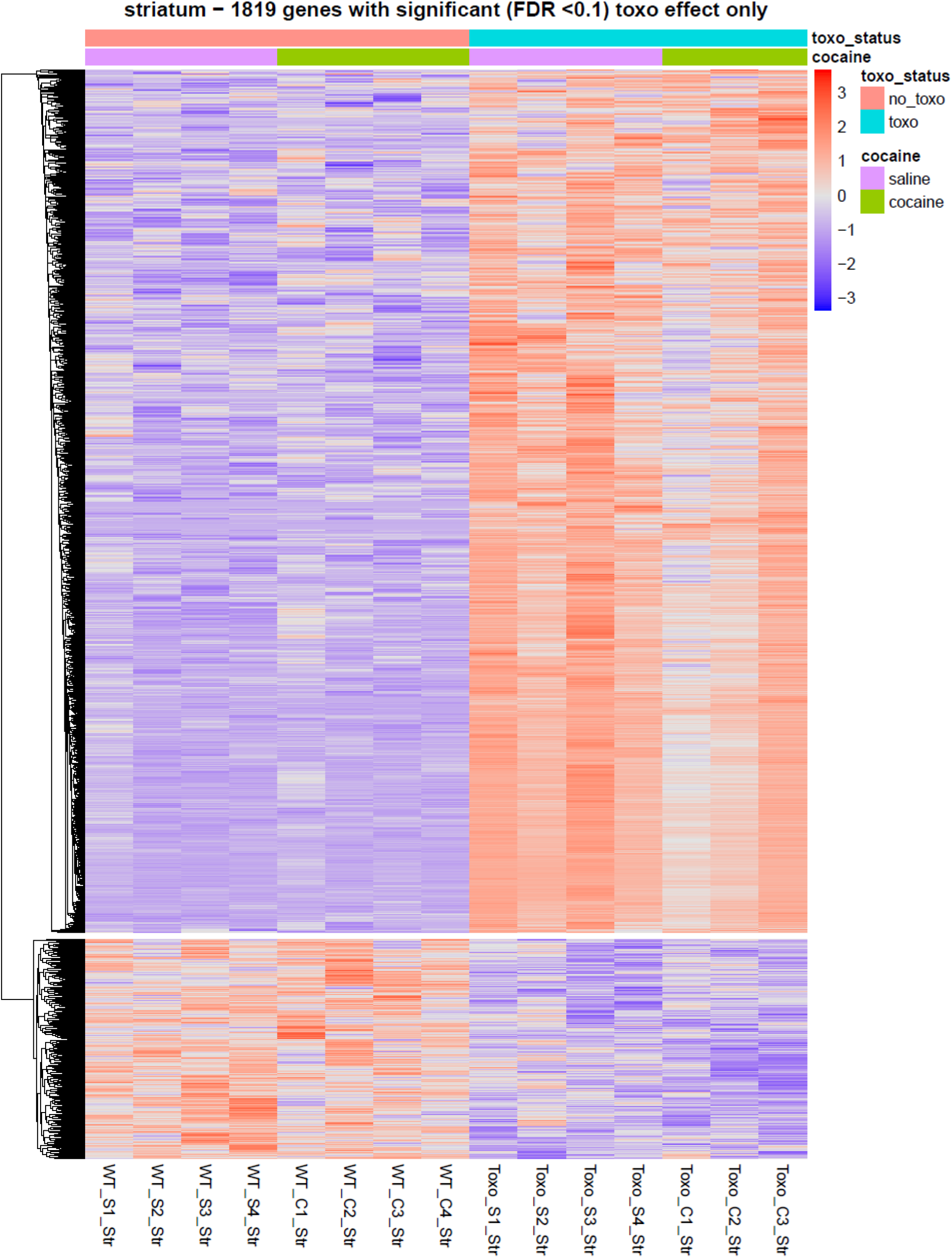
Heatmaps of gene expression defining DE gene responses by type and subtype.

**Suppl. Figure 13:**
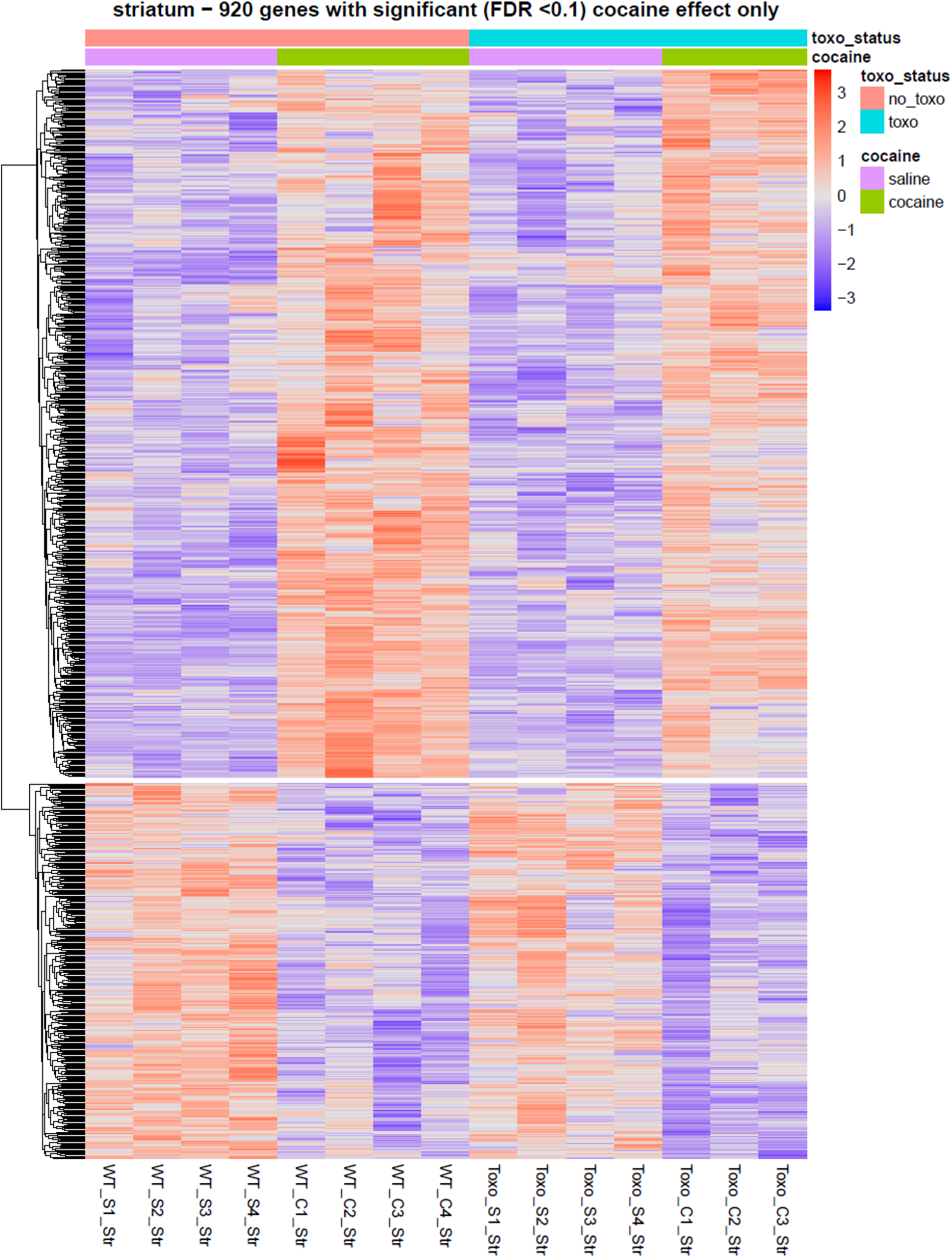
Heatmaps of gene expression defining DE gene responses by type and subtype.

## SUPPLEMENTARY FILES

S1. Supplementary spreadsheet (macro-enabled) containing colorized heatmap of significant, differentially expressed genes in mouse hippocampus and striatum with accompanying metadata.

S2. Supplementary spreadsheet detailing the statistical breakdown of DESeq2 significant gene sets in hippocampus.

S3. Supplementary spreadsheet detailing the statistical breakdown of DESeq2 significant gene sets in striatum.

S4. Supplementary spreadsheet with statistical analysis of genes in categories

S5. Zip archive containing functional enrichment details (i.e., BiNGO output).

S6. Zip archive of ontologies and annotations used in this study.

S7. Supplementary spreadsheet with curation of neurologically relevant functional terms.

S8. Supplementary spreadsheet with context enrichment analysis.

S9. Supplementary spreadsheet with analysis of significant CxT interactions.

S10. Cytoscape session file with functional annotations associated with context enriched genes.

