## Supplemental File 6 for "*Toxoplasma* infection alters dopamine-sensitive behaviors and host gene expression patterns associated with neuropsychiatric disease": reactome_ontology.txt

(curator=REACTOME)(type=pathway)
00001 = Reactome_Pathway
9612973 = Autophagy [isa: 00001]
1640170 = Cell Cycle [isa: 00001]
1500931 = Cell-Cell communication [isa: 00001]
8953897 = Cellular responses to external stimuli [isa: 00001]
4839726 = Chromatin organization [isa: 00001]
1266738 = Developmental Biology [isa: 00001]
8963743 = Digestion and absorption [isa: 00001]
1643685 = Disease [isa: 00001]
73894 = DNA Repair [isa: 00001]
69306 = DNA Replication [isa: 00001]
1474244 = Extracellular matrix organization [isa: 00001]
74160 = Gene expression (Transcription) [isa: 00001]
109582 = Hemostasis [isa: 00001]
168256 = Immune System [isa: 00001]
1430728 = Metabolism [isa: 00001]
392499 = Metabolism of proteins [isa: 00001]
8953854 = Metabolism of RNA [isa: 00001]
397014 = Muscle contraction [isa: 00001]
112316 = Neuronal System [isa: 00001]
1852241 = Organelle biogenesis and maintenance [isa: 00001]
5357801 = Programmed Cell Death [isa: 00001]
9609507 = Protein localization [isa: 00001]
1474165 = Reproduction [isa: 00001]
162582 = Signal Transduction [isa: 00001]
382551 = Transport of small molecules [isa: 00001]
5653656 = Vesicle-mediated transport [isa: 00001]
109581 = Apoptosis [isa: 5357801]
169911 = Regulation of Apoptosis [isa: 109581]
5357769 = Caspase activation via extrinsic apoptotic signalling pathway [isa: 109581]
75153 = Apoptotic execution phase [isa: 109581]
140877 = Formation of Fibrin Clot (Clotting Cascade) [isa: 109582]
202733 = Cell surface interactions at the vascular wall [isa: 109582]
418346 = Platelet homeostasis [isa: 109582]
75205 = Dissolution of Fibrin Clot [isa: 109582]
75892 = Platelet Adhesion to exposed collagen [isa: 109582]
76002 = Platelet activation, signaling and aggregation [isa: 109582]
983231 = Factors involved in megakaryocyte development and platelet production [isa: 109582]
109606 = Intrinsic Pathway for Apoptosis [isa: 109581]
111453 = BH3-only proteins associate with and inactivate anti-apoptotic BCL-2 members [isa: 109606]
111471 = Apoptotic factor-mediated response [isa: 109606]
114294 = Activation, translocation and oligomerization of BAX [isa: 109606]
114452 = Activation of BH3-only proteins [isa: 109606]
75108 = Activation, myristolyation of BID and translocation to mitochondria [isa: 109606]
9006934 = Signaling by Receptor Tyrosine Kinases [isa: 162582]
2404192 = Signaling by Type 1 Insulin-like Growth Factor 1 Receptor (IGF1R) [isa: 9006934]
2428924 = IGF1R signaling cascade [isa: 2404192]
73893 = DNA Damage Bypass [isa: 73894]
110313 = Translesion synthesis by Y family DNA polymerases bypasses lesions on DNA template [isa: 73893]
5655862 = Translesion synthesis by POLK [isa: 110313]
5656121 = Translesion synthesis by POLI [isa: 110313]
5656169 = Termination of translesion DNA synthesis [isa: 110313]
73884 = Base Excision Repair [isa: 73894]
73933 = Resolution of Abasic Sites (AP sites) [isa: 73884]
110381 = Resolution of AP sites via the single-nucleotide replacement pathway [isa: 73933]
111461 = Cytochrome c-mediated apoptotic response [isa: 111471]
111458 = Formation of apoptosome [isa: 111461]
111465 = Apoptotic cleavage of cellular proteins [isa: 75153]
351906 = Apoptotic cleavage of cell adhesion  proteins [isa: 111465]
352238 = Breakdown of the nuclear lamina [isa: 111465]
111469 = SMAC, XIAP-regulated apoptotic response [isa: 111471]
111464 = SMAC(DIABLO)-mediated dissociation of IAP:caspase complexes  [isa: 111469]
111457 = Release of apoptotic factors from the mitochondria [isa: 111471]
9627069 = Regulation of the apoptosome activity [isa: 111458]
111463 = SMAC (DIABLO) binds to IAPs  [isa: 111469]
372790 = Signaling by GPCR [isa: 162582]
388396 = GPCR downstream signalling [isa: 372790]
418594 = G alpha (i) signalling events [isa: 388396]
111885 = Opioid Signalling [isa: 418594]
112040 = G-protein mediated events [isa: 111885]
112043 = PLC beta mediated events [isa: 112040]
111996 = Ca-dependent events [isa: 112043]
9006925 = Intracellular signaling by second messengers [isa: 162582]
1489509 = DAG and IP3 signaling [isa: 9006925]
111997 = CaM pathway [isa: 111996 1489509]
111933 = Calmodulin induced events [isa: 111997]
170660 = Adenylate cyclase activating pathway [isa: 112040]
112315 = Transmission across Chemical Synapses [isa: 112316]
111995 = phospho-PLA2 pathway [isa: 111996]
112307 = Transmission across Electrical Synapses  [isa: 112316]
112314 = Neurotransmitter receptors and postsynaptic signal transmission [isa: 112315]
112310 = Neurotransmitter release cycle [isa: 112315]
210500 = Glutamate Neurotransmitter Release Cycle [isa: 112310]
212676 = Dopamine Neurotransmitter Release Cycle [isa: 112310]
264642 = Acetylcholine Neurotransmitter Release Cycle [isa: 112310]
888590 = GABA synthesis, release, reuptake and degradation [isa: 112310]
112311 = Neurotransmitter clearance [isa: 112315]
380615 = Serotonin clearance from the synaptic cleft [isa: 112311]
112313 = Neurotransmitter uptake and metabolism In glial cells [isa: 112315]
977443 = GABA receptor activation [isa: 112314]
399721 = Glutamate binding, activation of AMPA receptors and synaptic plasticity [isa: 112314]
442755 = Activation of NMDA receptors and postsynaptic events [isa: 112314]
451326 = Activation of kainate receptors upon glutamate binding [isa: 112314]
977444 = GABA B receptor activation [isa: 977443]
181430 = Norepinephrine Neurotransmitter Release Cycle [isa: 112310]
181429 = Serotonin Neurotransmitter Release Cycle [isa: 112310]
379401 = Dopamine clearance from the synaptic cleft [isa: 112311]
210455 = Astrocytic Glutamate-Glutamine Uptake And Metabolism [isa: 112313]
181431 = Acetylcholine binding and downstream events [isa: 112314]
112303 = Electric Transmission Across Gap Junctions [isa: 112307]
112308 = Presynaptic depolarization and calcium channel opening [isa: 112315]
1296071 = Potassium Channels [isa: 112316]
6794362 = Protein-protein interactions at synapses [isa: 112316]
2428928 = IRS-related events triggered by IGF1R [isa: 2428924]
74752 = Signaling by Insulin receptor [isa: 9006934]
5683057 = MAPK family signaling cascades [isa: 162582]
5684996 = MAPK1/MAPK3 signaling [isa: 5683057]
69278 = Cell Cycle, Mitotic [isa: 1640170]
111446 = Activation of BIM and translocation to mitochondria  [isa: 114452]
111447 = Activation of BAD and translocation to mitochondria  [isa: 114452]
111448 = Activation of NOXA and translocation to mitochondria [isa: 114452]
139910 = Activation of BMF and translocation to mitochondria [isa: 114452]
416476 = G alpha (q) signalling events [isa: 388396]
1280218 = Adaptive Immune System [isa: 168256]
983705 = Signaling by the B Cell Receptor (BCR) [isa: 1280218]
1168372 = Downstream signaling events of B Cell Receptor (BCR) [isa: 983705]
1280215 = Cytokine Signaling in Immune system [isa: 168256]
913531 = Interferon Signaling [isa: 1280215]
1187000 = Fertilization [isa: 1474165]
1300645 = Acrosome Reaction and Sperm:Oocyte Membrane Binding [isa: 1187000]
2534343 = Interaction With Cumulus Cells And The Zona Pellucida [isa: 1187000]
168249 = Innate Immune System [isa: 168256]
74751 = Insulin receptor signalling cascade [isa: 74752]
1227986 = Signaling by ERBB2 [isa: 9006934]
1963640 = GRB2 events in ERBB2 signaling [isa: 1227986]
1963642 = PI3K events in ERBB2 signaling [isa: 1227986]
6785631 = ERBB2 Regulates Cell Motility [isa: 1227986]
9006927 = Signaling by Non-Receptor Tyrosine Kinases [isa: 162582]
8863795 = Downregulation of ERBB2 signaling [isa: 1227986]
2262752 = Cellular responses to stress [isa: 8953897]
1234174 = Cellular response to hypoxia [isa: 2262752]
1236394 = Signaling by ERBB4 [isa: 9006934]
1250347 = SHC1 events in ERBB4 signaling [isa: 1236394]
1251985 = Nuclear signaling by ERBB4 [isa: 1236394]
1253288 = Downregulation of ERBB4 signaling [isa: 1236394]
983169 = Class I MHC mediated antigen processing & presentation [isa: 1280218]
1236975 = Antigen processing-Cross presentation [isa: 983169]
1236977 = Endosomal/Vacuolar pathway [isa: 1236975]
1236978 = Cross-presentation of soluble exogenous antigens (endosomes) [isa: 1236975]
111957 = Cam-PDE 1 activation [isa: 111933]
1257604 = PIP3 activates AKT signaling [isa: 9006925]
199418 = Negative regulation of the PI3K/AKT network [isa: 1257604]
6807070 = PTEN Regulation [isa: 1257604]
1181150 = Signaling by NODAL [isa: 1266738]
186712 = Regulation of beta-cell development [isa: 1266738]
381340 = Transcriptional regulation of white adipocyte differentiation [isa: 1266738]
422475 = Axon guidance [isa: 1266738]
525793 = Myogenesis [isa: 1266738]
5682910 = LGI-ADAM interactions [isa: 1266738]
6805567 = Keratinization [isa: 1266738]
9616222 = Transcriptional regulation of granulopoiesis [isa: 1266738]
1169410 = Antiviral mechanism by IFN-stimulated genes [isa: 913531]
449147 = Signaling by Interleukins [isa: 1280215]
5668541 = TNFR2 non-canonical NF-kB pathway [isa: 1280215]
8983711 = OAS antiviral response [isa: 1169410]
9607240 = FLT3 Signaling [isa: 1280215]
982772 = Growth hormone receptor signaling [isa: 1280215]
2025928 = Calcineurin activates NFAT [isa: 1168372]
202403 = TCR signaling [isa: 1280218]
2132295 = MHC class II antigen presentation [isa: 1280218]
388841 = Costimulation by the CD28 family [isa: 1280218]
392517 = Rap1 signalling [isa: 1280218]
8851680 = Butyrophilin (BTN) family interactions [isa: 1280218]
1236974 = ER-Phagosome pathway [isa: 1236975]
1169092 = Activation of RAS in B cells [isa: 1168372]
1296065 = Inwardly rectifying K+ channels [isa: 1296071]
1296059 = G protein gated Potassium channels [isa: 1296065]
1296053 = Classical Kir channels [isa: 1296065]
1296041 = Activation of G protein gated Potassium channels [isa: 1296059]
1296067 = Potassium transport channels [isa: 1296065]
1296052 = Ca2+ activated K+ channels [isa: 1296071]
1296061 = HCN channels [isa: 1296071]
1296025 = ATP sensitive Potassium channels [isa: 1296065]
1296072 = Voltage gated Potassium channels [isa: 1296071]
1296346 = Tandem pore domain potassium channels [isa: 1296071]
1299287 = Tandem pore domain halothane-inhibited K+ channel (THIK) [isa: 1296346]
1299308 = Tandem of pore domain in a weak inwardly rectifying K+ channels (TWIK) [isa: 1296346]
1299316 = TWIK-releated acid-sensitive K+ channel (TASK) [isa: 1296346]
1299344 = TWIK-related spinal cord K+ channel (TRESK) [isa: 1296346]
1299361 = TWIK-related alkaline pH activated K+ channel (TALK) [isa: 1296346]
1299503 = TWIK related potassium channel (TREK) [isa: 1296346]
1362409 = Mitochondrial iron-sulfur cluster biogenesis [isa: 1430728]
211859 = Biological oxidations [isa: 1430728]
211945 = Phase I - Functionalization of compounds [isa: 211859]
140534 = Caspase activation via Death Receptors in the presence of ligand [isa: 5357769]
69416 = Dimerization of procaspase-8 [isa: 140534]
140834 = Extrinsic Pathway of Fibrin Clot Formation [isa: 140877]
140837 = Intrinsic Pathway of Fibrin Clot Formation [isa: 140877]
140875 = Common Pathway of Fibrin Clot Formation [isa: 140877]
453276 = Regulation of mitotic cell cycle [isa: 69278]
69620 = Cell Cycle Checkpoints [isa: 1640170]
1428517 = The citric acid (TCA) cycle and respiratory electron transport [isa: 1430728]
71406 = Pyruvate metabolism and Citric Acid (TCA) cycle [isa: 1428517]
2395516 = Electron transport from NADPH to Ferredoxin [isa: 1362409]
163200 = Respiratory electron transport, ATP synthesis by chemiosmotic coupling, and heat production by uncoupling proteins. [isa: 1428517]
1475029 = Reversible hydration of carbon dioxide [isa: 1430728]
1483249 = Inositol phosphate metabolism [isa: 1430728]
15869 = Metabolism of nucleotides [isa: 1430728]
163685 = Integration of energy metabolism [isa: 1430728]
189445 = Metabolism of porphyrins [isa: 1430728]
196854 = Metabolism of vitamins and cofactors [isa: 1430728]
202131 = Metabolism of nitric oxide: eNOS activation and regulation [isa: 1430728]
140179 = Amine Oxidase reactions [isa: 211945]
2161522 = Abacavir transport and metabolism [isa: 1430728]
556833 = Metabolism of lipids [isa: 1430728]
71291 = Metabolism of amino acids and derivatives [isa: 1430728]
71387 = Metabolism of carbohydrates [isa: 1430728]
71737 = Pyrophosphate hydrolysis [isa: 1430728]
1433557 = Signaling by SCF-KIT [isa: 9006934]
6803157 = Antimicrobial peptides [isa: 168249]
1461973 = Defensins [isa: 6803157]
1300642 = Sperm Motility And Taxes [isa: 1187000]
1474228 = Degradation of the extracellular matrix [isa: 1474244]
1592389 = Activation of Matrix Metalloproteinases [isa: 1474228]
1442490 = Collagen degradation [isa: 1474228]
1474290 = Collagen formation [isa: 1474244]
1566948 = Elastic fibre formation [isa: 1474244]
1566977 = Fibronectin matrix formation [isa: 1474244]
216083 = Integrin cell surface interactions [isa: 1474244]
3000157 = Laminin interactions [isa: 1474244]
3000171 = Non-integrin membrane-ECM interactions [isa: 1474244]
3000178 = ECM proteoglycans [isa: 1474244]
8941237 = Invadopodia formation [isa: 1474244]
1650814 = Collagen biosynthesis and modifying enzymes [isa: 1474290]
2022090 = Assembly of collagen fibrils and other multimeric structures [isa: 1474290]
1480926 = O2/CO2 exchange in erythrocytes [isa: 382551]
1247673 = Erythrocytes take up oxygen and release carbon dioxide [isa: 1480926]
1483257 = Phospholipid metabolism [isa: 556833]
1483206 = Glycerophospholipid biosynthesis [isa: 1483257]
1482801 = Acyl chain remodelling of PS [isa: 1483206]
1482839 = Acyl chain remodelling of PE [isa: 1483206]
1482883 = Acyl chain remodeling of DAG and TAG [isa: 1483206]
1482922 = Acyl chain remodelling of PI [isa: 1483206]
1482925 = Acyl chain remodelling of PG [isa: 1483206]
1483076 = Synthesis of CL [isa: 1483206]
1483101 = Synthesis of PS [isa: 1483206]
1483115 = Hydrolysis of LPC [isa: 1483206]
1483148 = Synthesis of PG [isa: 1483206]
1483152 = Hydrolysis of LPE [isa: 1483206]
1483166 = Synthesis of PA [isa: 1483206]
1483191 = Synthesis of PC [isa: 1483206]
1483255 = PI Metabolism [isa: 1483257]
1483213 = Synthesis of PE [isa: 1483206]
1483226 = Synthesis of PI [isa: 1483206]
1855167 = Synthesis of pyrophosphates in the cytosol [isa: 1483249]
1855183 = Synthesis of IP2, IP, and Ins in the cytosol [isa: 1483249]
1855191 = Synthesis of IPs in the nucleus [isa: 1483249]
1855204 = Synthesis of IP3 and IP4 in the cytosol [isa: 1483249]
1855231 = Synthesis of IPs in the ER lumen [isa: 1483249]
1483196 = PI and PC transport between ER and Golgi membranes [isa: 1483206 1483255]
1483248 = Synthesis of PIPs at the ER membrane [isa: 1483255]
1660499 = Synthesis of PIPs at the plasma membrane [isa: 1483255]
1660514 = Synthesis of PIPs at the Golgi membrane [isa: 1483255]
1660516 = Synthesis of PIPs at the early endosome membrane [isa: 1483255]
1660517 = Synthesis of PIPs at the late endosome membrane [isa: 1483255]
6814848 = Glycerophospholipid catabolism [isa: 1483255]
8847453 = Synthesis of PIPs in the nucleus [isa: 1483255]
1482798 = Acyl chain remodeling of CL [isa: 1483206]
111931 = PKA-mediated phosphorylation of CREB [isa: 111933]
373753 = Nephrin family interactions [isa: 1500931]
391160 = Signal regulatory protein family interactions [isa: 1500931]
446728 = Cell junction organization [isa: 1500931]
9006936 = Signaling by TGF-beta family members [isa: 162582]
156580 = Phase II - Conjugation of compounds [isa: 211859]
156582 = Acetylation [isa: 156580]
156584 = Cytosolic sulfonation of small molecules [isa: 156580]
156587 = Amino Acid conjugation [isa: 156580]
156588 = Glucuronidation [isa: 156580]
156590 = Glutathione conjugation [isa: 156580]
1630316 = Glycosaminoglycan metabolism [isa: 71387]
159424 = Conjugation of carboxylic acids [isa: 156587]
173599 = Formation of the active cofactor, UDP-glucuronate [isa: 156588]
174403 = Glutathione synthesis and recycling [isa: 156590]
2129379 = Molecules associated with elastic fibres [isa: 1566948]
72766 = Translation [isa: 392499]
157118 = Signaling by NOTCH [isa: 162582]
1980143 = Signaling by NOTCH1 [isa: 157118]
9012852 = Signaling by NOTCH3 [isa: 157118]
9013694 = Signaling by NOTCH4 [isa: 157118]
73886 = Chromosome Maintenance [isa: 1640170]
157579 = Telomere Maintenance [isa: 73886]
199991 = Membrane Trafficking [isa: 5653656]
157858 = Gap junction trafficking and regulation [isa: 199991]
499943 = Interconversion of nucleotide di- and triphosphates [isa: 15869]
8956319 = Nucleobase catabolism [isa: 15869]
8956320 = Nucleobase biosynthesis [isa: 15869]
8956321 = Nucleotide salvage [isa: 15869]
1592230 = Mitochondrial biogenesis [isa: 1852241]
8949613 = Cristae formation [isa: 1592230]
72203 = Processing of Capped Intron-Containing Pre-mRNA [isa: 8953854]
72202 = Transport of Mature Transcript to Cytoplasm [isa: 72203]
159234 = Transport of Mature mRNAs Derived from Intronless Transcripts [isa: 72202]
177128 = Conjugation of salicylate with glycine [isa: 159424]
177135 = Conjugation of benzoate with glycine [isa: 159424]
177162 = Conjugation of phenylacetate with glutamine [isa: 159424]
597592 = Post-translational protein modification [isa: 392499]
163841 = Gamma carboxylation, hypusine formation and arylsulfatase activation [isa: 597592]
159854 = Gamma-carboxylation, transport, and amino-terminal cleavage of proteins [isa: 163841]
1834949 = Cytosolic sensors of pathogen-associated DNA  [isa: 168249]
1606322 = ZBP1(DAI) mediated induction of type I IFNs [isa: 1834949]
1614635 = Sulfur amino acid metabolism [isa: 71291]
1614558 = Degradation of cysteine and homocysteine [isa: 1614635]
1614517 = Sulfide oxidation to sulfate [isa: 1614558]
1614603 = Cysteine formation from homocysteine [isa: 1614635]
1912422 = Pre-NOTCH Expression and Processing [isa: 157118]
165159 = mTOR signalling [isa: 162582]
194315 = Signaling by Rho GTPases [isa: 162582]
195721 = Signaling by WNT [isa: 162582]
2028269 = Signaling by Hippo [isa: 162582]
2586552 = Signaling by Leptin [isa: 162582]
76009 = Platelet Aggregation (Plug Formation) [isa: 76002]
163615 = PKA activation [isa: 111931]
5358351 = Signaling by Hedgehog [isa: 162582]
112409 = RAF-independent MAPK1/3 activation [isa: 5684996]
73887 = Death Receptor Signalling [isa: 162582]
9006335 = Signaling by Erythropoietin [isa: 162582]
198693 = AKT phosphorylates targets in the nucleus [isa: 1257604]
8848021 = Signaling by PTK6 [isa: 9006927]
9006931 = Signaling by Nuclear Receptors [isa: 162582]
1306955 = GRB7 events in ERBB2 signaling [isa: 1227986]
1502540 = Signaling by Activin [isa: 9006936]
174362 = Transport and synthesis of PAPS [isa: 156584 1630316]
1638091 = Heparan sulfate/heparin (HS-GAG) metabolism [isa: 1630316]
1793185 = Chondroitin sulfate/dermatan sulfate metabolism [isa: 1630316]
2142845 = Hyaluronan metabolism [isa: 1630316]
159782 = Removal of aminoterminal propeptides from gamma-carboxylated proteins [isa: 159854]
163125 = Post-translational modification: synthesis of GPI-anchored proteins [isa: 597592]
162791 = Attachment of GPI anchor to uPAR [isa: 163125]
163210 = Formation of ATP by chemiosmotic coupling [isa: 163200]
166187 = Mitochondrial Uncoupling [isa: 163200]
611105 = Respiratory electron transport [isa: 163200]
1632852 = Macroautophagy [isa: 9612973]
163359 = Glucagon signaling in metabolic regulation [isa: 163685]
163358 = PKA-mediated phosphorylation of key metabolic factors [isa: 163685]
164378 = PKA activation in glucagon signalling [isa: 163359]
163680 = AMPK inhibits chREBP transcriptional activation activity [isa: 163685]
163754 = Insulin effects increased synthesis of Xylulose-5-Phosphate [isa: 163685]
163765 = ChREBP activates metabolic gene expression [isa: 163685]
422356 = Regulation of insulin secretion [isa: 163685]
1638074 = Keratan sulfate/keratin metabolism [isa: 1630316]
2022857 = Keratan sulfate degradation [isa: 1638074]
1971475 = A tetrasaccharide linker sequence is required for GAG synthesis [isa: 1638091 1793185]
2022928 = HS-GAG biosynthesis [isa: 1638091]
2024096 = HS-GAG degradation [isa: 1638091]
159763 = Transport of gamma-carboxylated protein precursors from the endoplasmic reticulum to the Golgi apparatus [isa: 159854]
1663150 = The activation of arylsulfatases [isa: 163841]
204626 = Hypusine synthesis from eIF5A-lysine [isa: 163841]
5358493 = Synthesis of diphthamide-EEF2 [isa: 163841]
453279 = Mitotic G1 phase and G1/S transition [isa: 69278]
69618 = Mitotic Spindle Checkpoint [isa: 69620]
180786 = Extension of Telomeres [isa: 157579]
8948216 = Collagen chain trimerization [isa: 1650814]
165181 = Inhibition of TSC complex formation by PKB [isa: 165159]
166208 = mTORC1-mediated signalling [isa: 165159]
380972 = Energy dependent regulation of mTOR by LKB1-AMPK [isa: 165159]
168898 = Toll-like Receptor Cascades [isa: 168249]
166016 = Toll Like Receptor 4 (TLR4) Cascade [isa: 168898]
166166 = MyD88-independent TLR4 cascade  [isa: 166016]
181438 = Toll Like Receptor 2 (TLR2) Cascade [isa: 168898]
168181 = Toll Like Receptor 7/8 (TLR7/8) Cascade [isa: 168898]
168176 = Toll Like Receptor 5 (TLR5) Cascade [isa: 168898]
168188 = Toll Like Receptor TLR6:TLR2 Cascade [isa: 181438]
448424 = Interleukin-17 signaling [isa: 449147]
167826 = The fatty acid cycling model [isa: 166187]
167827 = The proton buffering model [isa: 166187]
166520 = Signaling by NTRKs [isa: 9006934]
9006115 = Signaling by NTRK2 (TRKB) [isa: 166520]
9034015 = Signaling by NTRK3 (TRKC) [isa: 166520]
9036866 = Expression and Processing of Neurotrophins [isa: 166520]
166658 = Complement cascade [isa: 168249]
166665 = Terminal pathway of complement [isa: 166658]
174577 = Activation of C3 and C5 [isa: 166658]
977606 = Regulation of Complement cascade [isa: 166658]
166663 = Initial triggering of complement [isa: 166658]
166786 = Creation of C4 and C2 activators [isa: 166663]
173736 = Alternative complement activation [isa: 166663]
166662 = Lectin pathway of complement activation [isa: 166786]
173623 = Classical antibody-mediated complement activation [isa: 166786]
187037 = Signaling by NTRK1 (TRKA) [isa: 166520]
168138 = Toll Like Receptor 9 (TLR9) Cascade [isa: 168898]
168142 = Toll Like Receptor 10 (TLR10) Cascade [isa: 168898]
168164 = Toll Like Receptor 3 (TLR3) Cascade [isa: 168898]
9013973 = TICAM1-dependent activation of IRF3/IRF7 [isa: 168164]
975871 = MyD88 cascade initiated on plasma membrane [isa: 168142 168176]
168179 = Toll Like Receptor TLR1:TLR2 Cascade [isa: 181438]
975155 = MyD88 dependent cascade initiated on endosome [isa: 168138 168181]
166058 = MyD88:MAL(TIRAP) cascade initiated on plasma membrane [isa: 166016 168179 168188]
1222556 = ROS and RNS production in phagocytes [isa: 168249]
2855086 = Ficolins bind to repetitive carbohydrate structures on the target cell surface [isa: 166662]
168643 = Nucleotide-binding domain, leucine rich repeat containing receptor (NLR) signaling pathways [isa: 168249]
446652 = Interleukin-1 family signaling [isa: 449147]
168928 = DDX58/IFIH1-mediated induction of interferon-alpha/beta [isa: 168249]
1810476 = RIP-mediated NFkB activation via ZBP1 [isa: 1606322]
2029480 = Fcgamma receptor (FCGR) dependent phagocytosis [isa: 168249]
2172127 = DAP12 interactions [isa: 168249]
2454202 = Fc epsilon receptor (FCERI) signaling [isa: 168249]
5621481 = C-type lectin receptors (CLRs) [isa: 168249]
6798695 = Neutrophil degranulation [isa: 168249]
1462054 = Alpha-defensins [isa: 1461973]
879415 = Advanced glycosylation endproduct receptor signaling [isa: 168249]
9645460 = Alpha-protein kinase 1 signaling pathway [isa: 168249]
1170546 = Prolactin receptor signaling [isa: 1280215]
198933 = Immunoregulatory interactions between a Lymphoid and a non-Lymphoid cell [isa: 1280218]
8941413 = Events associated with phagocytolytic activity of PMN cells [isa: 1222556]
168638 = NOD1/2 Signaling Pathway [isa: 168643]
622312 = Inflammasomes [isa: 168643]
166020 = Transfer of LPS from LBP carrier to CD14 [isa: 166016]
1679131 = Trafficking and processing of endosomal TLR [isa: 168898]
937061 = TRIF(TICAM1)-mediated TLR4 signaling  [isa: 166166]
975138 = TRAF6 mediated induction of NFkB and MAP kinases upon TLR7/8 or 9 activation [isa: 975155]
937039 = IRAK1 recruits IKK complex [isa: 166058 975871]
5686938 = Regulation of TLR by endogenous ligand [isa: 168898]
918233 = TRAF3-dependent IRF activation pathway [isa: 168928]
933541 = TRAF6 mediated IRF7 activation [isa: 168928]
933542 = TRAF6 mediated NF-kB activation [isa: 168928]
936440 = Negative regulators of DDX58/IFIH1 signaling [isa: 168928]
187687 = Signalling to ERKs [isa: 187037]
169893 = Prolonged ERK activation events [isa: 187687]
170834 = Signaling by TGF-beta Receptor Complex [isa: 9006936]
2173791 = TGF-beta receptor signaling in EMT (epithelial to mesenchymal transition) [isa: 170834]
73857 = RNA Polymerase II Transcription [isa: 74160]
174143 = APC/C-mediated degradation of cell cycle proteins [isa: 453276]
174178 = APC/C:Cdh1 mediated degradation of Cdc20 and other APC/C:Cdh1 targeted proteins in late mitosis/early G1 [isa: 174143]
176407 = Conversion from APC/C:Cdc20 to APC/C:Cdh1 in late anaphase [isa: 174143]
176408 = Regulation of APC/C activators between G1/S and early anaphase [isa: 174143]
176814 = Activation of APC/C and APC/C:Cdc20 mediated degradation of mitotic proteins [isa: 174143]
174417 = Telomere C-strand (Lagging Strand) Synthesis [isa: 180786]
174414 = Processive synthesis on the C-strand of the telomere [isa: 174417]
174437 = Removal of the Flap Intermediate from the C-strand [isa: 174414]
174430 = Telomere C-strand synthesis initiation [isa: 174417]
174824 = Plasma lipoprotein assembly, remodeling, and clearance [isa: 382551]
8963899 = Plasma lipoprotein remodeling [isa: 174824]
8964043 = Plasma lipoprotein clearance [isa: 174824]
141405 = Inhibition of the proteolytic activity of APC/C required for the onset of anaphase by mitotic spindle checkpoint components [isa: 176408 69618]
176417 = Phosphorylation of Emi1 [isa: 176408]
176409 = APC/C:Cdc20 mediated degradation of mitotic proteins [isa: 176814]
174154 = APC/C:Cdc20 mediated degradation of Securin [isa: 176409]
179419 = APC:Cdc20 mediated degradation of cell cycle proteins prior to satisfation of the cell cycle checkpoint [isa: 176409]
174048 = APC/C:Cdc20 mediated degradation of Cyclin B [isa: 176409]
176412 = Phosphorylation of the APC/C [isa: 176814]
177929 = Signaling by EGFR [isa: 9006934]
180292 = GAB1 signalosome [isa: 177929]
180336 = SHC1 events in EGFR signaling [isa: 177929]
182971 = EGFR downregulation [isa: 177929]
212718 = EGFR interacts with phospholipase C-gamma [isa: 177929]
2022870 = Chondroitin sulfate biosynthesis [isa: 1793185]
2022923 = Dermatan sulfate biosynthesis [isa: 1793185]
2024101 = CS/DS degradation [isa: 1793185]
174184 = Cdc20:Phospho-APC/C mediated degradation of Cyclin A [isa: 179419]
179409 = APC-Cdc20 mediated degradation of Nek2A [isa: 179419]
171319 = Telomere Extension By Telomerase [isa: 180786]
174411 = Polymerase switching on the C-strand of the telomere [isa: 174417]
622323 = Presynaptic nicotinic acetylcholine receptors [isa: 181431]
622327 = Postsynaptic nicotinic acetylcholine receptors [isa: 181431]
1834941 = STING mediated induction of host immune responses [isa: 1834949]
3270619 = IRF3-mediated induction of type I IFN [isa: 1834941]
1606341 = IRF3 mediated activation of type 1 IFN [isa: 1606322]
3249367 = STAT6-mediated induction of chemokines [isa: 1834941]
3134963 = DEx/H-box helicases activate type I IFN and inflammatory cytokines production  [isa: 1834949]
3134973 = LRR FLII-interacting protein 1 (LRRFIP1) activates type I IFN production [isa: 1834949]
3134975 = Regulation of innate immune responses to cytosolic DNA [isa: 1834949]
2151201 = Transcriptional activation of mitochondrial biogenesis [isa: 1592230]
5617833 = Cilium Assembly [isa: 1852241]
210745 = Regulation of gene expression in beta cells [isa: 186712]
186797 = Signaling by PDGF [isa: 9006934]
170984 = ARMS-mediated activation [isa: 169893]
177504 = Retrograde neurotrophin signalling [isa: 187037]
187015 = Activation of TRKA receptors [isa: 187037]
167044 = Signalling to RAS [isa: 187687]
198203 = PI3K/AKT activation [isa: 187037]
198725 = Nuclear Events (kinase and transcription factor activation) [isa: 187037]
198765 = Signalling to ERK5 [isa: 187037]
171007 = p38MAPK events [isa: 167044]
170968 = Frs2-mediated activation [isa: 169893]
189451 = Heme biosynthesis [isa: 189445]
189483 = Heme degradation [isa: 189445]
190236 = Signaling by FGFR [isa: 9006934]
5654738 = Signaling by FGFR2 [isa: 190236]
5654741 = Signaling by FGFR3 [isa: 190236]
5654743 = Signaling by FGFR4 [isa: 190236]
190239 = FGFR3 ligand binding and activation [isa: 5654741]
190372 = FGFR3c ligand binding and activation [isa: 190239]
190241 = FGFR2 ligand binding and activation [isa: 5654738]
190377 = FGFR2b ligand binding and activation [isa: 190241]
5654736 = Signaling by FGFR1 [isa: 190236]
190242 = FGFR1 ligand binding and activation [isa: 5654736]
190374 = FGFR1c and Klotho ligand binding and activation [isa: 190242]
190322 = FGFR4 ligand binding and activation [isa: 5654743]
190828 = Gap junction trafficking [isa: 157858]
190873 = Gap junction degradation [isa: 190828]
190861 = Gap junction assembly [isa: 190828]
190872 = Transport of connexons to the plasma membrane [isa: 190861]
190840 = Microtubule-dependent trafficking of connexons from Golgi to the plasma membrane [isa: 190872]
196025 = Formation of annular gap junctions [isa: 190873]
1912408 = Pre-NOTCH Transcription and Translation [isa: 1912422]
1912420 = Pre-NOTCH Processing in Golgi [isa: 1912422]
8957322 = Metabolism of steroids [isa: 556833]
191273 = Cholesterol biosynthesis [isa: 8957322]
194068 = Bile acid and bile salt metabolism [isa: 8957322]
192105 = Synthesis of bile acids and bile salts [isa: 194068]
193807 = Synthesis of bile acids and bile salts via 27-hydroxycholesterol [isa: 192105]
193704 = p75 NTR receptor-mediated signalling [isa: 73887]
193639 = p75NTR signals via NF-kB [isa: 193704]
193697 = p75NTR regulates axonogenesis [isa: 193704]
209563 = Axonal growth stimulation [isa: 193697]
209560 = NF-kB is activated and signals survival [isa: 193639]
193692 = Regulated proteolysis of p75NTR [isa: 193704]
193634 = Axonal growth inhibition (RHOA activation) [isa: 193697]
204998 = Cell death signalling via NRAGE, NRIF and NADE [isa: 193704]
205017 = NFG and proNGF binds to p75NTR [isa: 193704]
193775 = Synthesis of bile acids and bile salts via 24-hydroxycholesterol [isa: 192105]
193368 = Synthesis of bile acids and bile salts via 7alpha-hydroxycholesterol [isa: 192105]
194138 = Signaling by VEGF [isa: 9006934]
194313 = VEGF ligand-receptor interactions [isa: 194138]
4420097 = VEGFA-VEGFR2 Pathway [isa: 194138]
195399 = VEGF binds to VEGFR leading to receptor dimerization [isa: 194313]
194840 = Rho GTPase cycle [isa: 194315]
195258 = RHO GTPase Effectors [isa: 194315]
194441 = Metabolism of non-coding RNA [isa: 8953854]
195253 = Degradation of beta-catenin by the destruction complex [isa: 195721]
4641265 = Repression of WNT target genes [isa: 195253]
5625740 = RHO GTPases activate PKNs [isa: 195258]
5625900 = RHO GTPases activate CIT [isa: 195258]
5625970 = RHO GTPases activate KTN1 [isa: 195258]
5626467 = RHO GTPases activate IQGAPs [isa: 195258]
5627083 = RHO GTPases regulate CFTR trafficking [isa: 195258]
5627117 = RHO GTPases Activate ROCKs [isa: 195258]
5627123 = RHO GTPases activate PAKs [isa: 195258]
5663213 = RHO GTPases Activate WASPs and WAVEs [isa: 195258]
5663220 = RHO GTPases Activate Formins [isa: 195258]
5666185 = RHO GTPases Activate Rhotekin and Rhophilins [isa: 195258]
5668599 = RHO GTPases Activate NADPH Oxidases [isa: 195258]
196299 = Beta-catenin phosphorylation cascade [isa: 195253]
201681 = TCF dependent signaling in response to WNT [isa: 195721]
3238698 = WNT ligand biogenesis and trafficking [isa: 195721]
3858494 = Beta-catenin independent WNT signaling [isa: 195721]
196071 = Metabolism of steroid hormones [isa: 8957322]
193144 = Estrogen biosynthesis [isa: 196071]
193993 = Mineralocorticoid biosynthesis [isa: 196071]
194002 = Glucocorticoid biosynthesis [isa: 196071]
196108 = Pregnenolone biosynthesis [isa: 196071]
196849 = Metabolism of water-soluble vitamins and cofactors [isa: 196854]
196807 = Nicotinate metabolism [isa: 196849]
196757 = Metabolism of folate and pterines [isa: 196849]
196780 = Biotin transport and metabolism [isa: 196849]
197264 = Nicotinamide salvaging [isa: 196807]
196819 = Vitamin B1 (thiamin) metabolism [isa: 196849]
196836 = Vitamin C (ascorbate) metabolism [isa: 196849]
196843 = Vitamin B2 (riboflavin) metabolism [isa: 196849]
199220 = Vitamin B5 (pantothenate) metabolism [isa: 196849]
947581 = Molybdenum cofactor biosynthesis [isa: 196849]
964975 = Vitamins B6 activation to pyridoxal phosphate [isa: 196849]
196741 = Cobalamin (Cbl, vitamin B12) transport and metabolism [isa: 196849]
6806667 = Metabolism of fat-soluble vitamins [isa: 196854]
8978934 = Metabolism of cofactors [isa: 196854]
2122947 = NOTCH1 Intracellular Domain Regulates Transcription [isa: 1980143]
2122948 = Activated NOTCH1 Transmits Signal to the Nucleus [isa: 1980143]
450294 = MAP kinase activation [isa: 166058 168164 448424 937061 975138 975871]
450282 = MAPK targets/ Nuclear events mediated by MAP kinases [isa: 450294]
198753 = ERK/MAPK targets [isa: 198725 450282]
196783 = Coenzyme A biosynthesis [isa: 199220]
6811558 = PI5P, PP2A and IER3 Regulate PI3K/AKT Signaling [isa: 199418]
446203 = Asparagine N-linked glycosylation [isa: 597592]
948021 = Transport to the Golgi and subsequent modification [isa: 446203]
199977 = ER to Golgi Anterograde Transport [isa: 199991 948021]
191650 = Regulation of gap junction activity [isa: 157858]
204005 = COPII-mediated vesicle transport [isa: 199977]
199992 = trans-Golgi Network Vesicle Budding [isa: 199991]
6811442 = Intra-Golgi and retrograde Golgi-to-ER traffic [isa: 199991]
8856828 = Clathrin-mediated endocytosis [isa: 199991]
9007101 = Rab regulation of trafficking [isa: 199991]
917729 = Endosomal Sorting Complex Required For Transport (ESCRT) [isa: 199991]
432720 = Lysosome Vesicle Biogenesis [isa: 199992]
432722 = Golgi Associated Vesicle Biogenesis [isa: 199992]
201688 = WNT mediated activation of DVL [isa: 201681]
201722 = Formation of the beta-catenin:TCF transactivating complex [isa: 201681]
3769402 = Deactivation of the beta-catenin transactivating complex [isa: 201681]
3772470 = Negative regulation of TCF-dependent signaling by WNT ligand antagonists [isa: 201681]
4641257 = Degradation of AXIN [isa: 201681]
4641258 = Degradation of DVL [isa: 201681]
4641262 = Disassembly of the destruction complex and recruitment of AXIN to the membrane [isa: 201681]
4641263 = Regulation of FZD by ubiquitination [isa: 201681]
203615 = eNOS activation [isa: 202131]
203641 = NOSTRIN mediated eNOS trafficking [isa: 202131]
203754 = NOSIP mediated eNOS trafficking [isa: 202131]
2214320 = Anchoring fibril formation [isa: 2022090]
2243919 = Crosslinking of collagen fibrils [isa: 2022090]
202424 = Downstream TCR signaling [isa: 202403]
202427 = Phosphorylation of CD3 and TCR zeta chains [isa: 202403]
202430 = Translocation of ZAP-70 to Immunological synapse [isa: 202403]
202433 = Generation of second messenger molecules [isa: 202403]
210990 = PECAM1 interactions [isa: 202733]
210991 = Basigin interactions [isa: 202733]
210993 = Tie2 Signaling [isa: 202733]
2029481 = FCGR activation [isa: 2029480]
2029482 = Regulation of actin dynamics for phagocytic cup formation [isa: 2029480]
2029485 = Role of phospholipids in phagocytosis [isa: 2029480]
8978868 = Fatty acid metabolism [isa: 556833]
2046104 = alpha-linolenic (omega3) and linoleic (omega6) acid metabolism [isa: 8978868]
193648 = NRAGE signals death through JNK [isa: 204998]
205025 = NADE modulates death signalling [isa: 204998]
205043 = NRIF signals cell death from the nucleus [isa: 204998]
209776 = Metabolism of amine-derived hormones [isa: 71291]
209931 = Serotonin and melatonin biosynthesis [isa: 209776]
209968 = Thyroxine biosynthesis [isa: 209776]
2980736 = Peptide hormone metabolism [isa: 392499]
350864 = Regulation of thyroid hormone activity [isa: 209968]
211163 = AKT-mediated inactivation of FOXO1A [isa: 210745]
211000 = Gene Silencing by RNA [isa: 74160]
426486 = Small interfering RNA (siRNA) biogenesis [isa: 211000]
426496 = Post-transcriptional silencing by small RNAs [isa: 211000]
5578749 = Transcriptional regulation by small RNAs [isa: 211000]
156581 = Methylation [isa: 156580]
141334 = PAOs oxidise polyamines to amines [isa: 140179]
5423646 = Aflatoxin activation and detoxification [isa: 211859]
211897 = Cytochrome P450 - arranged by substrate type [isa: 211945]
211935 = Fatty acids [isa: 211897]
211958 = Miscellaneous substrates [isa: 211897]
211976 = Endogenous sterols [isa: 211897]
211979 = Eicosanoids [isa: 211897]
211981 = Xenobiotics [isa: 211897]
141333 = Biogenic amines are oxidatively deaminated to aldehydes by MAOA and MAOB [isa: 140179]
140180 = COX reactions [isa: 211945]
211916 = Vitamins [isa: 211897]
217271 = FMO oxidises nucleophiles [isa: 211945]
71384 = Ethanol oxidation [isa: 211945]
8937144 = Aryl hydrocarbon receptor signalling [isa: 211945]
211994 = Sterols are 12-hydroxylated by CYP8B1 [isa: 211976]
211999 = CYP2E1 reactions [isa: 211981]
212165 = Epigenetic regulation of gene expression [isa: 74160]
5221030 = TET1,2,3 and TDG demethylate DNA [isa: 212165]
5250913 = Positive epigenetic regulation of rRNA expression [isa: 212165]
5250941 = Negative epigenetic regulation of rRNA expression [isa: 212165]
212436 = Generic Transcription Pathway [isa: 73857]
2173793 = Transcriptional activity of SMAD2/SMAD3:SMAD4 heterotrimer [isa: 170834 212436]
350054 = Notch-HLH transcription pathway [isa: 212436]
3700989 = Transcriptional Regulation by TP53 [isa: 212436]
383280 = Nuclear Receptor transcription pathway [isa: 212436]
8853884 = Transcriptional Regulation by VENTX [isa: 212436]
8864260 = Transcriptional regulation by the AP-2 (TFAP2) family of transcription factors [isa: 212436]
8878159 = Transcriptional regulation by RUNX3 [isa: 212436]
8878166 = Transcriptional regulation by RUNX2 [isa: 212436]
8878171 = Transcriptional regulation by RUNX1 [isa: 212436]
8953750 = Transcriptional Regulation by E2F6 [isa: 212436]
8986944 = Transcriptional Regulation by MECP2 [isa: 212436]
9614085 = FOXO-mediated transcription [isa: 212436]
2142753 = Arachidonic acid metabolism [isa: 8978868]
2142688 = Synthesis of 5-eicosatetraenoic acids [isa: 2142753]
2142691 = Synthesis of Leukotrienes (LT) and Eoxins (EX) [isa: 2142753]
2142696 = Synthesis of Hepoxilins (HX) and Trioxilins (TrX) [isa: 2142753]
2142712 = Synthesis of 12-eicosatetraenoic acid derivatives [isa: 2142753]
2142770 = Synthesis of 15-eicosatetraenoic acid derivatives [isa: 2142753]
2142816 = Synthesis of (16-20)-hydroxyeicosatetraenoic acids (HETE) [isa: 2142753]
2162123 = Synthesis of Prostaglandins (PG) and Thromboxanes (TX) [isa: 2142753]
2142850 = Hyaluronan biosynthesis and export [isa: 2142845]
2160916 = Hyaluronan uptake and degradation [isa: 2142845]
2161517 = Abacavir transmembrane transport [isa: 2161522]
2161541 = Abacavir metabolism [isa: 2161522]
2424491 = DAP12 signaling [isa: 2172127]
2173782 = Binding and Uptake of Ligands by Scavenger Receptors [isa: 5653656]
3000471 = Scavenging by Class B Receptors [isa: 2173782]
3000480 = Scavenging by Class A Receptors [isa: 2173782]
3000497 = Scavenging by Class H Receptors [isa: 2173782]
2173789 = TGF-beta receptor signaling activates SMADs [isa: 170834]
2173795 = Downregulation of SMAD2/3:SMAD4 transcriptional activity [isa: 2173793]
2173796 = SMAD2/SMAD3:SMAD4 heterotrimer regulates transcription [isa: 2173793]
2187338 = Visual phototransduction [isa: 418594]
2453902 = The canonical retinoid cycle in rods (twilight vision) [isa: 2187338]
2514856 = The phototransduction cascade [isa: 2187338]
975634 = Retinoid metabolism and transport [isa: 2187338 6806667]
1234176 = Oxygen-dependent proline hydroxylation of Hypoxia-inducible Factor Alpha [isa: 1234174]
2559583 = Cellular Senescence [isa: 2262752]
3299685 = Detoxification of Reactive Oxygen Species [isa: 2262752]
3371497 = HSP90 chaperone cycle for steroid hormone receptors (SHR) [isa: 2262752]
3371556 = Cellular response to heat stress [isa: 2262752]
9639288 = Amino acids regulate mTORC1 [isa: 2262752]
112399 = IRS-mediated signalling [isa: 2428928 74751]
2408522 = Selenoamino acid metabolism [isa: 71291]
2408557 = Selenocysteine synthesis [isa: 2408522]
5263617 = Metabolism of ingested MeSeO2H into MeSeH [isa: 2408522]
109704 = PI3K Cascade [isa: 112399 9607240]
2428933 = SHC-related events triggered by IGF1R [isa: 2428924]
109703 = PKB-mediated events [isa: 109704]
2730905 = Role of LAT2/NTAL/LAB on calcium mobilization [isa: 2454202]
2871796 = FCERI mediated MAPK activation [isa: 2454202]
2871809 = FCERI mediated Ca+2 mobilization [isa: 2454202]
2871837 = FCERI mediated NF-kB activation [isa: 2454202]
2485179 = Activation of the phototransduction cascade [isa: 2514856]
2514859 = Inactivation, recovery and regulation of the phototransduction cascade [isa: 2514856]
68886 = M Phase [isa: 69278]
2555396 = Mitotic Metaphase and Anaphase [isa: 68886]
2559580 = Oxidative Stress Induced Senescence [isa: 2559583]
2559582 = Senescence-Associated Secretory Phenotype (SASP) [isa: 2559583]
2559585 = Oncogene Induced Senescence [isa: 2559583]
2559586 = DNA Damage/Telomere Stress Induced Senescence [isa: 2559583]
2559584 = Formation of Senescence-Associated Heterochromatin Foci (SAHF) [isa: 2559586]
983712 = Ion channel transport [isa: 382551]
2682334 = EPH-Ephrin signaling [isa: 422475]
3928663 = EPHA-mediated growth cone collapse [isa: 2682334]
3928664 = Ephrin signaling [isa: 2682334]
3928665 = EPH-ephrin mediated repulsion of cells [isa: 2682334]
209952 = Peptide hormone biosynthesis [isa: 2980736]
209822 = Glycoprotein hormones [isa: 209952]
264876 = Insulin processing [isa: 2980736]
400508 = Incretin synthesis, secretion, and inactivation [isa: 2980736]
422085 = Synthesis, secretion, and deacylation of Ghrelin [isa: 2980736]
68875 = Mitotic Prophase [isa: 68886]
2980766 = Nuclear Envelope Breakdown [isa: 68875]
4419969 = Depolymerisation of the Nuclear Lamina [isa: 2980766]
2990846 = SUMOylation [isa: 597592]
3215018 = Processing and activation of SUMO [isa: 2990846]
68882 = Mitotic Anaphase [isa: 2555396]
3000170 = Syndecan interactions [isa: 3000171]
3108232 = SUMO E3 ligases SUMOylate target proteins [isa: 2990846]
3232118 = SUMOylation of transcription factors [isa: 3108232]
3232142 = SUMOylation of ubiquitinylation proteins [isa: 3108232]
3899300 = SUMOylation of transcription cofactors [isa: 3108232]
4085377 = SUMOylation of SUMOylation proteins [isa: 3108232]
4090294 = SUMOylation of intracellular receptors [isa: 3108232]
4551638 = SUMOylation of chromatin organization proteins [isa: 3108232]
4570464 = SUMOylation of RNA binding proteins [isa: 3108232]
4615885 = SUMOylation of DNA replication proteins [isa: 3108232]
4655427 = SUMOylation of DNA methylation proteins [isa: 3108232]
4755510 = SUMOylation of immune response proteins [isa: 3108232]
3065676 = SUMO is conjugated to E1 (UBA2:SAE1) [isa: 3215018]
3065678 = SUMO is transferred from E1 to E2 (UBE2I, UBC9) [isa: 3215018]
3065679 = SUMO is proteolytically processed [isa: 3215018]
3247509 = Chromatin modifying enzymes [isa: 4839726]
3214842 = HDMs demethylate histones [isa: 3247509]
3214847 = HATs acetylate histones [isa: 3247509]
3214858 = RMTs methylate histone arginines [isa: 3247509]
3371453 = Regulation of HSF1-mediated heat shock response [isa: 3371556]
3371511 = HSF1 activation [isa: 3371556]
3371571 = HSF1-dependent transactivation [isa: 3371556]
3371568 = Attenuation phase [isa: 3371571]
351202 = Metabolism of polyamines [isa: 71291]
351143 = Agmatine biosynthesis [isa: 351202]
351200 = Interconversion of polyamines [isa: 351202]
354192 = Integrin signaling [isa: 162582 76009]
372708 = p130Cas linkage to MAPK signaling for integrins [isa: 354192]
5628897 = TP53 Regulates Metabolic Genes [isa: 3700989]
5633007 = Regulation of TP53 Activity [isa: 3700989]
5633008 = TP53 Regulates Transcription of Cell Death Genes [isa: 3700989]
6791312 = TP53 Regulates Transcription of Cell Cycle Genes [isa: 3700989]
6796648 = TP53 Regulates Transcription of DNA Repair Genes [isa: 3700989]
202040 = G-protein activation [isa: 111885]
500792 = GPCR ligand binding [isa: 372790]
373076 = Class A/1 (Rhodopsin-like receptors) [isa: 500792]
375276 = Peptide ligand-binding receptors [isa: 373076]
375280 = Amine ligand-binding receptors [isa: 373076]
375281 = Hormone ligand-binding receptors [isa: 373076]
391903 = Eicosanoid ligand-binding receptors [isa: 373076]
418038 = Nucleotide-like (purinergic) receptors [isa: 373076]
419408 = Lysosphingolipid and LPA receptors [isa: 373076]
419771 = Opsins [isa: 373076]
444209 = Free fatty acid receptors [isa: 373076]
373080 = Class B/2 (Secretin family receptors) [isa: 500792]
420092 = Glucagon-type ligand receptors [isa: 373080]
373752 = Netrin-1 signaling [isa: 422475]
418885 = DCC mediated attractive signaling [isa: 373752]
373755 = Semaphorin interactions [isa: 422475]
399955 = SEMA3A-Plexin repulsion signaling by inhibiting Integrin adhesion [isa: 373755]
399956 = CRMPs in Sema3A signaling [isa: 373755]
400685 = Sema4D in semaphorin signaling [isa: 373755]
416700 = Other semaphorin interactions [isa: 373755]
373760 = L1CAM interactions [isa: 422475]
445095 = Interaction between L1 and Ankyrins [isa: 373760]
445144 = Signal transduction by L1 [isa: 373760]
447043 = Neurofascin interactions [isa: 373760]
375165 = NCAM signaling for neurite out-growth [isa: 422475]
380095 = Tachykinin receptors bind tachykinins [isa: 375276]
380108 = Chemokine receptors bind chemokines [isa: 375276]
388479 = Vasopressin-like receptors [isa: 375276]
389397 = Orexin and neuropeptides FF and QRFP bind to their respective receptors [isa: 375276]
444473 = Formyl peptide receptors bind formyl peptides and many other ligands [isa: 375276]
444821 = Relaxin receptors [isa: 375276]
390648 = Muscarinic acetylcholine receptors [isa: 375280]
390650 = Histamine receptors [isa: 375280]
390651 = Dopamine receptors [isa: 375280]
390666 = Serotonin receptors [isa: 375280]
390696 = Adrenoceptors [isa: 375280]
376176 = Signaling by ROBO receptors [isa: 422475]
8985586 = SLIT2:ROBO1 increases RHOA activity [isa: 376176]
9010553 = Regulation of expression of SLITs and ROBOs [isa: 376176]
379397 = Enzymatic degradation of dopamine by COMT [isa: 379401]
379398 = Enzymatic degradation of Dopamine by monoamine oxidase [isa: 379401]
379724 = tRNA Aminoacylation [isa: 72766]
379726 = Mitochondrial tRNA aminoacylation [isa: 379724]
453274 = Mitotic G2-G2/M phases [isa: 69278]
69275 = G2/M Transition [isa: 453274]
380287 = Centrosome maturation [isa: 69275]
380612 = Metabolism of serotonin [isa: 380615]
381119 = Unfolded Protein Response (UPR) [isa: 392499]
381042 = PERK regulates gene expression [isa: 381119]
381070 = IRE1alpha activates chaperones [isa: 381119]
8874211 = CREB3 factors activate genes [isa: 381119]
1237044 = Erythrocytes take up carbon dioxide and release oxygen [isa: 1480926]
8963898 = Plasma lipoprotein assembly [isa: 174824]
382556 = ABC-family proteins mediated transport [isa: 382551]
425407 = SLC-mediated transmembrane transport [isa: 382551]
445717 = Aquaporin-mediated transport [isa: 382551]
5223345 = Miscellaneous transport and binding events [isa: 382551]
8949215 = Mitochondrial calcium ion transport [isa: 382551]
8981607 = Intracellular oxygen transport [isa: 382551]
917937 = Iron uptake and transport [isa: 382551]
2672351 = Stimuli-sensing channels [isa: 983712]
1369007 = Mitochondrial ABC transporters [isa: 382556]
1369062 = ABC transporters in lipid homeostasis [isa: 382556]
4086398 = Ca2+ pathway [isa: 3858494]
4086400 = PCP/CE pathway [isa: 3858494]
397795 = G-protein beta:gamma signalling [isa: 388396]
114508 = Effects of PIP2 hydrolysis [isa: 416476 76002]
416482 = G alpha (12/13) signalling events [isa: 388396]
418555 = G alpha (s) signalling events [isa: 388396]
180024 = DARPP-32 events [isa: 111885]
418597 = G alpha (z) signalling events [isa: 388396]
389356 = CD28 co-stimulation [isa: 388841]
389513 = CTLA4 inhibitory signaling [isa: 388841]
389948 = PD-1 signaling [isa: 388841]
389357 = CD28 dependent PI3K/Akt signaling [isa: 389356]
389359 = CD28 dependent Vav1 pathway [isa: 389356]
389661 = Glyoxylate metabolism and glycine degradation [isa: 71291]
391251 = Protein folding [isa: 392499]
390466 = Chaperonin-mediated protein folding [isa: 391251]
390918 = Peroxisomal lipid metabolism [isa: 8978868]
389887 = Beta-oxidation of pristanoyl-CoA [isa: 390918]
390247 = Beta-oxidation of very long chain fatty acids [isa: 390918]
6814122 = Cooperation of PDCL (PhLP1) and TRiC/CCT in G-protein beta folding [isa: 390466]
391906 = Leukotriene receptors [isa: 391903]
391908 = Prostanoid ligand receptors [isa: 391903]
392154 = Nitric oxide stimulates guanylate cyclase [isa: 418346]
2022377 = Metabolism of Angiotensinogen to Angiotensins [isa: 2980736]
381033 = ATF6 (ATF6-alpha) activates chaperones [isa: 381119]
381426 = Regulation of Insulin-like Growth Factor (IGF) transport and uptake by Insulin-like Growth Factor Binding Proteins (IGFBPs) [isa: 392499]
390471 = Association of TriC/CCT with target proteins during biosynthesis [isa: 390466]
5676934 = Protein repair [isa: 392499]
5683826 = Surfactant metabolism [isa: 392499]
162710 = Synthesis of glycosylphosphatidylinositol (GPI) [isa: 163125]
156842 = Eukaryotic Translation Elongation [isa: 72766]
392518 = Signal amplification [isa: 76002]
418592 = ADP signalling through P2Y purinoceptor 1 [isa: 392518]
428930 = Thromboxane signalling through TP receptor [isa: 392518]
390522 = Striated Muscle Contraction [isa: 397014]
445355 = Smooth Muscle Contraction [isa: 397014]
5576891 = Cardiac conduction [isa: 397014]
392451 = G beta:gamma signalling through PI3Kgamma [isa: 397795]
418217 = G beta:gamma signalling through PLC beta [isa: 397795]
8964315 = G beta:gamma signalling through BTK [isa: 397795]
8964616 = G beta:gamma signalling through CDC42 [isa: 397795]
399719 = Trafficking of AMPA receptors [isa: 399721]
399710 = Activation of AMPA receptors [isa: 399721]
416993 = Trafficking of GluR2-containing AMPA receptors [isa: 399719]
400451 = Free fatty acids regulate insulin secretion [isa: 422356]
434316 = Fatty Acids bound to GPR40 (FFAR1) regulate insulin secretion [isa: 400451]
381771 = Synthesis, secretion, and inactivation of Glucagon-like Peptide-1 (GLP-1) [isa: 400508]
400511 = Synthesis, secretion, and inactivation of Glucose-dependent Insulinotropic Polypeptide (GIP) [isa: 400508]
416550 = Sema4D mediated inhibition of cell attachment and migration [isa: 400685]
416572 = Sema4D induced cell migration and growth-cone collapse [isa: 400685]
4608870 = Asymmetric localization of PCP proteins [isa: 4086400]
5099900 = WNT5A-dependent internalization of FZD4 [isa: 4086400]
5140745 = WNT5A-dependent internalization of FZD2, FZD5 and ROR2 [isa: 4086400]
426048 = Arachidonate production from DAG [isa: 114508]
881907 = Gastrin-CREB signalling pathway via PKC and MAPK [isa: 416476]
417957 = P2Y receptors [isa: 418038]
417973 = Adenosine P1 receptors [isa: 418038]
418457 = cGMP effects [isa: 392154]
392851 = Prostacyclin signalling through prostacyclin receptor [isa: 418346]
418360 = Platelet calcium homeostasis [isa: 418346]
432142 = Platelet sensitization by LDL [isa: 418346]
139853 = Elevation of cytosolic Ca2+ levels [isa: 418360]
418359 = Reduction of cytosolic Ca++ levels [isa: 418360]
381753 = Olfactory Signaling Pathway [isa: 418555]
111932 = CaMK IV-mediated phosphorylation of CREB [isa: 111933]
2187335 = The retinoid cycle in cones (daylight vision) [isa: 2187338]
421270 = Cell-cell junction organization [isa: 446728]
418990 = Adherens junctions interactions [isa: 421270]
420597 = Nectin/Necl  trans heterodimerization [isa: 418990]
420029 = Tight junction interactions [isa: 421270]
381676 = Glucagon-like Peptide-1 (GLP1) regulates insulin secretion [isa: 422356]
399997 = Acetylcholine regulates insulin secretion [isa: 422356]
400042 = Adrenaline,noradrenaline inhibits insulin secretion [isa: 422356]
434313 = Intracellular metabolism of fatty acids regulates insulin secretion [isa: 400451]
3928662 = EPHB-mediated forward signaling [isa: 2682334]
376172 = DSCAM interactions [isa: 373752]
399954 = Sema3A PAK dependent Axon repulsion [isa: 373755]
437239 = Recycling pathway of L1 [isa: 373760]
419037 = NCAM1 interactions [isa: 375165]
428890 = Role of ABL in ROBO-SLIT signaling [isa: 376176]
8853659 = RET signaling [isa: 422475]
8866376 = Reelin signalling pathway [isa: 422475]
425366 = Transport of bile salts and organic acids, metal ions and amine compounds [isa: 425407]
429593 = Inositol transporters [isa: 425366]
433137 = Sodium-coupled sulphate, di- and tri-carboxylate transporters [isa: 425366]
433692 = Proton-coupled monocarboxylate transport [isa: 425366]
442660 = Na+/Cl- dependent neurotransmitter transporters [isa: 425366]
444411 = Rhesus glycoproteins mediate ammonium transport. [isa: 425366]
549132 = Organic cation/anion/zwitterion transport [isa: 425366]
425393 = Transport of inorganic cations/anions and amino acids/oligopeptides [isa: 425407]
425381 = Bicarbonate transporters [isa: 425393]
425561 = Sodium/Calcium exchangers [isa: 425393]
425986 = Sodium/Proton exchangers [isa: 425393]
426117 = Cation-coupled Chloride cotransporters [isa: 425393]
427601 = Multifunctional anion exchangers [isa: 425393]
427652 = Sodium-coupled phosphate cotransporters [isa: 425393]
427975 = Proton/oligopeptide cotransporters [isa: 425393]
428559 = Proton-coupled neutral amino acid transporters [isa: 425393]
428643 = Organic anion transporters [isa: 425393]
425397 = Transport of vitamins, nucleosides, and related molecules [isa: 425407]
804914 = Transport of fatty acids [isa: 425397]
83936 = Transport of nucleosides and free purine and pyrimidine bases across the plasma membrane [isa: 425397]
879518 = Transport of organic anions [isa: 425397]
189200 = Cellular hexose transport [isa: 425407]
425410 = Metal ion SLC transporters [isa: 425366]
352230 = Amino acid transport across the plasma membrane [isa: 425393]
727802 = Transport of nucleotide sugars [isa: 425397]
435354 = Zinc transporters [isa: 425410]
427589 = Type II Na+/Pi cotransporters [isa: 427652]
428157 = Sphingolipid metabolism [isa: 556833]
1660662 = Glycosphingolipid metabolism [isa: 428157]
429914 = Deadenylation-dependent mRNA decay [isa: 8953854]
429958 = mRNA decay by 3' to 5' exoribonuclease [isa: 429914]
430039 = mRNA decay by 5' to 3' exoribonuclease [isa: 429914]
435368 = Zinc efflux and compartmentalization by the SLC30 family [isa: 435354]
442380 = Zinc influx into cells by the SLC39 gene family [isa: 435354]
438064 = Post NMDA receptor activation events [isa: 442755]
442729 = CREB1 phosphorylation through the activation of CaMKII/CaMKK/CaMKIV cascasde [isa: 438064]
442742 = CREB1 phosphorylation through NMDA receptor-mediated activation of RAS signaling [isa: 438064]
9619229 = Activation of RAC1 downstream of NMDARs [isa: 438064]
9620244 = Long-term potentiation [isa: 438064]
5218920 = VEGFR2 mediated vascular permeability [isa: 4420097]
5218921 = VEGFR2 mediated cell proliferation [isa: 4420097]
444257 = RSK activation [isa: 442742]
442720 = CREB1 phosphorylation through the activation of Adenylate Cyclase [isa: 438064]
438066 = Unblocking of NMDA receptors, glutamate binding and activation [isa: 442755]
9609736 = Assembly and cell surface presentation of NMDA receptors [isa: 442755]
432030 = Transport of glycerol from adipocytes to the liver by Aquaporins [isa: 445717]
432040 = Vasopressin regulates renal water homeostasis via Aquaporins [isa: 445717]
432047 = Passive transport by Aquaporins [isa: 445717]
6807878 = COPI-mediated anterograde transport [isa: 199977]
446193 = Biosynthesis of the N-glycan precursor (dolichol lipid-linked oligosaccharide, LLO) and transfer to a nascent protein [isa: 446203]
532668 = N-glycan trimming in the ER and Calnexin/Calreticulin cycle [isa: 446203]
5694530 = Cargo concentration in the ER [isa: 199977]
446219 = Synthesis of substrates in N-glycan biosythesis [isa: 446193]
4085001 = Sialic acid metabolism [isa: 446219]
446199 = Synthesis of Dolichyl-phosphate [isa: 446219]
446205 = Synthesis of GDP-mannose [isa: 446219]
446210 = Synthesis of UDP-N-acetyl-glucosamine [isa: 446219]
480985 = Synthesis of dolichyl-phosphate-glucose [isa: 446219]
6787639 = GDP-fucose biosynthesis [isa: 446219]
446353 = Cell-extracellular matrix interactions [isa: 446728]
446388 = Regulation of cytoskeletal remodeling and cell spreading by IPP complex components [isa: 446353]
9020702 = Interleukin-1 signaling [isa: 446652]
9007892 = Interleukin-38 signaling [isa: 446652]
9008059 = Interleukin-37 signaling [isa: 446652]
9014826 = Interleukin-36 pathway [isa: 446652]
9014843 = Interleukin-33 signaling [isa: 446652]
445989 = TAK1 activates NFkB by phosphorylation and activation of IKKs complex [isa: 166058 168164 9020702 937061 975138 975871]
373756 = SDK interactions [isa: 421270]
446107 = Type I hemidesmosome assembly [isa: 446728]
446343 = Localization of the PINCH-ILK-PARVIN complex to focal adhesions [isa: 446353]
447115 = Interleukin-12 family signaling [isa: 449147]
9020591 = Interleukin-12 signaling [isa: 447115]
9020933 = Interleukin-23 signaling [isa: 447115]
9020956 = Interleukin-27 signaling [isa: 447115]
937042 = IRAK2 mediated activation of TAK1 complex [isa: 166058 975871]
1266695 = Interleukin-7 signaling [isa: 449147]
448706 = Interleukin-1 processing [isa: 446652]
8984722 = Interleukin-35 Signalling [isa: 447115]
449836 = Other interleukin signaling [isa: 449147]
451927 = Interleukin-2 family signaling [isa: 449147]
512988 = Interleukin-3, Interleukin-5 and GM-CSF signaling [isa: 449147]
6783589 = Interleukin-6 family signaling [isa: 449147]
6783783 = Interleukin-10 signaling [isa: 449147]
6785807 = Interleukin-4 and Interleukin-13 signaling [isa: 449147]
8854691 = Interleukin-20 family signaling [isa: 449147]
202670 = ERKs are inactivated [isa: 198753]
450341 = Activation of the AP-1 family of transcription factors [isa: 450282]
450302 = activated TAK1 mediates p38 MAPK activation [isa: 450294]
450321 = JNK (c-Jun kinases) phosphorylation and  activation mediated by activated human TAK1 [isa: 450294]
5684264 = MAP3K8 (TPL2)-dependent MAPK1/3 activation [isa: 450294 9020702]
450531 = Regulation of mRNA stability by proteins that bind AU-rich elements [isa: 8953854]
450408 = AUF1 (hnRNP D0) binds and destabilizes mRNA [isa: 450531]
450513 = Tristetraprolin (TTP, ZFP36) binds and destabilizes mRNA [isa: 450531]
450520 = HuR (ELAVL1) binds and stabilizes mRNA [isa: 450531]
450604 = KSRP (KHSRP) binds and destabilizes mRNA [isa: 450531]
451306 = Ionotropic activity of kainate receptors [isa: 451326]
451308 = Activation of Ca-permeable Kainate Receptor [isa: 451306]
451307 = Activation of Na-permeable kainate receptors [isa: 451306]
500657 = Presynaptic function of Kainate receptors [isa: 451326]
8983432 = Interleukin-15 signaling [isa: 451927]
8985947 = Interleukin-9 signaling [isa: 451927]
9020558 = Interleukin-2 signaling [isa: 451927]
9020958 = Interleukin-21 signaling [isa: 451927]
912526 = Interleukin receptor SHC signaling [isa: 451927 512988]
380284 = Loss of proteins required for interphase microtubule organization from the centrosome [isa: 380287]
380259 = Loss of Nlp from mitotic centrosomes [isa: 380284]
141430 = Inactivation of APC/C via direct inhibition of the APC/C complex [isa: 141405]
69206 = G1/S Transition [isa: 453279]
113510 = E2F mediated regulation of DNA replication [isa: 69206]
69236 = G1 Phase [isa: 453279]
3214841 = PKMTs methylate histone lysines [isa: 3247509]
3296197 = Hydroxycarboxylic acid-binding receptors [isa: 373076]
419812 = Calcitonin-like ligand receptors [isa: 373080]
420499 = Class C/3 (Metabotropic glutamate/pheromone receptors) [isa: 500792]
912631 = Regulation of signaling by CBL [isa: 512988]
5173105 = O-linked glycosylation [isa: 597592]
913709 = O-linked glycosylation of mucins [isa: 5173105]
9663891 = Selective autophagy [isa: 1632852]
5205647 = Mitophagy [isa: 9663891]
5218859 = Regulated Necrosis [isa: 5357801]
5213460 = RIPK1-mediated regulated necrosis [isa: 5218859]
5250924 = B-WICH complex positively regulates rRNA expression [isa: 5250913]
427359 = SIRT1 negatively regulates rRNA expression [isa: 5250941]
427413 = NoRC negatively regulates rRNA expression [isa: 5250941]
901042 = Calnexin/calreticulin cycle [isa: 532668]
3371378 = Regulation by c-FLIP [isa: 140534]
418889 = Caspase activation via Dependence Receptors in the absence of ligand [isa: 5357769]
111452 = Activation and oligomerization of BAK protein [isa: 109606]
5675482 = Regulation of necroptotic cell death [isa: 5213460]
5358346 = Hedgehog ligand biogenesis [isa: 5358351]
5362798 = Release of Hh-Np from the secreting cell [isa: 5358346]
5610787 = Hedgehog 'off' state [isa: 5358351]
5632684 = Hedgehog 'on' state [isa: 5358351]
5358508 = Mismatch Repair [isa: 73894]
5358606 = Mismatch repair (MMR) directed by MSH2:MSH3 (MutSbeta) [isa: 5358508]
5362517 = Signaling by Retinoic Acid [isa: 9006931]
5368287 = Mitochondrial translation [isa: 72766]
5419276 = Mitochondrial translation termination [isa: 5368287]
549127 = Organic cation transport [isa: 549132]
561048 = Organic anion transport [isa: 549132]
1482788 = Acyl chain remodelling of PC [isa: 1483206]
400206 = Regulation of lipid metabolism by PPARalpha [isa: 556833]
1660661 = Sphingolipid de novo biosynthesis [isa: 428157]
74182 = Ketone body metabolism [isa: 556833]
8848584 = Wax and plasmalogen biosynthesis [isa: 556833]
6807062 = Cholesterol biosynthesis via lathosterol [isa: 191273]
8964572 = Lipid particle organization [isa: 556833]
2046106 = alpha-linolenic acid (ALA) metabolism [isa: 2046104]
8979227 = Triglyceride metabolism [isa: 556833]
9018678 = Biosynthesis of specialized proresolving mediators (SPMs) [isa: 556833]
5576886 = Phase 4 - resting membrane potential [isa: 5576891]
5576890 = Phase 3 - rapid repolarisation [isa: 5576891]
5576892 = Phase 0 - rapid depolarisation [isa: 5576891]
5576893 = Phase 2 - plateau phase [isa: 5576891]
5576894 = Phase 1 - inactivation of fast Na+ channels [isa: 5576891]
5578768 = Physiological factors [isa: 5576891]
5578775 = Ion homeostasis [isa: 5576891]
5607764 = CLEC7A (Dectin-1) signaling [isa: 5621481]
5607763 = CLEC7A (Dectin-1) induces NFAT activation [isa: 5607764]
5660668 = CLEC7A/inflammasome pathway [isa: 5607764]
5610780 = Degradation of GLI1 by the proteasome [isa: 5610787]
5610785 = GLI3 is processed to GLI3R by the proteasome [isa: 5610787]
5620912 = Anchoring of the basal body to the plasma membrane [isa: 5617833]
5620920 = Cargo trafficking to the periciliary membrane [isa: 5617833]
5620924 = Intraflagellar transport [isa: 5617833]
5620916 = VxPx cargo-targeting to cilium [isa: 5620920]
5620922 = BBSome-mediated cargo-targeting to cilium [isa: 5620920]
5624138 = Trafficking of myristoylated proteins to the cilium [isa: 5620920]
5624958 = ARL13B-mediated ciliary trafficking of INPP5E [isa: 5620920]
5607761 = Dectin-1 mediated noncanonical NF-kB signaling [isa: 5607764]
5621480 = Dectin-2 family [isa: 5621481]
5621575 = CD209 (DC-SIGN) signaling [isa: 5621481]
5625886 = Activated PKN1 stimulates transcription of AR (androgen receptor) regulated genes KLK2 and KLK3 [isa: 5625740]
5632681 = Ligand-receptor interactions [isa: 5632684]
5635838 = Activation of SMO [isa: 5632684]
6804756 = Regulation of TP53 Activity through Phosphorylation [isa: 5633007]
6804758 = Regulation of TP53 Activity through Acetylation [isa: 5633007]
6804759 = Regulation of TP53 Activity through Association with Co-factors [isa: 5633007]
6804760 = Regulation of TP53 Activity through Methylation [isa: 5633007]
6806003 = Regulation of TP53 Expression and Degradation [isa: 5633007]
6803204 = TP53 Regulates Transcription of Genes Involved in Cytochrome C Release [isa: 5633008]
6803205 = TP53 regulates transcription of several additional cell death genes whose specific roles in p53-dependent apoptosis remain uncertain [isa: 5633008]
6803207 = TP53 Regulates Transcription of Caspase Activators and Caspases [isa: 5633008]
6803211 = TP53 Regulates Transcription of Death Receptors and Ligands [isa: 5633008]
5652084 = Fructose metabolism [isa: 71387]
70350 = Fructose catabolism [isa: 5652084]
190704 = Oligomerization of connexins into connexons [isa: 190861]
2168880 = Scavenging of heme from plasma [isa: 2173782]
5654687 = Downstream signaling of activated FGFR1 [isa: 5654736]
5654688 = SHC-mediated cascade:FGFR1 [isa: 5654687]
5654689 = PI-3K cascade:FGFR1 [isa: 5654687]
5654693 = FRS-mediated FGFR1 signaling [isa: 5654687]
5654696 = Downstream signaling of activated FGFR2 [isa: 5654738]
5654695 = PI-3K cascade:FGFR2 [isa: 5654696]
5654699 = SHC-mediated cascade:FGFR2 [isa: 5654696]
5654700 = FRS-mediated FGFR2 signaling [isa: 5654696]
5654708 = Downstream signaling of activated FGFR3 [isa: 5654741]
5654704 = SHC-mediated cascade:FGFR3 [isa: 5654708]
5654706 = FRS-mediated FGFR3 signaling [isa: 5654708]
5654710 = PI-3K cascade:FGFR3 [isa: 5654708]
5654716 = Downstream signaling of activated FGFR4 [isa: 5654743]
5654712 = FRS-mediated FGFR4 signaling [isa: 5654716]
5654719 = SHC-mediated cascade:FGFR4 [isa: 5654716]
5654720 = PI-3K cascade:FGFR4 [isa: 5654716]
5654726 = Negative regulation of FGFR1 signaling [isa: 5654736]
5654727 = Negative regulation of FGFR2 signaling [isa: 5654738]
5654732 = Negative regulation of FGFR3 signaling [isa: 5654741]
5654733 = Negative regulation of FGFR4 signaling [isa: 5654743]
190370 = FGFR1b ligand binding and activation [isa: 190242]
5654219 = Phospholipase C-mediated cascade: FGFR1 [isa: 5654687]
1295596 = Spry regulation of FGF signaling [isa: 5654726 5654727 5654732 5654733]
5658623 = FGFRL1 modulation of FGFR1 signaling [isa: 5654736]
190375 = FGFR2c ligand binding and activation [isa: 190241]
5654221 = Phospholipase C-mediated cascade; FGFR2 [isa: 5654696]
6803529 = FGFR2 alternative splicing [isa: 5654738]
190371 = FGFR3b ligand binding and activation [isa: 190239]
5654227 = Phospholipase C-mediated cascade; FGFR3 [isa: 5654708]
1307965 = betaKlotho-mediated ligand binding [isa: 190322]
5654228 = Phospholipase C-mediated cascade; FGFR4 [isa: 5654716]
5660526 = Response to metal ions [isa: 8953897]
5669034 = TNFs bind their physiological receptors [isa: 5668541]
5676590 = NIK-->noncanonical NF-kB signaling [isa: 5668541]
5676594 = TNF receptor superfamily (TNFSF) members mediating non-canonical NF-kB pathway [isa: 5668541]
5673001 = RAF/MAP kinase cascade [isa: 5684996 9607240]
5673000 = RAF activation [isa: 5673001]
5674135 = MAP2K and MAPK activation [isa: 5673001]
5675221 = Negative regulation of MAPK pathway [isa: 5673001]
5674499 = Negative feedback regulation of MAPK pathway [isa: 5675221]
5218900 = CASP8 activity is inhibited [isa: 5675482]
112411 = MAPK1 (ERK2) activation [isa: 112409]
5687128 = MAPK6/MAPK4 signaling [isa: 5683057]
110056 = MAPK3 (ERK1) activation [isa: 112409]
5658442 = Regulation of RAS by GAPs [isa: 5673001]
5693532 = DNA Double-Strand Break Repair [isa: 73894]
5693538 = Homology Directed Repair [isa: 5693532]
5688426 = Deubiquitination [isa: 597592]
5689877 = Josephin domain DUBs [isa: 5688426]
5689880 = Ub-specific processing proteases [isa: 5688426]
5689896 = Ovarian tumor domain proteases [isa: 5688426]
5689901 = Metalloprotease DUBs [isa: 5688426]
5693567 = HDR through Homologous Recombination (HRR) or Single Strand Annealing (SSA) [isa: 5693538]
5693571 = Nonhomologous End-Joining (NHEJ) [isa: 5693532]
5693606 = DNA Double Strand Break Response [isa: 5693532]
5685942 = HDR through Homologous Recombination (HRR) [isa: 5693567]
5693537 = Resolution of D-Loop Structures [isa: 5685942]
5693568 = Resolution of D-loop Structures through Holliday Junction Intermediates [isa: 5693537]
5693579 = Homologous DNA Pairing and Strand Exchange [isa: 5685942]
5685938 = HDR through Single Strand Annealing (SSA) [isa: 5693567]
5693554 = Resolution of D-loop Structures through Synthesis-Dependent Strand Annealing (SDSA) [isa: 5693537]
5693607 = Processing of DNA double-strand break ends [isa: 5693567]
5693616 = Presynaptic phase of homologous DNA pairing and strand exchange [isa: 5693579]
5693548 = Sensing of DNA Double Strand Breaks [isa: 5693606]
5693565 = Recruitment and ATM-mediated phosphorylation of repair and signaling proteins at DNA double strand breaks [isa: 5693606]
5696398 = Nucleotide Excision Repair [isa: 73894]
6781827 = Transcription-Coupled Nucleotide Excision Repair (TC-NER) [isa: 5696398]
5696399 = Global Genome Nucleotide Excision Repair (GG-NER) [isa: 5696398]
5696395 = Formation of Incision Complex in GG-NER [isa: 5696399]
5696397 = Gap-filling DNA repair synthesis and ligation in GG-NER [isa: 5696399]
5696400 = Dual Incision in GG-NER [isa: 5696399]
162699 = Synthesis of dolichyl-phosphate mannose [isa: 163125 446219]
159740 = Gamma-carboxylation of protein precursors [isa: 159854]
3108214 = SUMOylation of DNA damage response and repair proteins [isa: 3108232]
5173214 = O-glycosylation of TSR domain-containing proteins [isa: 5173105]
5689603 = UCH proteinases [isa: 5688426]
8852135 = Protein ubiquitination [isa: 597592]
8873719 = RAB geranylgeranylation [isa: 597592]
8876725 = Protein methylation [isa: 597592]
8951664 = Neddylation [isa: 597592]
8955332 = Carboxyterminal post-translational modifications of tubulin [isa: 597592]
8957275 = Post-translational protein phosphorylation [isa: 597592]
6799198 = Complex I biogenesis [isa: 611105]
844455 = The NLRP1 inflammasome [isa: 622312]
844456 = The NLRP3 inflammasome [isa: 622312]
844615 = The AIM2 inflammasome [isa: 622312]
629587 = Highly sodium permeable postsynaptic acetylcholine nicotinic receptors [isa: 622323 622327]
629597 = Highly calcium permeable nicotinic acetylcholine receptors [isa: 622323]
629594 = Highly calcium permeable postsynaptic nicotinic acetylcholine receptors [isa: 622327]
6781823 = Formation of TC-NER Pre-Incision Complex [isa: 6781827]
6782135 = Dual incision in TC-NER [isa: 6781827]
6782210 = Gap-filling DNA repair synthesis and ligation in TC-NER [isa: 6781827]
1059683 = Interleukin-6 signaling [isa: 6783589]
6788467 = IL-6-type cytokine receptor ligand interactions [isa: 6783589]
6804114 = TP53 Regulates Transcription of Genes Involved in G2 Cell Cycle Arrest [isa: 6791312]
6804115 = TP53 regulates transcription of additional cell cycle genes whose exact role in the p53 pathway remain uncertain [isa: 6791312]
6804116 = TP53 Regulates Transcription of Genes Involved in G1 Cell Cycle Arrest [isa: 6791312]
388844 = Receptor-type tyrosine-protein phosphatases [isa: 6794362]
6794361 = Neurexins and neuroligins [isa: 6794362]
8849932 = Synaptic adhesion-like molecules [isa: 6794362]
1461957 = Beta defensins [isa: 1461973]
6799990 = Metal sequestration by antimicrobial proteins [isa: 6803157]
6803544 = Ion influx/efflux at host-pathogen interface [isa: 6803157]
6811555 = PI5P Regulates TP53 Acetylation [isa: 6804758]
6809371 = Formation of the cornified envelope [isa: 6805567]
6804754 = Regulation of TP53 Expression [isa: 6806003]
6804757 = Regulation of TP53 Degradation [isa: 6806003]
6806664 = Metabolism of vitamin K [isa: 6806667]
8877627 = Vitamin E [isa: 6806667]
6806834 = Signaling by MET [isa: 9006934]
6807004 = Negative regulation of MET activity [isa: 6806834]
8851805 = MET activates RAS signaling [isa: 6806834]
8851907 = MET activates PI3K/AKT signaling [isa: 6806834]
8865999 = MET activates PTPN11 [isa: 6806834]
8875656 = MET receptor recycling [isa: 6806834]
8875791 = MET activates STAT3 [isa: 6806834]
8875878 = MET promotes cell motility [isa: 6806834]
8943724 = Regulation of PTEN gene transcription [isa: 6807070]
8948747 = Regulation of PTEN localization [isa: 6807070]
8948751 = Regulation of PTEN stability and activity [isa: 6807070]
6811438 = Intra-Golgi traffic [isa: 6811442]
6811440 = Retrograde transport at the Trans-Golgi-Network [isa: 6811442]
8856688 = Golgi-to-ER retrograde transport [isa: 6811442]
69002 = DNA Replication Pre-Initiation [isa: 69306]
68867 = Assembly of the pre-replicative complex [isa: 69002]
68827 = CDT1 association with the CDC6:ORC:origin complex [isa: 68867]
3301854 = Nuclear Pore Complex (NPC) Disassembly [isa: 2980766]
2299718 = Condensation of Prophase Chromosomes [isa: 68875]
2465910 = MASTL Facilitates Mitotic Progression [isa: 68875]
2980767 = Activation of NIMA Kinases NEK9, NEK6, NEK7 [isa: 2980766]
68877 = Mitotic Prometaphase [isa: 68886]
2514853 = Condensation of Prometaphase Chromosomes [isa: 68877]
380320 = Recruitment of NuMA to mitotic centrosomes [isa: 68877]
9648025 = EML4 and NUDC in mitotic spindle formation [isa: 68877]
2467813 = Separation of Sister Chromatids [isa: 68882]
2995410 = Nuclear Envelope (NE) Reassembly [isa: 68882]
68884 = Mitotic Telophase/Cytokinesis [isa: 68886]
2995383 = Initiation of Nuclear Envelope (NE) Reformation [isa: 2995410]
162658 = Golgi Cisternae Pericentriolar Stack Reorganization [isa: 68875]
2500257 = Resolution of Sister Chromatid Cohesion [isa: 68877]
2470946 = Cohesin Loading onto Chromatin [isa: 68884]
68689 = CDC6 association with the ORC:origin complex [isa: 68867]
68962 = Activation of the pre-replicative complex [isa: 69002 69206]
69242 = S Phase [isa: 69278]
69239 = Synthesis of DNA [isa: 69242 69306]
69190 = DNA strand elongation [isa: 69239]
69186 = Lagging Strand Synthesis [isa: 69190]
69183 = Processive synthesis on the lagging strand [isa: 69186]
69166 = Removal of the Flap Intermediate [isa: 69183]
69109 = Leading Strand Synthesis [isa: 69190]
69091 = Polymerase switching [isa: 69109 69186]
69202 = Cyclin E associated events during G1/S transition  [isa: 69206]
69200 = Phosphorylation of proteins involved in G1/S transition by active Cyclin E:Cdk2 complexes [isa: 69202]
113501 = Inhibition of replication initiation of damaged DNA by RB1/E2F1 [isa: 113510]
69656 = Cyclin A:Cdk2-associated events at S phase entry [isa: 69242]
69231 = Cyclin D associated events in G1 [isa: 69236]
69052 = Switching of origins to a post-replicative state [isa: 69239]
68949 = Orc1 removal from chromatin [isa: 69052]
176974 = Unwinding of DNA [isa: 69190]
69017 = CDK-mediated phosphorylation and removal of Cdc6 [isa: 69052]
68952 = DNA replication initiation [isa: 69239]
187577 = SCF(Skp2)-mediated degradation of p27/p21 [isa: 69202 69656]
75815 = Ubiquitin-dependent degradation of Cyclin D [isa: 69242]
156711 = Polo-like kinase mediated events [isa: 69275]
2565942 = Regulation of PLK1 Activity at G2/M Transition [isa: 69275]
380270 = Recruitment of mitotic centrosome proteins and complexes [isa: 380287]
69273 = Cyclin A/B1/B2 associated events during G2/M transition [isa: 69275]
8852276 = The role of GTSE1 in G2/M progression after G2 checkpoint [isa: 69275]
8854050 = FBXL7 down-regulates AURKA during mitotic entry and in early mitosis [isa: 69275]
8854518 = AURKA Activation by TPX2 [isa: 69275]
8854521 = Interaction between PHLDA1 and AURKA [isa: 69275]
68911 = G2 Phase [isa: 453274]
174084 = Autodegradation of Cdh1 by Cdh1:APC/C [isa: 174143]
1538133 = G0 and Early G1 [isa: 453279]
68881 = Mitotic Metaphase/Anaphase Transition [isa: 2555396]
2468052 = Establishment of Sister Chromatid Cohesion [isa: 69242]
68616 = Assembly of the ORC complex at the origin of replication [isa: 68867]
69481 = G2/M Checkpoints [isa: 69620]
69473 = G2/M DNA damage checkpoint [isa: 69481]
75035 = Chk1/Chk2(Cds1) mediated inactivation of Cyclin B:Cdk1 complex [isa: 69473]
69478 = G2/M DNA replication checkpoint [isa: 69481]
69615 = G1/S DNA Damage Checkpoints [isa: 69620]
69580 = p53-Dependent G1/S DNA damage checkpoint [isa: 69615]
69563 = p53-Dependent G1 DNA Damage Response [isa: 69580]
69560 = Transcriptional activation of p53 responsive genes   [isa: 69563]
69895 = Transcriptional  activation of  cell cycle inhibitor p21  [isa: 69560]
69613 = p53-Independent G1/S DNA damage checkpoint [isa: 69615]
69610 = p53-Independent DNA Damage Response [isa: 69613]
69541 = Stabilization of p53 [isa: 69563]
69601 = Ubiquitin Mediated Degradation of Phosphorylated Cdc25A [isa: 69610]
141424 = Amplification of signal from the kinetochores [isa: 69618]
141444 = Amplification  of signal from unattached  kinetochores via a MAD2  inhibitory signal [isa: 141424]
176187 = Activation of ATR in response to replication stress [isa: 69481]
349425 = Autodegradation of the E3 ubiquitin ligase COP1 [isa: 69541]
70326 = Glucose metabolism [isa: 71387]
70171 = Glycolysis [isa: 70326]
70268 = Pyruvate metabolism [isa: 71406]
9634600 = Regulation of glycolysis by fructose 2,6-bisphosphate metabolism [isa: 70171]
70263 = Gluconeogenesis [isa: 70326]
1237112 = Methionine salvage pathway [isa: 1614635]
209905 = Catecholamine biosynthesis [isa: 209776]
2408508 = Metabolism of ingested SeMet, Sec, MeSec into H2Se [isa: 2408522]
350562 = Regulation of ornithine decarboxylase (ODC) [isa: 351202]
6783984 = Glycine degradation [isa: 389661]
5662702 = Melanin biosynthesis [isa: 71291]
6798163 = Choline catabolism [isa: 71291]
70635 = Urea cycle [isa: 71291]
70688 = Proline catabolism [isa: 71291]
70895 = Branched-chain amino acid catabolism [isa: 71291]
70921 = Histidine catabolism [isa: 71291]
71064 = Lysine catabolism [isa: 71291]
71240 = Tryptophan catabolism [isa: 71291]
71262 = Carnitine synthesis [isa: 71291]
71288 = Creatine metabolism [isa: 71291]
8849175 = Threonine catabolism [isa: 71291]
8963691 = Phenylalanine and tyrosine metabolism [isa: 71291]
8963693 = Aspartate and asparagine metabolism [isa: 71291]
8964539 = Glutamate and glutamine metabolism [isa: 71291]
8964540 = Alanine metabolism [isa: 71291]
977347 = Serine biosynthesis [isa: 71291]
71336 = Pentose phosphate pathway [isa: 71387]
2022854 = Keratan sulfate biosynthesis [isa: 1638074]
5652227 = Fructose biosynthesis [isa: 5652084]
5653890 = Lactose synthesis [isa: 71387]
5661270 = Formation of xylulose-5-phosphate [isa: 71387]
170822 = Regulation of Glucokinase by Glucokinase Regulatory Protein [isa: 70171]
70370 = Galactose catabolism [isa: 71387]
73843 = 5-Phosphoribose 1-diphosphate biosynthesis [isa: 71336]
8853383 = Lysosomal oligosaccharide catabolism [isa: 71387]
8982491 = Glycogen metabolism [isa: 71387]
9033658 = Blood group systems biosynthesis [isa: 71387]
204174 = Regulation of pyruvate dehydrogenase (PDH) complex [isa: 70268]
71403 = Citric acid cycle (TCA cycle) [isa: 71406]
880009 = Interconversion of 2-oxoglutarate and 2-hydroxyglutarate [isa: 71406]
159231 = Transport of Mature mRNA Derived from an Intronless Transcript [isa: 159234]
72172 = mRNA Splicing [isa: 72203]
75072 = mRNA Editing [isa: 8953854]
159230 = Transport of the SLBP Dependant Mature mRNA [isa: 159234]
159236 = Transport of Mature mRNA derived from an Intron-Containing Transcript [isa: 72202]
72165 = mRNA Splicing - Minor Pathway [isa: 72172]
72187 = mRNA 3'-end processing [isa: 72203]
159227 = Transport of the SLBP independent Mature mRNA [isa: 159234]
72312 = rRNA processing [isa: 8953854]
72613 = Eukaryotic Translation Initiation [isa: 72766]
72737 = Cap-dependent Translation Initiation [isa: 72613]
72662 = Activation of the mRNA upon binding of the cap-binding complex and eIFs, and subsequent binding to 43S [isa: 72737]
72649 = Translation initiation complex formation [isa: 72662]
72689 = Formation of a pool of free 40S subunits [isa: 72737]
72695 = Formation of the ternary complex, and subsequently, the 43S complex [isa: 72737]
72702 = Ribosomal scanning and start codon recognition [isa: 72737]
72706 = GTP hydrolysis and joining of the 60S ribosomal subunit [isa: 72737]
72731 = Recycling of eIF2:GDP [isa: 72737]
156902 = Peptide chain elongation [isa: 156842]
1799339 = SRP-dependent cotranslational protein targeting to membrane [isa: 72766]
379716 = Cytosolic tRNA aminoacylation [isa: 379724]
5389840 = Mitochondrial translation elongation [isa: 5368287]
156827 = L13a-mediated translational silencing of Ceruloplasmin expression [isa: 72613]
72764 = Eukaryotic Translation Termination [isa: 72766]
73864 = RNA Polymerase I Transcription [isa: 74160]
73854 = RNA Polymerase I Promoter Clearance [isa: 73864]
73772 = RNA Polymerase I Promoter Escape [isa: 73854]
674695 = RNA Polymerase II Pre-transcription Events [isa: 73857]
6807505 = RNA polymerase II transcribes snRNA genes [isa: 73857]
73779 = RNA Polymerase II Transcription Pre-Initiation And Promoter Opening [isa: 73857]
73856 = RNA Polymerase II Transcription Termination [isa: 73857]
75955 = RNA Polymerase II Transcription Elongation [isa: 73857]
76042 = RNA Polymerase II Transcription Initiation And Promoter Clearance [isa: 73857]
77075 = RNA Pol II CTD phosphorylation and interaction with CE [isa: 73857]
73762 = RNA Polymerase I Transcription Initiation [isa: 73854]
73863 = RNA Polymerase I Transcription Termination [isa: 73864]
110373 = Resolution of AP sites via the multiple-nucleotide patch replacement pathway [isa: 73933]
5651801 = PCNA-Dependent Long Patch Base Excision Repair [isa: 110373]
171306 = Packaging Of Telomere Ends [isa: 157579]
774815 = Nucleosome assembly [isa: 73886]
209543 = p75NTR recruits signalling complexes [isa: 193639]
75157 = FasL/ CD95L signaling [isa: 73887]
75158 = TRAIL  signaling [isa: 73887]
75893 = TNF signaling [isa: 73887]
110320 = Translesion Synthesis by POLH [isa: 110313]
110314 = Recognition of DNA damage by PCNA-containing replication complex [isa: 73893]
5358565 = Mismatch repair (MMR) directed by MSH2:MSH6 (MutSalpha) [isa: 5358508]
5685939 = HDR through MMEJ (alt-NHEJ) [isa: 5693538]
5696394 = DNA Damage Recognition in GG-NER [isa: 5696399]
6783310 = Fanconi Anemia Pathway [isa: 73894]
73929 = Base-Excision Repair, AP Site Formation [isa: 73884]
110312 = Translesion synthesis by REV1 [isa: 110313]
73942 = DNA Damage Reversal [isa: 73894]
73927 = Depurination [isa: 73929]
110331 = Cleavage of the damaged purine [isa: 73927]
73928 = Depyrimidination [isa: 73929]
110329 = Cleavage of the damaged pyrimidine  [isa: 73928]
110330 = Recognition and association of DNA glycosylase with site containing an affected purine [isa: 73927]
110328 = Recognition and association of DNA glycosylase with site containing an affected pyrimidine [isa: 73928]
110357 = Displacement of DNA glycosylase by APEX1 [isa: 73933]
110362 = POLB-Dependent Long Patch Base Excision Repair [isa: 110373]
73930 = Abasic sugar-phosphate removal via the single-nucleotide replacement pathway [isa: 110381]
5649702 = APEX1-Independent Resolution of AP Sites via the Single Nucleotide Replacement Pathway [isa: 73933]
73943 = Reversal of alkylation damage by DNA dioxygenases [isa: 73942]
74158 = RNA Polymerase III Transcription [isa: 74160]
203927 = MicroRNA (miRNA) biogenesis [isa: 211000]
212300 = PRC2 methylates histones and DNA [isa: 212165]
2032785 = YAP1- and WWTR1 (TAZ)-stimulated gene expression [isa: 212436]
73728 = RNA Polymerase I Promoter Opening [isa: 73854]
76046 = RNA Polymerase III Transcription Initiation [isa: 74158]
75944 = Transcription from mitochondrial promoters [isa: 74160]
77108 = Utilization of Ketone Bodies [isa: 74182]
77111 = Synthesis of Ketone Bodies [isa: 74182]
74259 = Purine catabolism [isa: 8956319]
165158 = Activation of AKT2 [isa: 109704]
112412 = SOS-mediated signalling [isa: 112399]
74749 = Signal attenuation [isa: 74751]
74713 = IRS activation [isa: 74751]
77387 = Insulin receptor recycling [isa: 74752]
75064 = mRNA Editing: A to I Conversion [isa: 75072]
77042 = Formation of editosomes by ADAR proteins [isa: 75064]
75067 = Processing of Capped Intronless Pre-mRNA [isa: 8953854]
77588 = SLBP Dependent Processing of Replication-Dependent Histone Pre-mRNAs [isa: 75067]
77595 = Processing of Intronless Pre-mRNAs [isa: 75067]
72200 = mRNA Editing: C to U Conversion [isa: 75072]
75102 = C6 deamination of adenosine [isa: 75064]
75105 = Fatty acyl-CoA biosynthesis [isa: 8978868]
264870 = Caspase-mediated cleavage of cytoskeletal proteins [isa: 111465]
140342 = Apoptosis induced DNA fragmentation [isa: 75153]
5357786 = TNFR1-induced proapoptotic signaling [isa: 75893]
5357905 = Regulation of TNFR1 signaling [isa: 75893]
5357956 = TNFR1-induced NFkappaB signaling pathway [isa: 75893]
5626978 = TNFR1-mediated ceramide production [isa: 75893]
163282 = Mitochondrial transcription initiation [isa: 75944]
163316 = Mitochondrial transcription termination [isa: 75944]
112382 = Formation of RNA Pol II elongation complex  [isa: 75955]
113418 = Formation of the Early Elongation Complex [isa: 75955]
114604 = GPVI-mediated activation cascade [isa: 76002]
392170 = ADP signalling through P2Y purinoceptor 12 [isa: 392518]
430116 = GP1b-IX-V activation signalling [isa: 76002]
456926 = Thrombin signalling through proteinase activated receptors (PARs) [isa: 76002]
76005 = Response to elevated platelet cytosolic Ca2+ [isa: 76002]
354194 = GRB2:SOS provides linkage to MAPK signaling for Integrins  [isa: 354192]
114516 = Disinhibition of SNARE formation [isa: 76005]
114608 = Platelet degranulation  [isa: 76005]
392023 = Adrenaline signalling through Alpha-2 adrenergic receptor [isa: 76009]
73776 = RNA Polymerase II Promoter Escape [isa: 76042]
75953 = RNA Polymerase II Transcription Initiation [isa: 76042]
76061 = RNA Polymerase III Transcription Initiation From Type 1 Promoter [isa: 76046]
76066 = RNA Polymerase III Transcription Initiation From Type 2 Promoter [isa: 76046]
76071 = RNA Polymerase III Transcription Initiation From Type 3 Promoter [isa: 76046]
77289 = Mitochondrial Fatty Acid Beta-Oxidation [isa: 8978868]
77286 = mitochondrial fatty acid beta-oxidation of saturated fatty acids [isa: 77289]
77310 = Beta oxidation of lauroyl-CoA to decanoyl-CoA-CoA [isa: 77286]
77346 = Beta oxidation of decanoyl-CoA to octanoyl-CoA-CoA [isa: 77286]
77348 = Beta oxidation of octanoyl-CoA to hexanoyl-CoA [isa: 77286]
77350 = Beta oxidation of hexanoyl-CoA to butanoyl-CoA [isa: 77286]
77352 = Beta oxidation of butanoyl-CoA to acetyl-CoA [isa: 77286]
77305 = Beta oxidation of palmitoyl-CoA to myristoyl-CoA [isa: 77286]
77285 = Beta oxidation of myristoyl-CoA to lauroyl-CoA [isa: 77286]
77288 = mitochondrial fatty acid beta-oxidation of unsaturated fatty acids [isa: 77289]
606279 = Deposition of new CENPA-containing nucleosomes at the centromere [isa: 774815]
877300 = Interferon gamma signaling [isa: 913531]
2179392 = EGFR Transactivation by Gastrin [isa: 881907]
8847993 = ERBB2 Activates PTK6 Signaling [isa: 1227986 8848021]
8849468 = PTK6 Regulates Proteins Involved in RNA Processing [isa: 8848021]
8849469 = PTK6 Regulates RTKs and Their Effectors AKT1 and DOK1 [isa: 8848021]
8849470 = PTK6 Regulates Cell Cycle [isa: 8848021]
8849471 = PTK6 Regulates RHO GTPases, RAS GTPase and MAP kinases [isa: 8848021]
8849472 = PTK6 Down-Regulation [isa: 8848021]
8849474 = PTK6 Activates STAT3 [isa: 8848021]
8857538 = PTK6 promotes HIF1A stabilization [isa: 8848021]
75896 = Plasmalogen biosynthesis [isa: 8848584]
9640463 = Wax biosynthesis [isa: 8848584]
8866652 = Synthesis of active ubiquitin: roles of E1 and E2 enzymes [isa: 8852135]
8866654 = E3 ubiquitin ligases ubiquitinate target proteins [isa: 8852135]
6811434 = COPI-dependent Golgi-to-ER retrograde traffic [isa: 8856688]
6811436 = COPI-independent Golgi-to-ER retrograde traffic [isa: 8856688]
8856825 = Cargo recognition for clathrin-mediated endocytosis [isa: 8856828]
1358803 = Downregulation of ERBB2:ERBB3 signaling [isa: 8863795]
8866904 = Negative regulation of activity of TFAP2 (AP-2) family transcription factors [isa: 8864260]
8866907 = Activation of the TFAP2 (AP-2) family of transcription factors [isa: 8864260]
8866910 = TFAP2 (AP-2) family regulates transcription of growth factors and their receptors [isa: 8864260]
8866911 = TFAP2 (AP-2) family regulates transcription of cell cycle factors [isa: 8864260]
8869496 = TFAP2A acts as a transcriptional repressor during retinoic acid induced cell differentiation [isa: 8864260]
8868773 = rRNA processing in the nucleus and cytosol [isa: 72312]
8874081 = MET activates PTK2 signaling [isa: 8875878]
8875513 = MET interacts with TNS proteins [isa: 8875878]
8875555 = MET activates RAP1 and RAC1 [isa: 8875878]
8941855 = RUNX3 regulates CDKN1A transcription [isa: 8878159]
8941856 = RUNX3 regulates NOTCH signaling [isa: 8878159]
8941858 = Regulation of RUNX3 expression and activity [isa: 8878159]
8951430 = RUNX3 regulates WNT signaling [isa: 8878159]
8951671 = RUNX3 regulates YAP1-mediated transcription [isa: 8878159]
8951936 = RUNX3 regulates p14-ARF [isa: 8878159]
8939902 = Regulation of RUNX2 expression and activity [isa: 8878166]
8941326 = RUNX2 regulates bone development [isa: 8878166]
8877330 = RUNX1 and FOXP3 control the development of regulatory T lymphocytes (Tregs) [isa: 8878171]
8931987 = RUNX1 regulates estrogen receptor mediated transcription [isa: 8878171]
8934593 = Regulation of RUNX1 Expression and Activity [isa: 8878171]
8936459 = RUNX1 regulates genes involved in megakaryocyte differentiation and platelet function [isa: 8878171]
8939236 = RUNX1 regulates transcription of genes involved in differentiation of HSCs [isa: 8878171]
8939242 = RUNX1 regulates transcription of genes involved in differentiation of keratinocytes [isa: 8878171]
8939243 = RUNX1 interacts with co-factors whose precise effect on RUNX1 targets is not known [isa: 8878171]
8939245 = RUNX1 regulates transcription of genes involved in BCR signaling [isa: 8878171]
8939246 = RUNX1 regulates transcription of genes involved in differentiation of myeloid cells [isa: 8878171]
8939247 = RUNX1 regulates transcription of genes involved in interleukin signaling [isa: 8878171]
888568 = GABA synthesis [isa: 888590]
888593 = Reuptake of GABA [isa: 888590]
916853 = Degradation of GABA [isa: 888590]
8935690 = Digestion [isa: 8963743]
192456 = Digestion of dietary lipid [isa: 8935690]
8939211 = ESR-mediated signaling [isa: 9006931]
9018519 = Estrogen-dependent gene expression [isa: 8939211]
8940973 = RUNX2 regulates osteoblast differentiation [isa: 8941326]
8949664 = Processing of SMDT1 [isa: 8949215]
191859 = snRNP Assembly [isa: 194441]
429947 = Deadenylation of mRNA [isa: 429914]
450385 = Butyrate Response Factor 1 (BRF1) binds and destabilizes mRNA [isa: 450531]
72086 = mRNA Capping [isa: 8953854]
72163 = mRNA Splicing - Major Pathway [isa: 72172]
6791226 = Major pathway of rRNA processing in the nucleolus and cytosol [isa: 8868773]
111367 = SLBP independent Processing of Histone Pre-mRNAs [isa: 75067]
75094 = Formation of the Editosome [isa: 72200]
927802 = Nonsense-Mediated Decay (NMD) [isa: 8953854]
1234158 = Regulation of gene expression by Hypoxia-inducible Factor [isa: 1234174]
5661231 = Metallothioneins bind metals [isa: 5660526]
73621 = Pyrimidine catabolism [isa: 8956319]
2393930 = Phosphate bond hydrolysis by NUDT proteins [isa: 74259]
8850843 = Phosphate bond hydrolysis by NTPDase proteins [isa: 8956319]
500753 = Pyrimidine biosynthesis [isa: 8956320]
73817 = Purine ribonucleoside monophosphate biosynthesis [isa: 8956320]
73614 = Pyrimidine salvage [isa: 8956321]
74217 = Purine salvage [isa: 8956321]
1655829 = Regulation of cholesterol biosynthesis by SREBP (SREBF) [isa: 8957322]
6807047 = Cholesterol biosynthesis via desmosterol [isa: 191273]
159418 = Recycling of bile acids and salts [isa: 194068]
193048 = Androgen biosynthesis [isa: 196071]
196791 = Vitamin D (calciferol) metabolism [isa: 8957322]
8963676 = Intestinal absorption [isa: 8963743]
8981373 = Intestinal hexose absorption [isa: 8963676]
8963684 = Tyrosine catabolism [isa: 8963691]
8964208 = Phenylalanine metabolism [isa: 8963691]
189085 = Digestion of dietary carbohydrate [isa: 8935690]
8963678 = Intestinal lipid absorption [isa: 8963676]
8866423 = VLDL assembly [isa: 8963898]
8963888 = Chylomicron assembly [isa: 8963898]
8963896 = HDL assembly [isa: 8963898]
8963889 = Assembly of active LPL and LIPC lipase complexes [isa: 8963899]
8963901 = Chylomicron remodeling [isa: 8963899]
8964041 = LDL remodeling [isa: 8963899]
8964058 = HDL remodeling [isa: 8963899]
8866427 = VLDLR internalisation and degradation [isa: 8964043]
8964011 = HDL clearance [isa: 8964043]
8964026 = Chylomicron clearance [isa: 8964043]
8964038 = LDL clearance [isa: 8964043]
8964046 = VLDL clearance [isa: 8964043]
200425 = Carnitine metabolism [isa: 8978868]
2046105 = Linoleic acid (LA) metabolism [isa: 2046104]
2142670 = Synthesis of epoxy (EET) and dihydroxyeicosatrienoic acids (DHET) [isa: 2142753]
389599 = Alpha-oxidation of phytanate [isa: 390918]
75876 = Synthesis of very long-chain fatty acyl-CoAs [isa: 75105]
71032 = Propionyl-CoA catabolism [isa: 77289]
1474151 = Tetrahydrobiopterin (BH4) synthesis, recycling, salvage and regulation [isa: 8978934]
2142789 = Ubiquinol biosynthesis [isa: 8978934]
389542 = NADPH regeneration [isa: 8978934]
163560 = Triglyceride catabolism [isa: 8979227]
75109 = Triglyceride biosynthesis [isa: 8979227]
3322077 = Glycogen synthesis [isa: 8982491]
70221 = Glycogen breakdown (glycogenolysis) [isa: 8982491]
9022692 = Regulation of MECP2 expression and activity [isa: 8986944]
9026527 = Activated NTRK2 signals through PLCG1 [isa: 9006115]
9028731 = Activated NTRK2 signals through FRS2 and FRS3 [isa: 9006115]
9032500 = Activated NTRK2 signals through FYN [isa: 9006115]
9027276 = Erythropoietin activates Phosphoinositide-3-kinase (PI3K) [isa: 9006335]
9027284 = Erythropoietin activates RAS [isa: 9006335]
198323 = AKT phosphorylates targets in the cytosol [isa: 1257604]
5365859 = RA biosynthesis pathway [isa: 5362517]
9009391 = Extra-nuclear estrogen signaling [isa: 8939211]
9024446 = NR1H2 and NR1H3-mediated signaling [isa: 9006931]
1250196 = SHC1 events in ERBB2 signaling [isa: 1227986]
1250342 = PI3K events in ERBB4 signaling [isa: 1236394]
1433559 = Regulation of KIT signaling [isa: 1433557]
187042 = TRKA activation by NGF [isa: 187015]
179812 = GRB2 events in EGFR signaling [isa: 177929]
186763 = Downstream signal transduction [isa: 186797]
190373 = FGFR1c ligand binding and activation [isa: 190242]
194306 = Neurophilin interactions with VEGF and VEGFR [isa: 194138]
165160 = PDE3B signalling [isa: 109703]
6806942 = MET Receptor Activation [isa: 6806834]
8852405 = Signaling by MST1 [isa: 9006934]
2473224 = Antagonism of Activin by Follistatin [isa: 1502540]
2173788 = Downregulation of TGF-beta receptor signaling [isa: 2173789]
201451 = Signaling by BMP [isa: 9006936]
8854214 = TBC/RABGAPs [isa: 9007101]
8876198 = RAB GEFs exchange GTP for GDP on RABs [isa: 9007101]
9634635 = Estrogen-stimulated signaling through PRKCZ [isa: 9009391]
9634638 = Estrogen-dependent nuclear events downstream of ESR-membrane signaling [isa: 9009391]
901032 = ER Quality Control Compartment (ERQC) [isa: 901042]
9013507 = NOTCH3 Activation and Transmission of Signal to the Nucleus [isa: 9012852]
9017802 = Noncanonical activation of NOTCH3 [isa: 9013507]
9013700 = NOTCH4 Activation and Transmission of Signal to the Nucleus [isa: 9013694]
9604323 = Negative regulation of NOTCH4 signaling [isa: 9013694]
9018677 = Biosynthesis of DHA-derived SPMs [isa: 9018678]
9018681 = Biosynthesis of protectins [isa: 9018677]
9018682 = Biosynthesis of maresins [isa: 9018677]
9020265 = Biosynthesis of aspirin-triggered D-series resolvins [isa: 9018677]
9026395 = Biosynthesis of DHA-derived sulfido conjugates [isa: 9018677]
2142700 = Synthesis of Lipoxins (LX) [isa: 9018678]
9018676 = Biosynthesis of D-series resolvins [isa: 9018677]
9018679 = Biosynthesis of EPA-derived SPMs [isa: 9018678]
9018683 = Biosynthesis of DPA-derived SPMs [isa: 9018678]
9027604 = Biosynthesis of electrophilic ω-3 PUFA oxo-derivatives [isa: 9018678]
9018896 = Biosynthesis of E-series 18(S)-resolvins [isa: 9018679]
9023661 = Biosynthesis of E-series 18(R)-resolvins [isa: 9018679]
9027307 = Biosynthesis of maresin-like SPMs [isa: 9018682]
9025094 = Biosynthesis of DPAn-3 SPMs [isa: 9018683]
9025106 = Biosynthesis of DPAn-6 SPMs [isa: 9018683]
9029569 = NR1H3 & NR1H2 regulate gene expression linked to cholesterol transport and efflux [isa: 9024446]
9623433 = NR1H2 & NR1H3 regulate gene expression to control bile acid homeostasis [isa: 9024446]
9026286 = Biosynthesis of DPAn-3-derived protectins and resolvins [isa: 9025094]
9026290 = Biosynthesis of DPAn-3-derived maresins [isa: 9025094]
9026403 = Biosynthesis of DPAn-3-derived 13-series resolvins [isa: 9025094]
9026762 = Biosynthesis of maresin conjugates in tissue regeneration (MCTR) [isa: 9026395]
9026766 = Biosynthesis of protectin and resolvin conjugates in tissue regeneration (PCTR and RCTR) [isa: 9026395]
9032759 = NTRK2 activates RAC1 [isa: 9032500]
9033241 = Peroxisomal protein import [isa: 9609507]
9033807 = ABO blood group biosynthesis [isa: 9033658]
9037629 = Lewis blood group biosynthesis [isa: 9033658]
9034013 = NTF3 activates NTRK3 signaling [isa: 9034015]
9034793 = Activated NTRK3 signals through PLCG1 [isa: 9034015]
9603381 = Activated NTRK3 signals through PI3K [isa: 9034015]
167060 = NGF processing [isa: 9036866]
909733 = Interferon alpha/beta signaling [isa: 913531]
1169408 = ISG15 antiviral mechanism [isa: 1169410]
877312 = Regulation of IFNG signaling [isa: 877300]
912694 = Regulation of IFNA signaling [isa: 909733]
977068 = Termination of O-glycan biosynthesis [isa: 913709]
917977 = Transferrin endocytosis and recycling [isa: 917937]
975956 = Nonsense Mediated Decay (NMD) independent of the Exon Junction Complex (EJC) [isa: 927802]
975957 = Nonsense Mediated Decay (NMD) enhanced by the Exon Junction Complex (EJC) [isa: 927802]
2562578 = TRIF-mediated programmed cell death [isa: 937061]
936964 = Activation of IRF3/IRF7 mediated by TBK1/IKK epsilon [isa: 937061]
937041 = IKK complex recruitment mediated by RIP1 [isa: 937061]
937072 = TRAF6-mediated induction of TAK1 complex within TLR4 complex [isa: 937061]
964739 = N-glycan trimming and elongation in the cis-Golgi [isa: 948021]
975576 = N-glycan antennae elongation in the medial/trans-Golgi [isa: 948021]
1268020 = Mitochondrial protein import [isa: 9609507]
9033500 = TYSND1 cleaves peroxisomal proteins [isa: 9033241]
9603798 = Class I peroxisomal membrane protein import [isa: 9609507]
9609523 = Insertion of tail-anchored proteins into the endoplasmic reticulum membrane [isa: 9609507]
8934903 = Receptor Mediated Mitophagy [isa: 5205647]
9614399 = Regulation of localization of FOXO transcription factors [isa: 9614085]
9615017 = FOXO-mediated transcription of oxidative stress, metabolic and neuronal genes [isa: 9614085]
9617629 = Regulation of FOXO transcriptional activity by acetylation [isa: 9614085]
9617828 = FOXO-mediated transcription of cell cycle genes [isa: 9614085]
964827 = Progressive trimming of alpha-1,2-linked mannose residues from Man9/8/7GlcNAc2 to produce Man5GlcNAc2 [isa: 964739]
5205685 = Pink/Parkin Mediated Mitophagy [isa: 5205647]
9646399 = Aggrephagy [isa: 9663891]
199920 = CREB phosphorylation [isa: 198725 450282]
975144 = IRAK1 recruits IKK complex upon TLR7/8 or 9 stimulation [isa: 975138]
975163 = IRAK2 mediated activation of TAK1 complex upon TLR7/8 or 9 stimulation [isa: 975138]
975110 = TRAF6 mediated IRF7 activation in TLR7/8 or 9 signaling [isa: 975155]
975574 = Reactions specific to the hybrid N-glycan synthesis pathway [isa: 975576]
975577 = N-Glycan antennae elongation [isa: 975576]
975578 = Reactions specific to the complex N-glycan synthesis pathway [isa: 975576]
991365 = Activation of GABAB receptors [isa: 977444]
170670 = Adenylate cyclase inhibitory pathway [isa: 112040 991365]
1236973 = Cross-presentation of particulate exogenous antigens (phagosomes) [isa: 1236975]
983168 = Antigen processing: Ubiquitination & Proteasome degradation [isa: 983169]
983170 = Antigen Presentation: Folding, assembly and peptide loading of class I MHC [isa: 983169]
983189 = Kinesins [isa: 983231]
1169091 = Activation of NF-kappaB in B cells [isa: 1168372]
5690714 = CD22 mediated BCR regulation [isa: 983705]
983695 = Antigen activates B Cell Receptor (BCR) leading to generation of second messengers [isa: 983705]
3295583 = TRP channels [isa: 2672351]
936837 = Ion transport by P-type ATPases [isa: 983712]
997272 = Inhibition  of voltage gated Ca2+ channels via Gbeta/gamma subunits [isa: 991365]
